# Overlapping yet distinct spatiomolecular profiles across mood disorders and biological sex in the human prefrontal cortex

**DOI:** 10.64898/2026.10.22.761067

**Authors:** Jacqueline R. Thompson, Seyun Oh, Sarah E. Maguire, Svitlana V. Bach, Bernard J. Mulvey, Geo Pertea, Madeline R. Abramson, Ryan A. Miller, Sang Ho Kwon, Madhavi Tippani, Madeline R. Valentine, Amy Deep-Soboslay, Yufeng Du, Annie B. Nguyen, Joel E. Kleinman, Kristen R. Maynard, Thomas M. Hyde, Shizhong Han, Leonardo Collado-Torres, Keri Martinowich, Stephanie C. Hicks, Stephanie C. Page

**Affiliations:** Department of Biostatistics, Johns Hopkins Bloomberg School of Public Health, Baltimore, MD, 21205, USA; Lieber Institute for Brain Development, Johns Hopkins Medical Campus, Baltimore, MD, 21205, USA; Department of Psychiatry and Behavioral Sciences, Johns Hopkins School of Medicine, Baltimore, MD, 21205, USA; Biochemistry, Cellular, and Molecular Biology Graduate Program, Johns Hopkins School of Medicine, Baltimore, MD, 21205, USA; Solomon H. Snyder Department of Neuroscience, Johns Hopkins School of Medicine, Baltimore, MD, 21205, USA; Department of Neurology, Johns Hopkins School of Medicine, Baltimore, MD, 21205, USA; Center for Computational Biology, Johns Hopkins University, Baltimore, MD, 21218, USA; Johns Hopkins Kavli Neuroscience Discovery Institute, Baltimore, MD, 21205, USA; Department of Biomedical Engineering, Johns Hopkins University, Baltimore, MD, USA; Malone Center for Engineering in Healthcare, Johns Hopkins University, Baltimore, MD, 21218, USA

**Author notes:** **Correspondence** Keri Martinowich, Stephanie C. Hicks, Stephanie C. Page. These authors jointly supervised the work (Keri Martinowich, Stephanie C. Hicks, Stephanie C. Page).

## Abstract

Major depressive disorder (MDD) and bipolar disorder (BD) exhibit both overlapping and distinct clinical features, including sex differences in prevalence and symptomatology, but how these similarities and differences are reflected at the molecular and cellular level in the human brain remains poorly understood. Here, we applied spatial transcriptomics to the human dorsolateral prefrontal cortex from individuals with MDD, BD, and neurotypical controls to resolve molecular programs associated with diagnosis and biological sex. Our analysis revealed sex- and diagnosis-specific differences across white matter, the neurovascular unit (NVU), and neuronal compartments. Within white matter, we identified female-specific survival-response programs shared in both MDD and BD, and divergent myelin-related gene expression patterns across disorders (*CLDN11*, *CNDP1*). Within the NVU, transcriptional evidence of impaired vascular integrity was accompanied by astrocytic response profiles in females with MDD and males with BD, with inflammatory responses further evident in females with MDD (*IFITM3*, *CEBPD*, *IL1R1*). In donors with BD of both sexes and in females with MDD, inhibitory neurons exhibited impaired neuropeptide expression (*SST*, *CORT*, *CRH*). Integration with donor genotype information highlighted that genetic risk for mood disorders influences *MAPK3* expression, implicating altered ERK signaling, which was further supported by disorder-associated changes in Ras/MEK/ERK pathway components in neurons (*MAPK3, DUSP6*, *RASD1*, *ELK1*). Together, these findings reveal coordinated, cell type- and spatial domain-specific molecular programs that distinguish shared and disorder-specific transcriptomic changes across mood disorders in a sex-dependent manner.

## 1 Introduction

Major depressive disorder (MDD) and bipolar disorder (BD) are common, complex neuropsychiatric conditions that carry major economic and medical burdens (Cui et al. 2024; Levenberg and Cordner 2022). At the genetic, molecular, cellular, circuit, and clinical levels, aspects of MDD and BD overlap while others are unique. Both MDD and BD are characterized by prolonged depressive episodes encompassing anhedonia, changes in appetite and sleep, and lack of motivation, and BD is further characterized by episodes of mania, disordered thinking, and, in some cases, psychosis (Levenberg and Cordner 2022; Cui et al. 2024; Marx et al. 2023). MDD is twice as common in females as males, with females exhibiting longer depressive episodes of greater severity, and antidepressant efficacy varying by sex (LeGates et al. 2019; Carvalho Silva et al. 2023; Y. Zhang et al. 2024). Although BD prevalence is largely similar across biological sex, recent studies suggest that clinical characteristics of BD may differ, including different comorbidities, females exhibiting earlier age onset and longer duration of untreated illness, and males presenting with more mixed features, especially during depressive episodes (Putz et al. 2025; Miola et al. 2025; Menculini et al. 2022; Loftus et al. 2020). Thus, a better understanding of the biological mechanisms mediating these sex-associated differences in both MDD and BD is critical to improving patient outcomes.

Although specific causes of MDD and BD are unclear, both genetic and environmental factors interact with biological sex to contribute to risk. Genome-wide association studies (GWAS) have identified shared genetic risk across psychiatric disorders (Grotzinger et al. 2026), with associations between common genetic variants and increased risk for MDD (Adams et al. 2025) and BD (O’Connell et al. 2025). However, other risk factors influence mood disorder incidence, including family history, socioeconomic factors, age, and stress (LeMoult et al. 2020; Agerbo et al. 2021; Freeman et al. 2025; Jelaga et al. 2025). Stress, herein defined as robust or sustained fight-or-flight response resulting from adverse life experience, is a risk factor for depression (Hammen 2005; M.-H. Han and Nestler 2017), and has been implicated in the neurobiological mechanisms of both MDD and BD (Meyer-Lindenberg and Tost 2012; McEwen and Akil 2020). Recent research emphasizes the gene-by-environment nature of the intersecting influences of genetic risk, stress, and biological sex in psychiatric and mood disorders and implicates specific cell types (inhibitory neurons, vascular endothelial cells) in potential causal mechanisms including neuronal excitability, dopamine signaling, and inflammatory and immune response (Nievergelt et al. 2019; Blokland et al. 2022; Silveira et al. 2023; Strom et al. 2026).

The dorsolateral prefrontal cortex (dlPFC) is involved in cognitive control, executive functioning, and emotional regulation, all of which are disrupted in MDD and BD (Price and Drevets 2010; Friedman and Robbins 2022). Assessments of functional connectivity implicated the dlPFC in BD, both within and across mood states (Townsend et al. 2010; Pomarol-Clotet et al. 2015; Martino et al. 2016), and in MDD, where dlPFC activity is associated with antidepressant response and depression symptoms (Chin Fatt et al. 2021; Taylor et al. 2022; Zheng et al. 2023). Accordingly, the dlPFC is a principal target for noninvasive brain stimulation therapies, including transcranial magnetic stimulation, which has demonstrated therapeutic efficacy for depression in MDD and BD (Sabé et al. 2024; Kishi et al. 2024; Kazemi et al. 2018; Miuli et al. 2021; Elias et al. 2021). The six-layered structure of the dlPFC influences its circuit-level function, with superficial layer neurons integrating largely cross-cortical input and deep layer neurons projecting to subcortical target regions. Although the structural organization of the dlPFC is linked to its function, most transcriptomics studies of MDD and BD have used dissociated tissue samples in which laminar organization and spatial context are not retained. Additionally, some studies grouped males and females together in the primary analyses, potentially obscuring sex-specific transcriptional changes. Targeted transcript quantification studies found reductions in neuropeptide inhibitory neuron markers in the dlPFC both MDD and BD (Fung et al. 2014; Chung et al. 2018; Dienel et al. 2023; Okuda et al. 2024). Bulk RNA-seq studies revealed associations between BD diagnosis and oligodendrocyte, microglia, and astrocyte function when male and female donors were analyzed together (Wei et al. 2024; Xia et al. 2024), while sex-stratified analysis suggested greater BD-associated changes in these cell types in males (Xia et al. 2024). In MDD, a large bulk RNA-seq study spanning two cortical and two amygdala subregions identified substantially fewer differences in the dlPFC compared to the dorsal anterior cingulate or the medial amygdala (Jaffe et al. 2022). Other bulk RNA-seq studies of the dlPFC have reported sex-specific transcriptional changes (Labonté et al. 2017; Mansouri et al. 2023), although re-analysis accounting for multiple comparisons only identified significant changes in females but not males (de Souza et al. 2024). Single-nucleus RNA-seq (snRNA-seq) studies of the dlPFC identified sex-specific alterations in glial cell types and inhibitory neuron populations in MDD (Nagy et al. 2020; Maitra et al. 2023), while snATAC-seq in the same cohort identified changes in chromatin accessibility in microglia, astrocytes, and oligodendrocytes, even when male and female donors were analyzed together (Chawla et al. 2025). Together, these findings highlight the importance of biological sex and cellular context in mood disorders, but leave unresolved how these molecular alterations are spatially organized across the cortical architecture.

To investigate how gene expression changes associated with mood disorders are distributed across the laminar organization of the human dlPFC, we used the Visium platform to generate spatially-resolved transcriptomic (SRT) data from donors with MDD, BD, and neurotypical controls, with approximately equal numbers of male and female donors in each diagnostic group. This approach preserved cortical architecture and enabled the capture of extranuclear transcripts and RNA from small, non-neuronal cell populations, including vascular cells, that are difficult to capture using droplet-based snRNA-seq. We hypothesized that biological sex influences the transcriptional mechanisms underlying mood disorders, and therefore examined diagnosis-associated gene expression separately in males and females. Our findings implicate alterations in inhibitory neurons and neurovascular cells across disorders, with prominent inflammatory responses in females with MDD and distinct oligodendrocyte-associated transcriptional programs in females with MDD and BD. We further integrated donor genotype data with spatial gene expression to identify genetic variants associated with expression in specific cellular contexts and examine their relationship to genetic risk for mood disorders. Together, these findings provide insight into the sex- and diagnosis-specific molecular architecture of mood disorders within the human dlPFC.

## 2 Results

### 2.1 Spatially-resolved transcriptomic data collection and domain annotation

Our cohort contained 119 postmortem human donors spanning major depressive disorder (MDD; N=39), bipolar disorder (BD; N=40) and neurotypical control (NTC; N=40), with each diagnosis group evenly split by biological sex and balanced for age (**Figure 1A-B**, **Table 1**, **ST1**, **Methods 4.1**). While genetic risk for the individual donors (aggregated across the genome) was evaluated by calculating polygenic risk scores (PRS) based on GWAS summary statistics for MDD and BD (Adams et al. 2025; O’Connell et al. 2025), we did not use PRS as a factor for donor selection. In our cohort, PRS for BD was predictive of BD diagnosis, while PRS for MDD did not predict MDD diagnosis (**SF1**, **Methods 4.1.3**). Given that the contribution of genetic risk to these disorders is relatively small, these findings are not unexpected, but do suggest some genetic liability for BD in our cohort.

**Figure 1.**
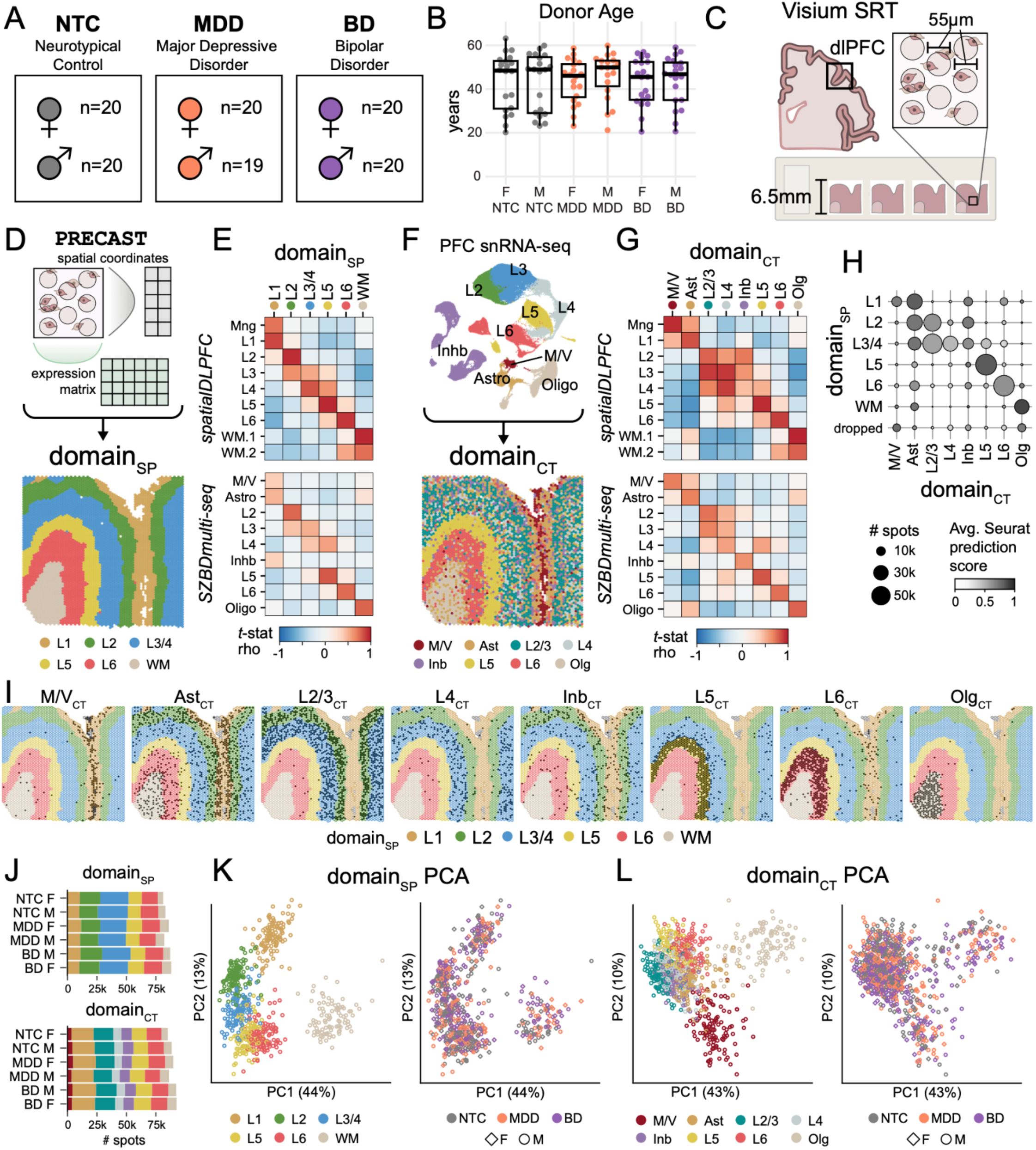
Overview of donor cohort and identification of spatial domains across mood disorders in the human dlPFC. (**A**) Schematic of the donor cohort across neurotypical control (NTC), Major Depressive Disorder (MDD), and Bipolar disorder (BD). (**B**) Distribution of donor age at time of death, by diagnosis and biological sex (female, F; male, M). (**C**) Schematic of spatially-resolved transcriptomics (SRT) data generation in postmortem human dorsolateral prefrontal cortex (dlPFC) tissue using 10x Genomics Visium. (**D**) Annotating spot-level spatial domains using `PRECAST` in an example tissue section (Br6432) reflecting traditional dlPFC cytoarchitectural layers (domain annotation spatial clustering, domain_SP_). Annotations included layers (L) L1, L2, L3/4, L5, L6, white matter (WM) domains. (**E**) Spatial registration of domain_SP_ (this study, *x*-axis) with previous dlPFC SRT data (*y*-axis, top (Huuki-Myers et al. 2024)) and PFC snRNA-seq data (*y*-axis, bottom (Ruzicka et al. 2024)). (**F**) Annotating spot-level spatial domains using label transfer in an example tissue section (Br6432) using snRNA-seq (Ruzicka et al. 2024) cell type expression profiles (domain annotation cell type-driven, domain_CT_). Annotations included included microglia and vascular (M/V), astrocytes (Ast), Layers (L) L2/3, L4, L5, L6, inhibitory neurons (Inb) and oligodendrocytes (Olg). (**G**) Spatial registration of domain_CT_ (this study, *x*-axis) with previous dlPFC SRT data (*y*-axis, top (Huuki-Myers et al. 2024)) and PFC snRNA-seq data (*y*-axis, bottom (Ruzicka et al. 2024)). (**H**) Dot plot showing the number of spots in agreement using the two annotation strategies (domains_SP_ and domains_CT_). Size of dots represents the number. Gray scale represents the average Seurat prediction score of domains_CT_ spots from domains_SP_ spots. (**I**) Spot plots illustrating agreement between domains_SP_ (colored spot outline) and domains_CT_ (spot fill in black). (**J**) Stacked bar plots of the number of spots in the pseudobulk samples annotated by domain_SP_ (top) and domain_CT_ (bottom) stratified by diagnosis and biological sex. (**K**) Principal component analysis (PCA) of pseudobulk domain_SP_, colored by domain (left) or diagnosis (right, shape indicates biological sex). (**L**) Similar to (**K**), but using pseudobulk samples annotated by domain_CT_.

**Table 1.** Cohort demographics summary. Summarized demographic information (number of donors or mean ± SD) for diagnosis-by-sex (dx-sex) groups. Age: years at time of death; BMI: body mass index in kg/m^2^; RIN: RNA integrity number; Nicotine use: as reported by next-of-kin, number of donors; Death by suicide: number of donors. Group differences were assessed via ANOVA (`∼ diagnosis*sex`) for continuous variables (age, BMI, RIN; *F*-statistic reported) or chi-squared tests (3 separate tests for diagnosis, sex, and dx-sex group) for categorical variables (nicotine use, death by suicide; Χ^2^ reported). ^a^ *p* < 0.05

|  | Diagnosis-by-sex group |  |  |  |  |  | Group differences test statistic |  |  |
| --- | --- | --- | --- | --- | --- | --- | --- | --- | --- |
|  | NTC F | NTC M | MDD F | MDD M | BD F | BD M | Diagnosis | Sex | Dx-sex group |
| Donors | 20 | 20 | 20 | 19 | 20 | 20 | - | - | - |
| Age | 43.6 ± 13.0 | 43.1 ± 13.2 | 43.6 ± 9.9 | 46.1 ± 10.6 | 43.1 ± 10.7 | 44.0 ± 11.2 | 0.199 | 0.197 | 0.167 |
| BMI | 30.2 ± 9.6 | 31.5 ± 9.9 | 32.2 ± 9.7 | 31.3 ± 6.6 | 31.3 ± 8.7 | 29.8 ± 6.4 | 0.201 | 0.0478 | 0.276 |
| RIN | 7.42 ± 0.93 | 7.93 ± 0.78 | 7.88 ± 0.66 | 8.02 ± 0.88 | 8.03 ± 0.61 | 7.97 ± 0.68 | 2.03 | 2.01 | 1.40 |
| Nicotine use | 3 | 3 | 13 | 12 | 16 | 12 | 29.1<sup>a</sup> | 0.413 | 30.7<sup>a</sup> |
| Death by suicide | 0 | 0 | 3 | 8 | 3 | 3 | 12.9<sup>a</sup> | 1.18 | 18.7<sup>a</sup> |

A single tissue section from the dlPFC was collected from each donor to generate spatially-resolved transcriptomic (SRT) profiles using the Visium platform (**Methods 4.2**). Visium SRT incorporates spatial barcodes for approximately 5,000 55 μm diameter spots across a tissue section, linking each mRNA transcript to spatial coordinates aligned to the corresponding H&E image (**Figure 1C**). Following quality control (**SF2**, **Methods 4.3**), we annotated SRT spots using two complementary strategies. First, using spatially-aware clustering (W. Liu et al. 2023), we generated spatial domains (domains_SP_) that recapitulated dlPFC cytoarchitecture, including layers (L) L1, L2, L3/4, L5, L6, and white matter (WM) (**Figure 1D**, **SF3**, **Methods 4.4.1**). While comparison of domains_SP_ to published dlPFC spatial domain annotations (spatialDLPFC (Huuki-Myers et al. 2024)) confirmed that they reflected canonical laminar structure, comparison to published single-nucleus RNA-sequencing (snRNA-seq) from the PFC (SZBDMulti-seq (Ruzicka et al. 2024)) revealed limited representation of cell types that transcend laminar boundaries, particularly glia and inhibitory neurons (**Figure 1E**). To better capture these cellular populations, we generated a second set of domain annotations by projecting the SZBDMulti-seq snRNA-seq cell type labels into our SRT dataset (domains_CT_; **Figure 1F**, **SF4**, **Methods 4.4.2**). These domains replicated the laminar organization of the dlPFC (L2/3, L4, L5, L6, Olg), but also captured domains representing microglia and vascular cells, (M/V_CT_), astrocytes (Ast_CT_), and inhibitory neurons (Inb_CT_) (**Figure 1G**, **Methods 4.4.3**). The domain_CT_ annotations replicating laminar organization were highly consistent with domain_SP_, and the translaminar annotations enabled the separation of cytoarchitectural layers into known cellular components (e.g. L1_SP_ split into M/V_CT_, Ast_CT_, and Inb_CT_) (**Figure 1H-I**, **SF5A**).

We verified that domain abundance and composition (via marker gene expression) were consistent across diagnosis and sex for both domain_SP_ and domain_CT_ strategies (**Figure 1J-L**, **SF3**, **SF4**, **SF5B**). The complementary information provided by the two annotation strategies allowed us to examine mood disorder-associated gene expression in the context of well-defined dlPFC lamina as well as in cell types that are distributed across the canonical laminar structure.

### 2.2 Differential expression across mood disorders converges on coordinated molecular programs

To identify sex-specific gene expression changes associated with mood disorders, we compared NTC, MDD, and BD donors separately within each sex, generating six diagnosis-by-sex contrast groups (F_NTC.MDD, F_NTC.BD, F_MDD.BD, M_NTC.MDD, M_NTC.BD, M_MDD.BD). Differential expression (DE) analyses were performed on pseudobulked spots from both domain annotation strategies (domain_SP_ and domain_CT_) using two complementary linear models (Ritchie et al. 2015). First, to identify gene expression changes at the tissue level, the whole-tissue linear model tested for diagnosis-by-sex differences across the entire tissue section while adjusting for differences in expression due to domain annotations (**Figure 2A-B**). Second, to localize gene expression changes to specific domains, the domain-restricted linear model evaluated diagnosis-by-sex differences within individual spatial domains (**Figure 2C-D**). Both models controlled for donor age, spots per pseudobulk sample, and technical variation (**SF6**; **Methods 4.5**).

**Figure 2.**
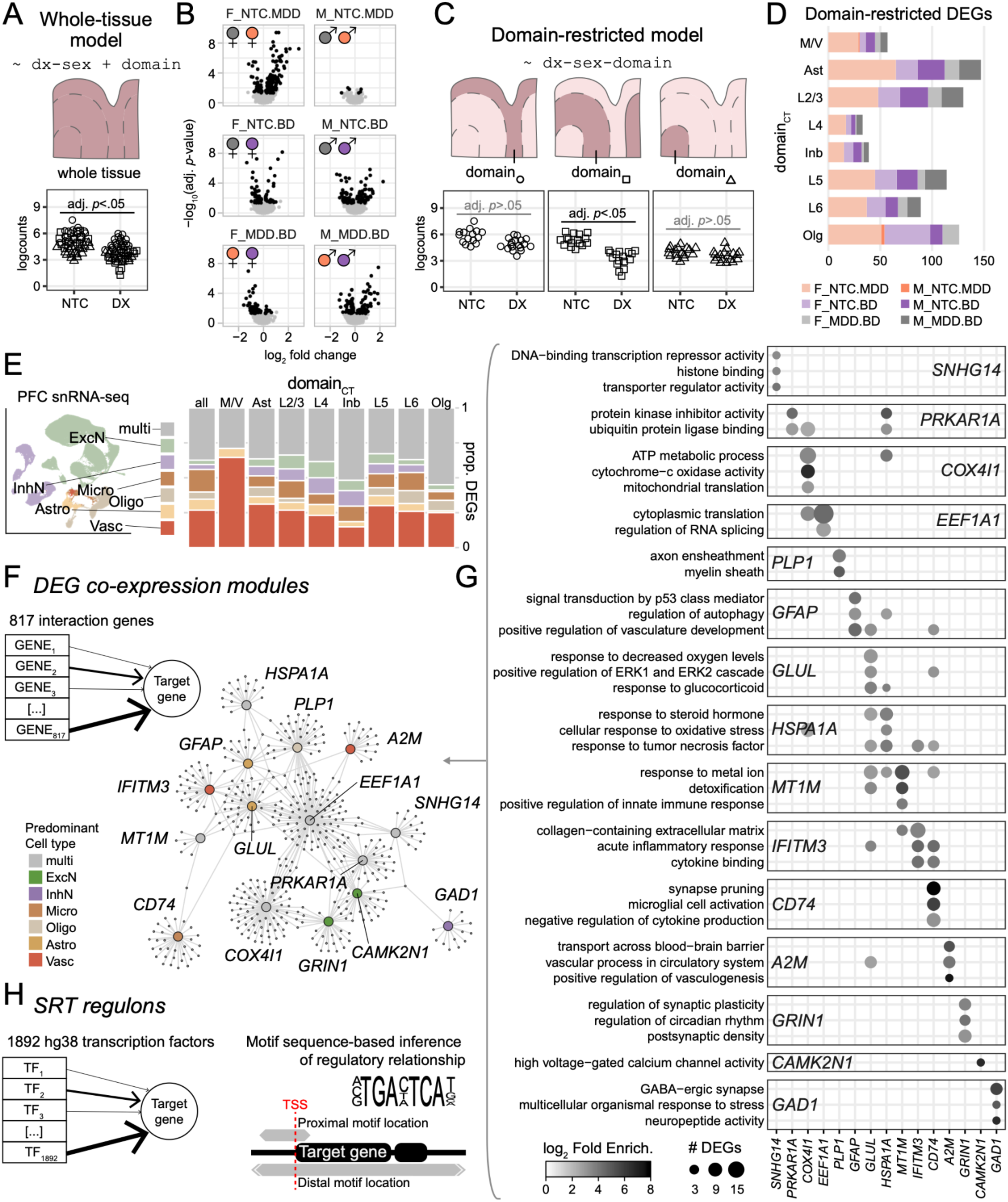
Identification and characterization of gene expression changes across mood disorders and spatial domains in the human dlPFC. (**A**) Top: Schematic of the whole-tissue model, which identifies changes in gene expression at the tissue level (using all spots across the tissue) associated with donor-level diagnosis and biological sex variables (dx-sex) while adjusting for domain composition. Bottom: Schematic of an example differentially expressed gene showing the distribution of log_2_-normalized gene expression (logcounts) using pseudobulk samples illustrating a higher logcounts for neurotypical control (NTC) compared to diagnosis (dx) groups. (**B**) Volcano plots showing the log_2_-fold change of differentially expressed genes using the domain_CT_ whole-tissue model. Within each sex, we compared neurotypical controls (NTC), major depressive disorder (MDD) and bipolar disorder (BD), generating six dx-sex contrasts (F_NTC.MDD, F_NTC.BD, F_MDD.BD, M_NTC.MDD, M_NTC.BD, M_MDD.BD). Colors correspond to diagnostic groups including NTC (grey), MDD (orange), and purple (BD). (**C**) Top: Schematic of the domain-restricted model, which identifies changes in gene expression restricted to specific domains (3 domains in the schematic as an example) associated with donor-level dx and sex variables. Bottom: Similar to (**A**), but shape of points corresponds to using pseudobulk samples restricted to specific domains for each differential expression test. (**D**) Bar plot of the number of differentially expressed genes (DEGs) tested using the domain-restricted model (omnibus *F*-test adj. *p*<.05 and *t*-test adj. *p*<.05) for each domain_CT_, colored by the six contrasts or comparison groups. (**E**) Stacked bar plot of the predominant cell type (evaluated by *MeanRatio* analysis of PFC snRNA-seq data (Ruzicka et al. 2024)) expressing the DEGs identified in whole-tissue (“all”) and domain-restricted models. (**F**) Inset diagram of *GRNBOOST2* pipeline utilizing 513,200 spots to generate co-expression modules for 817 genes with significant *F*-omnibus tests (adj. *p*<.05). The network plot highlights the 15 co-expression modules that encompasses 446 DEGs. Color represents the predominant cell type of origin for each DEG co-expression module. (**G**) Top results of overrepresentation analysis of GO terms (adjusted p < 0.05) for each DEG co-expression module. (**H**) Schematic of regulon analysis which aims to identify transcription factor networks (1892 human transcription factors) that could underlie the patterns of SRT expression.

We identified 503 unique differentially expressed genes (DEGs; omnibus *F*-test adj. *p* < 0.05 and moderated *t*-test adj. *p* < 0.05) across statistical models and spatial domain annotation strategies (**Methods 4.5.3**, **ST2**, **ST3**). These included 215 DEGs in females with MDD, 168 DEGs in females with BD, and 134 DEGs in males with BD, each compared to sex-matched controls (**SF7**, **SF8**). In contrast, only 13 unique DEGs were identified in any comparison involving males with MDD, with only two genes reaching significance criteria in both domain_SP_ and domain_CT_ whole-tissue models (*EDNRB* decreased vs NTC, *ALDOA* increased vs NTC). Despite the limited number of DEGs, decreased expression of *EDNRB* was consistent with a previous report of reduced *EDNRB* expression in astrocytes from males with MDD (Maitra et al. 2023). These findings reveal pronounced sex-dependent differences in MDD-associated gene expression in the dlPFC. DE patterns were highly concordant across the two spatial domain annotation strategies, and most domain-restricted DEGs were also detected in the whole-tissue model (**SF9**). The principal exception was the WM_SP_ and Olg_CT_ domains, where approximately 40 female-specific DEGs were detected exclusively in domain-restricted analyses, demonstrating the value of examining sex- and diagnosis-associated changes within specific spatial compartments.

To provide biological context for these DEGs, we implemented three complementary analytical approaches. First, we again leveraged the SZBDMulti-seq dlPFC snRNA-seq dataset (Ruzicka et al. 2024) (**Figure 1F**) to infer the predominant cell types expressing each DEG (**SF10**, **Methods 4.6**). Many DEGs were predominantly expressed in NVU cell populations and in inhibitory neurons (**Figure 2E**, **SF10E**). Second, we used spot-level co-expression analysis (Moerman et al. 2019) to organize genes into coordinated transcriptional modules, which were then filtered to identify diagnosis-by-sex DEGs representing shared biological programs (**Figure 2F, SF11**, **SF12**, **Methods 4.7.1**). This approach assigned 446 of the 503 DEGs (88.7%) to 15 co-expression modules that were associated with biological functions of all major dlPFC cell types: excitatory and inhibitory neurons, astrocytes, oligodendrocytes, microglia, and vascular cell types (**Figure 2G**, **ST4**, **SF13**, **SF14**). Third, we performed regulon analysis (Aibar et al. 2017), integrating spot-level co-expression with genomic motif enrichment to identify candidate transcriptional regulatory networks (**Figure 2H, Methods 4.7.2**). We focused on regulon component genes or transcription factors that were themselves identified as diagnosis-by-sex DEGs. Prior to constructing these regulatory networks, we excluded a small subset of donors exhibiting atypical activity-dependent transcriptional signatures unrelated to diagnosis or sex, likely reflecting peri-mortem neuronal activation (**SF15**). Regulon analysis identified transcription factor networks corresponding to oligodendrocyte identity, extracellular stimulus response programs, and inhibitory neuron identity, several of which correlated strongly with the DEG co-expression modules (**SF16**, **ST5**).

These complementary analyses linked diagnosis- and sex-specific gene expression to specific cell populations, biological processes, and transcriptional regulatory networks. Collectively, evidence from these analyses converged on three major biological programs: oligodendrocyte and white matter alterations, neurovascular and inflammatory responses, and neuronal dysregulation.

### 2.3 Female-specific white matter transcriptional programs distinguish MDD and BD

Across both spatial domain annotation strategies (domain_SP_ and domain_CT_), we identified pronounced transcriptional changes in white matter-associated domains, which were largely restricted to females (**Figure 2D**, **SF9**). In the Olg_CT_ and WM_SP_ domains, females with MDD or BD exhibited increased expression of genes including *BCL6*, *HSPA1B*, *ANP32B, TPT1*, and *UBA52*, consistent with anti-apoptotic and pro-survival inflammatory responses (Toney et al. 2000; Dent et al. 1997; Borges et al. 2013; Dokladny et al. 2010; Chemnitz et al. 2019; Miao et al. 2024; H. Ji et al. 2025) (**Figure 3A-B, SF17A-B**). Similar white matter-associated transcriptional changes in females with MDD and BD were further captured by the *EEF1A1-*module, which included *TPT1*, *UBA52*, *ANP32B*, and other DEGs upregulated in WM_SP_/Olg_CT_ (**SF14**, **SF17C**). *EEF1A1-*module DEGs were enriched for cytoplasmic translation and regulation of RNA splicing (**Figure 2G**), implicating these processes as shared features of the white matter-localized transcriptional response across mood disorders in females. In addition to oligodendrocytes, cortical white matter contains astrocytes, microglia and vascular cell types, which may also contribute to the observed transcriptional alterations associated with survival and growth.

**Figure 3.**
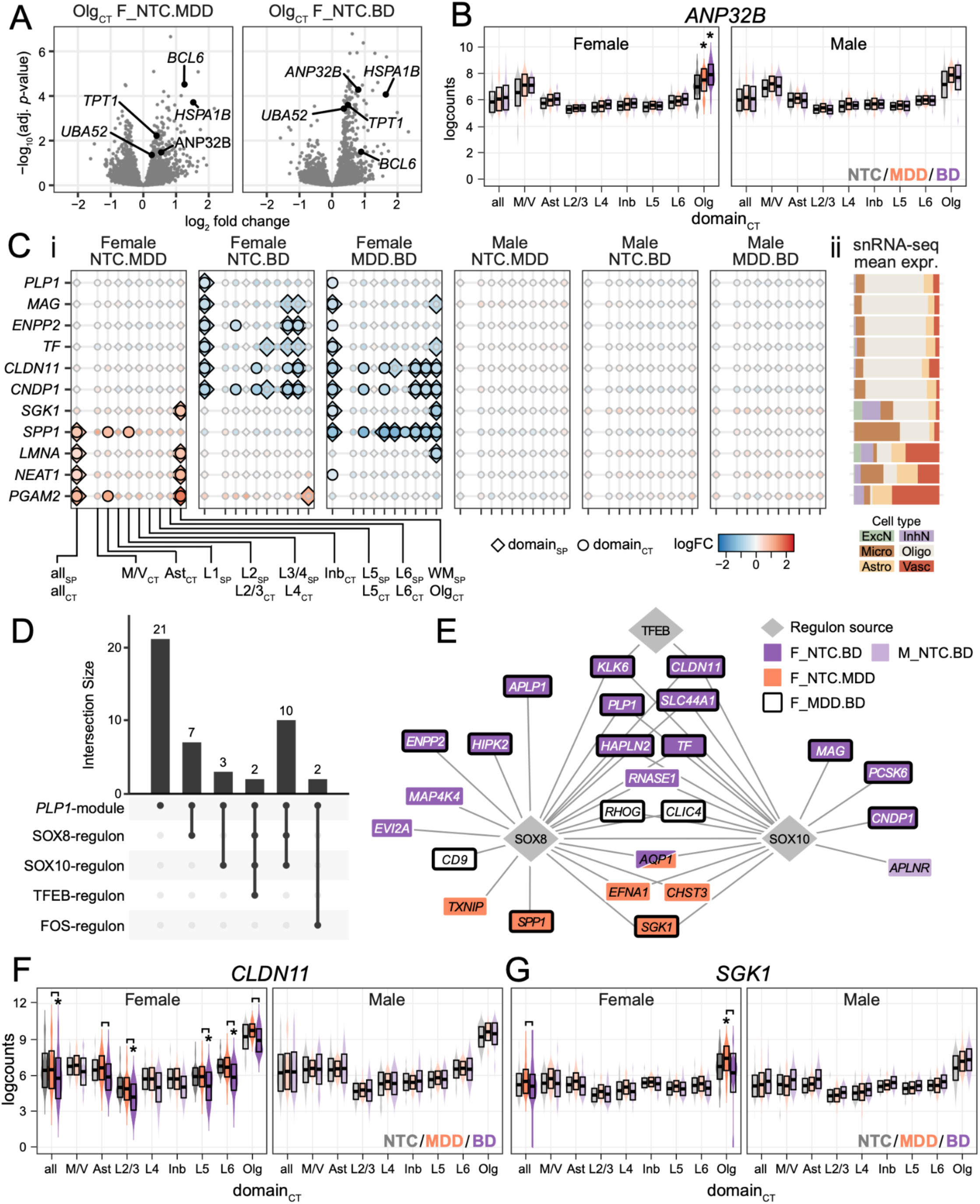
Female-specific transcriptomic changes in white matter across diagnosis. (**A**) Olg_CT_ domain volcano plots for F_NTC.MDD (left) and F_NTC.BPD (right). Highlighted genes indicate DEGs identified with domain-restricted Olg_CT_ and WM_SP_ DE models that were not found in whole-tissue analysis. (**B**) Violin and box plots of *ANP32B* expression for domain_CT_ pseudobulk data with logcounts on *y*-axis and domains on *x*-axis (‘all’ on *x*-axis represents whole-tissue model results from all pseudobulk samples). Statistically significant *t*-test (adj. *p*<.05) versus sex-matched control group indicated by asterisk. Colors correspond to different groups and group comparisons that are statistically significant are in a dark color. Crossbar indicates mean ± standard deviation. (**C**) (**i**) Dotplot of select *PLP1-*module genes across domain annotations (x-axis, ‘all’ represents whole-tissue model results) where large size indicates DEG status, shape indicates domain annotation strategy, and color reflects log_2_ fold change (logFC). Fold change is calculated for each diagnosis-by-sex group (facet) with the reference group listed first and the comparison group listed second (i.e. NTC.MDD with logFC < 0 reflects expression where MDD is less than NTC). (**ii**) Mean expression of y-axis *PLP1*-module genes for each broad cell type from snRNA-seq, scaled to fill x-axis. **(D)** Upset plot indicating overlap of *PLP1-*module DEGs with any regulon (empty intersections removed). (**E**) Network plot of all SOX8-regulon and SOX10-regulon targets that are DEGs, and any additional regulons the target DEGs were attributed to. Shape identifies regulon source transcription factor (diamond) from target genes (rectangle). Fill color and border color reflects diagnosis-by-sex significance. *AQP1* is the only gene significant in both F_NTC.MDD and F_NTC.BD, reflected by split fill color. (**F-G**) Violin plot of *CLDN11* and *SGK1* expression, respectively, in domains_CT_. Same as (**B**), with additional bar symbols indicating statistical significance (adj. *p*<.05) between MDD and BD.

While the *EEF1A1-*module captured white matter transcriptional responses in both MDD and BD females, the *PLP1-*module revealed distinct oligodendrocyte-associated transcriptional programs between the disorders. The *PLP1*-module contained 45 DEGs that were enriched for genes related to oligodendrocyte biology and myelination (**Figure 2G**) and were almost exclusively differentially expressed in females (**SF14B**). *PLP1*-module DEGs reflected a significant reduction of canonical myelination gene expression (*PLP1, MAG, TF*, *ENPP2, CLDN11* and *CNDP1*) in females with BD when compared to either NTC or MDD females (Cheli et al. 2023; Quarles 2007; Tanaka et al. 2009; Maheras et al. 2018; Marchis et al. 2000; Savaskan et al. 2007) (**Figure 3C**). Although previous studies demonstrated changes in myelin-related gene expression in MDD and BD (Rajkowska et al. 2015; Tham et al. 2011; Abraham et al. 2022; Ali et al. 2024; Zhou et al. 2024), our findings identify a pronounced, female-specific reduction in myelin-associated transcription in BD.

Compared to NTC or BD females, females with MDD exhibited increased expression of a distinct set of *PLP1*-module genes, including the WM_SP_/Olg_CT_-restricted genes *LMNA* and *NEAT1*, both of which are linked to the transcriptional maintenance of the myelinating state in oligodendrocytes (Katsel et al. 2019; Pruvost et al. 2023) (**Figure 3C**). *SGK1*, a central mediator of cellular responses to stress that is elevated in oligodendrocytes following stress and is linked to demyelination (Lang et al. 2018; Hinds et al. 2017; Z. Jiang et al. 2025; Deng et al. 2026), was upregulated in WM_SP_ and Olg_CT_ in females with MDD relative to sex-matched NTC or BD. These distinct transcriptional signatures in white matter-associated domains suggest diagnosis-specific alterations in stress-responsive and myelin-related oligodendrocyte programs in MDD and BD females.

To better understand the potential regulatory mechanisms associated with these distinct transcriptional responses, we examined how the diagnosis-associated *PLP1*-module DEGs overlapped with the transcription factor regulons detected in our SRT data (**Figure 2H**). While two *PLP1*-module DEGs (*NEAT1* and *PGAM2*) were present in the FOS-regulon, which mediates varied extracellular responses in oligodendrocytes (Cruz-Mendoza et al. 2022), approximately half of the *PLP1*-module DEGs were also represented in regulons associated with canonical oligodendrocyte identity transcription factors SOX8 and SOX10 (**Figure 3D**). Genes within the highly similar SOX8- and SOX10-regulons encompassed BD-associated DEGs (e.g. *CLDN11*) and MDD-associated DEGs (e.g. *SGK1*) (**Figure 3E**), linking both diagnosis-specific transcriptional patterns within the *PLP1*-module to canonical oligodendrocyte regulatory networks. Importantly, *CLDN11* and *SGK1* were among the key regulatory targets that exhibited significantly different WM_SP_ and Olg_CT_ expression in females with MDD compared to females with BD (**Figure 3E-G, SF17D-E**). Together, these findings suggest that differential engagement of SOX8/SOX10 regulatory networks may contribute to the distinct oligodendrocyte transcriptional programs in MDD and BD females, despite shared evidence of pro-surivival signaling.

### 2.4 Glial activation and vascular dysfunction in the neurovascular unit (NVU)

Our integrative analysis identified disorder-associated changes in dlPFC gene expression associated with the neurovascular unit (NVU), a coordinated network of astrocytes, microglia and vascular cells that is essential for neuronal nutrient uptake and blood-brain barrier (BBB) maintenance that has been implicated in stress and depression (Kaplan et al. 2020; Dudek et al. 2021; Welcome 2020). In our study, 40.7% of all DEGs found throughout dlPFC spatial domains were predominantly expressed in NVU cell types (**Figure 2E**, **SF10E**). Multiple DEG co-expression modules were enriched for genes involved in blood-brain barrier (BBB) function and neuronal homeostasis (**Figure 2G**, **Figure 4A**). For example, the *A2M*-module comprised disorder-associated DEGs that are highly expressed in vascular cells and microglia and were broadly decreased in females with MDD and in donors with BD of both sexes (**Figure 4B**). *A2M*-module DEGs included genes involved in vascular integrity (*TNFSF10, RAMP2,* and *LMO2* (Cartland et al. 2024; Zauli and Secchiero 2006; Xian et al. 2017; Koyama et al. 2013; Gratzinger et al. 2009; Yamada et al. 2022)), BBB transport (*ABCG2* (Schulz et al. 2023)), and astrocyte-dependent glutamate homeostasis (*SLC38A5* (Hamdani et al. 2012; Cubelos et al. 2005)). *A2M,* which encodes a protease with a protective role in inflammation (Rehman et al. 2013; Vandooren and Itoh 2021; Cucullo et al. 2003), was reduced in females with MDD and in donors of both sexes with BD. These findings are consistent with prior research identifying associations between reduced BBB integrity and psychiatric and mood disorders (Konstantinou and Warsh 2026; Ferraro et al. 2026). Given these transcriptional changes implicating BBB function, we further examined other NVU-associated DEG modules to understand whether other aspects of this crucial system were altered across mood disorders.

**Figure 4.**
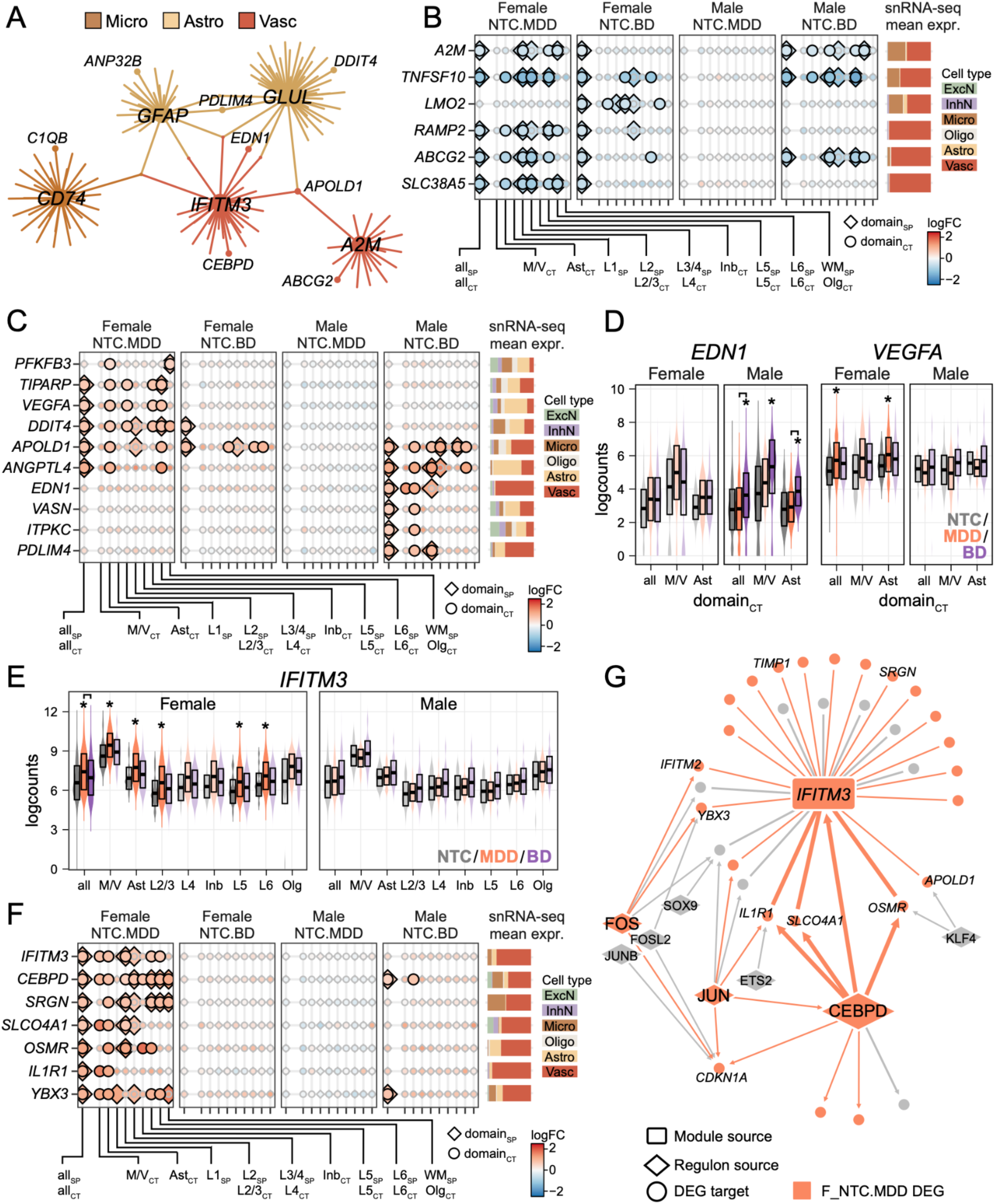
Transcriptomic changes indicate impairments in blood-brain-barrier integrity in mood disorders and sex- and diagnosis-specific changes in the neurovascular unit. (**A**) Network plot showing overlaps of neurovascular unit (NVU) modules examined, with key genes labeled. Node and edge color indicate which cell types of the NVU are best represented in each DEG module. (**B**) (**i**) Dotplot of select *A2M-*module genes across domain annotations (x-axis, ‘all’ represents whole-tissue model results) where large size indicates DEG status, shape indicates domain annotation strategy, and color reflects log_2_ fold change (logFC). Fold change is calculated for each diagnosis-by-sex group, with comparisons to NTC shown in facets. (**ii**) Mean expression of y-axis *A2M*-module genes for each broad cell type from snRNA-seq, scaled to fill x-axis. (**C**) Dotplot for key *GLUL*-module DEGs, see legend of (B) for details. (**D**) Violin and boxplots of two *GLUL*-module DEGs that have similar functions but are significant in different groups. Domain_CT_ DE results are shown for the whole-tissue model (‘all’) and the domain-restricted model for the NVU domains. Statistical significance (*t*-test (adj. *p*<.05) versus sex-matched control group indicated by asterisk. Bar symbols indicating statistical significance (adj. *p*<.05) between MDD and BD. (**E**) Violin and boxplot of *IFITM3* expression in domain_CT_. Statistical significance (*t*-test (adj. *p*<.05) versus sex-matched control group indicated by asterisk. Bar symbols indicating statistical significance (adj. *p*<.05) between MDD and BD. (**F**) Dotplot for key *IFITM3*-module DEGs, see legend of (B) for details. (**G**) Network plot of all *IFITM3*-module DEGs and their predicted regulons (if any), plus all components of the CEBPD-regulon. Circle nodes represent DEG components with key genes labeled with text, with a rectangular node for *IFITM3* indicating its status as source of the IFITM3-module. Diamond nodes represent regulon transcription factor sources and regulon edges include arrows to reflect directed regulatory relationships. Thick edges indicate overlaps of *IFITM3*-module and CEBPD-regulon. Nodes are colored orange to indicate F_NTC.MDD significance, and edges are colored orange when connecting two F_NTC.MDD DEGs.

Co-expression analysis identified two modules defined by canonical astrocyte markers (*GFAP*- and *GLUL*-modules) that exhibited distinct spatial distributions and were associated with different aspects of astrocyte function (**Figure 2G, SF12**). *GFAP*-module DEGs were prevalent in WM_SP_/Olg_CT_ spots (**SF11E-F**) and showed greater overlap with the *PLP1*-module than the *GLUL*-module, consistent with their co-expression in spots containing the oligodendrocyte marker *PLP1* (**SF18A**). In contrast, the *GLUL*-module DEGs were prevalent in Ast_CT_ spots and included genes associated with astrocyte-mediated oxygen delivery to neurons (Lia et al. 2023; Yaxing Chen et al. 2020) (**SF11F**). Disorder-associated differential expression within the *GLUL*-module was most pronounced in females with MDD and males with BD (**SF14B**). These included elevated Ast_CT_ DEGs associated with cellular responses to oxygen deprivation and metabolic demand (*VEGFA, VASN* and *TIPARP* (Y. Liu et al. 1995; Choksi et al. 2011; L. Zhang et al. 2020)) and vascular remodeling and angiogenesis (*APOLD1, VEGFA, EDN1,* and *ANGPTL4* (Rousseau et al. 2000; Sharma et al. 2018; Gras et al. 2016; Qiu et al. 2021; Chaube et al. 2023; Stritt et al. 2023; Fan et al. 2023)) (**Figure 4C**). These findings suggest that astrocyte-associated transcriptional responses in females with MDD and males with BD may reflect altered oxygen and metabolic demands in the dlPFC. Although *GLUL*-module DEGs implicate similar biological processes across disorders, many of the upregulated genes differed between MDD and BD, suggesting sex- and diagnosis-specific astrocyte profiles (**Figure 4D**). In males with BD, these changes were principally associated with neurovascular coupling, whereas females with MDD also exhibited elevated expression of *DDIT4* and *PFKFB3*, genes implicated in neuroinflammation and impaired GABAergic signaling (Chicote-González et al. 2025; Lv et al. 2015).

An astrocyte-associated inflammatory transcriptional response in females with MDD was evidenced by *IFITM3-*module DEGs, which were broadly upregulated across dlPFC domains and in Ast_CT_ (**Figure 4E-F**). *IFITM3*-module DEGs that were elevated in females with MDD included genes involved in neuroimmune signaling and glial activation (*IFITM3*, *CEBPD*, and *SRGN* (Ni et al. 2025; Prakash et al. 2024; Ko et al. 2014; Qian et al. 2024), endothelial function and BBB permeability (*YBX3* (Nie et al. 2012; W. Jiang et al. 2026; L. Liu et al. 2015)), and neurovascular inflammation (*CEBPD*, *OSMR* and *IL1R1* (S.-M. Wang et al. 2018; Ruprecht et al. 2001; Takata et al. 2019; Konsman et al. 2004; Luís et al. 2022)). *IFITM3*-module DEGs overlapped with the regulons of inflammation-responsive transcription factors CEBPD, JUN, and FOS (Raivich 2008; Karakaslar et al. 2023; Ko et al. 2015; Pulido-Salgado et al. 2015) (**Figure 4G**). Notably, *CEBPD*, a transcription factor involved in immune and inflammatory responses (S.-M. Wang et al. 2018; Ko et al. 2014), was itself a component of the *IFITM3*-module and, reciprocally, *IFITM3* was a component of the CEBPD-regulon. Together, these findings suggest a potential regulatory relationship between *CEBPD* and *IFITM3* in the inflammatory transcriptional response observed in females with MDD.

Inflammation is a homeostatic response to injury, but persistent activation of innate immune cells, including resident microglia and infiltrating macrophages, can impair NVU and neuronal function (Kempuraj et al. 2024). The *CD74*-module captured sparsely expressed DEGs associated with microglial activation and inflammatory immune responses that exhibited distinct diagnosis-by-sex patterns(**Figure 2G**, **SF18B-C**). Eleven *CD74-*module DEGs (e.g. *C1QB*) were increased in females with MDD (**SF14**, **SF18B**), providing transcriptional evidence of microglial immune activation in females with MDD that is consistent with *IFITM3*- and *CEBPD*-associated inflammatory responses (**Figure 4E-G**) and previous associations between inflammatory immune activation and mood disorders (Pinzi et al. 2025; Dudek et al. 2021). In contrast, seventeen *CD74*-module DEGs (e.g. *CX3CR1*) were decreased in males with BD (**SF18B**). These opposing patterns suggest distinct microglial immune-associated transcriptional responses in females with MDD and males with BD. Although previous studies have reported inconsistent evidence for microglial and inflammatory involvement in BD (Giridharan et al. 2020; Chaves-Filho et al. 2024), our findings suggest that biological sex may contribute to this heterogeneity and should be considered when interpreting microglial transcriptional changes in BD.

Taken together, our transcriptional findings implicate altered BBB integrity across both MDD and BD, with the exception of males with MDD. In females with MDD, we identified additional evidence of astrocyte reactivity and increased expression of innate immune-associated genes, consistent with an inflammatory response in the NVU. These inflammatory transcriptional responses were distinct from the oligodendrocyte-associated transcriptional changes identified in white matter (**Figure 3**), suggesting that inflammatory and stress-associated programs differ across cellular and spatial contexts in mood disorders. Although astrocyte-associated transcriptional changes were also evident in males with BD, this group exhibited decreased innate immune markers without the coordinated inflammatory transcriptional response observed in females with MDD. Our data support a diagnosis- and sex-specific link between mood disorders, BBB integrity, and inflammation that is consistent with the established physiological effects of stress linking NVU dysfunction with inflammation and depression (Dion-Albert et al. 2022; Welcome 2020; I.-B. Kim et al. 2022).

### 2.5 Differential expression of inhibitory neuron markers in mood disorders

Inhibitory neurons in cortical and subcortical regions have been strongly implicated in the behavioral outcomes of chronic stress and stress-related neuropsychiatric disorders, including post-traumatic stress disorder, anxiety disorder, MDD, and BD (Girgenti et al. 2019; Ghosal et al. 2017; Tomoda et al. 2022; Fogaça and Duman 2019; Jaffe et al. 2022; O’Connell et al. 2025; Hwang et al. 2025; Strom et al. 2026). Stress activates the hypothalamic-pituitary-adrenal (HPA) axis through hypothalamic release of corticotropin-releasing hormone (encoded by *CRH*), ultimately leading to increased circulating cortisol and glucocorticoid signaling. Although CRH neurons in the paraventricular nucleus of the hypothalamus are the primary source of CRH involved in HPA-axis activation, inhibitory neuron populations in the PFC also express *CRH*. Cortical *CRH* expression is associated with stress resilience in mice (Y.-T. Wang et al. 2025) and is decreased in donors with MDD and individuals who died by suicide (Pandey et al. 2019; Oh et al. 2022). Consistent with this previous literature, we found significantly decreased expression of *CRH*, as well as decreased expression of inhibitory neuron subtype markers *SST* and *CORT*, across domains in females with MDD and in donors of both sexes with BD (**Figure 5A-B**). *SST* and *CORT* encode highly related neuropeptides (somatostatin and cortistatin, respectively) that have been repeatedly implicated in inhibitory neuron changes in depression (Lin and Sibille 2015; Fee et al. 2017; Fuchs et al. 2017; Spier and de Lecea 2000; J. Jiang et al. 2018; Yao et al. 2022).

**Figure 5.**
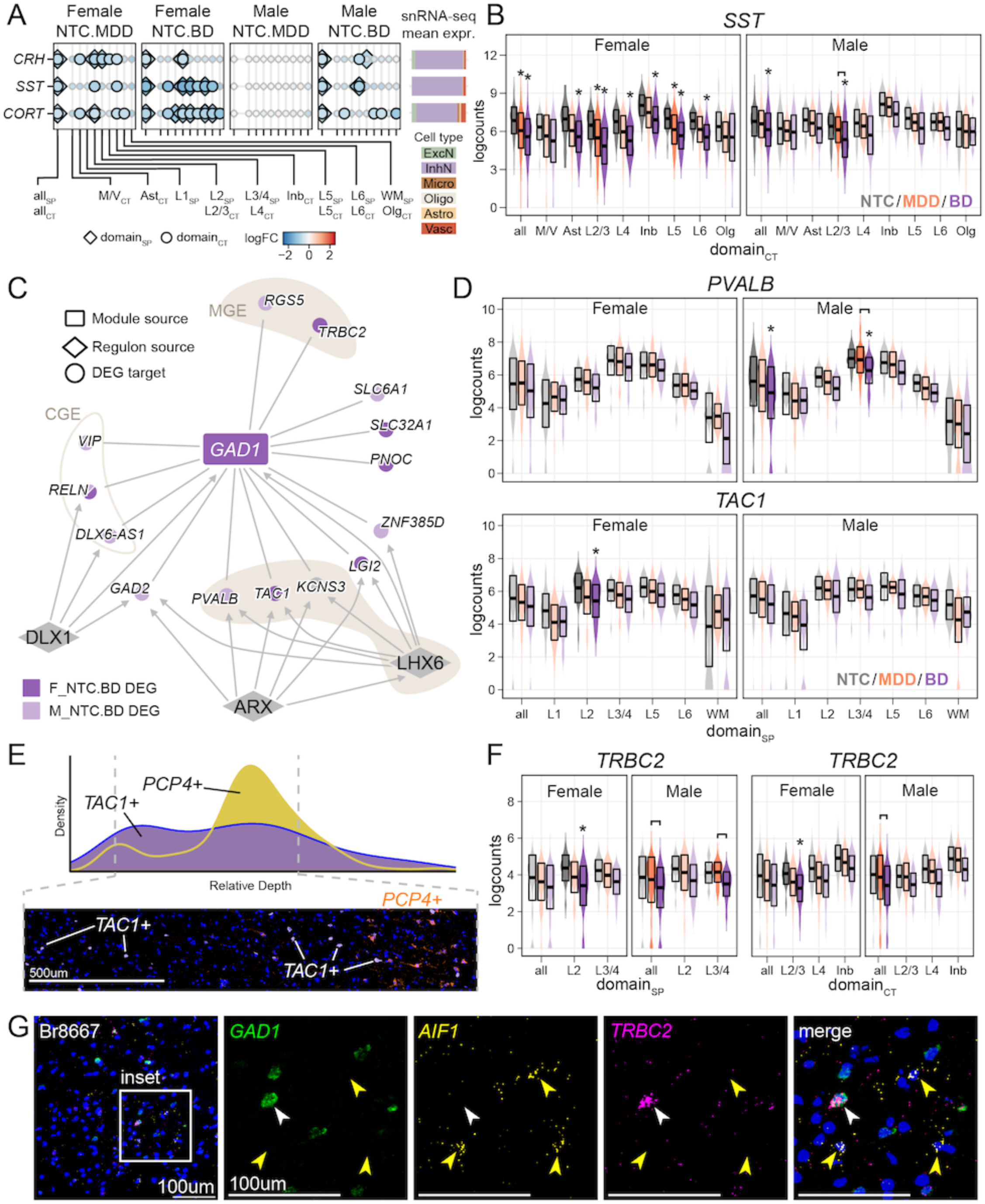
Differential expression of inhibitory neuron markers in mood disorders. (**A**) Dotplot of neuropeptide DEGs across domain annotations (x-axis, ‘all’ represents whole-tissue model results) where large size indicates DEG status, shape indicates domain annotation strategy, and color reflects log_2_ fold change (logFC). Fold change is calculated for each diagnosis-by-sex group, with comparisons to NTC shown in facets. (**B**) Violin and boxplots of pseudobulk expression (y-axis) for *SST*. Domain_CT_ DE results are shown for the whole-tissue model (‘all’) and the domain-restricted model (x-axis). Statistical significance (*t*-test (adj. *p*<.05) versus sex-matched control group indicated by asterisk. Bar symbols indicating statistical significance (adj. *p*<.05) between MDD and BD. (**C**) Network plot of all *GAD1*-module DEGs, their predicted regulons, and all DEG components of the three inhibitory neuron regulons (DLX1, ARX, and LHX6). Circle nodes represent DEG components, with a rectangular node for *GAD1* indicating its status as source of the GAD1-module. Diamond nodes represent regulon transcription factor sources and regulon edges include arrows to reflect directed regulatory relationships. Node color indicates DE group (light purple color = M_NTC.BD, dark purple color = F_NTC.BD, grey = neither M_NTC.BD nor F_NTC.BD). (**D**) Violin and boxplots of pseudobulk expression (y-axis) for *GAD1*-module DEGs and parvalbumin neuron markers *PVALB* and *TAC1*. Domain_SP_ DE results are shown for the whole-tissue model (‘all’) and the domain-restricted model (x-axis). Statistical significance (*t*-test (adj. *p*<.05) versus sex-matched control group indicated by asterisk. Bar symbols indicating statistical significance (adj. *p*<.05) between MDD and BD. (**E**) (Top) Density plot illustrating cell type abundance from 4 NTC donors (*y*-axis) across cortical layers (*x*-axis, depth relative to L1). (Bottom) Example RNAscope image (NTC M Br6432) spanning the cortical layers showing distribution of *PCP4*+ neurons (orange), *TAC1*+ neurons (white), and DAPI (blue). Scale bar = 500 micrometer (0.5 mm). (**F**) Violin and boxplots of pseudobulk expression (y-axis) for *GAD1*-module DEG *TRBC2*, for both the domain_SP_ pseudobulk model (left) and domain_CT_ pseudobulk model (right). DE results are shown for the whole-tissue model (‘all’) and for select domain-restricted models highlighting supragranular layers (x-axis). Statistical significance (*t*-test (adj. *p*<.05) versus sex-matched control group indicated by asterisk. Bar symbols indicating statistical significance (adj. *p*<.05) between MDD and BD. (**G**) Example RNAscope image (NTC F Br8667) with insets illustrating the co-expression of *TRBC2* (pink) with *GAD1* (green) (white arrowhead), but not with *AIF1* (yellow arrowheads). Scale bar = 100 micrometers.

Additional BD-associated inhibitory neuron transcriptional changes were captured by the *GAD1*-module (**Figure 2G**). All 15 *GAD1*-module DEGs were decreased in donors with BD, with no significant changes detected in donors with MDD (**Figure 5C**). These genes included markers of general inhibitory function (e.g. *GAD2*, *SLC32A1*), neurons derived from the central ganglionic eminence (CGE, e.g. *VIP*), and neurons derived from the medial ganglionic eminence (MGE, e.g. *PVALB*), some of which overlapped with the regulon of MGE master regulator LHX6 (Christodoulou et al. 2022). Although most *GAD1*-module DEGs were significant across the dlPFC tissue section (using the whole-tissue model, **Figure 2A-B**), canonical parvalbumin (PV+) neuron markers *TAC1* and *PVALB* were significantly reduced in supragranular layers (L2_SP_ and L3/4_SP_, respectively) (**Figure 5D**). Our DE results indicate that while pan-inhibitory and CGE-derived transcripts were depleted throughout the dlPFC in donors with BD, MGE-derived transcripts and LHX6-regulon targets exhibited spatially localized changes that implicate supragranular PV+ inhibitory neurons in BD.

To investigate whether our domain-restricted L2/L3/L4 results reflected differences in PV+ neuron density between supragranular and subgranular cortical domains in the dlPFC, we performed single-molecule fluorescence *in situ* hybridization (smFISH) for *TAC1* and the canonical L5 marker *PCP4* in four NTC donors (N=2M/2F). We quantified the distribution of cells expressing *TAC1* (**Methods 4.8**) and found that *TAC1*+ neurons were similarly distributed superficial to and throughout the *PCP4*+ region (**Figure 5E**). The smFISH results are consistent with prior analyses of PV+ neuron distribution (Druga et al. 2023; Becerra-Hernández et al. 2026), and suggest that the layer-specific changes in *TAC1* and *PVALB* expression in donors with BD are unlikely to be explained solely by the preferential localization of *TAC1+* cells to superficial cortical layers.

Together, our results identify BD-associated inhibitory neuron transcriptional changes, including spatially-restricted reductions in PV+ neuron markers within supragranular layers. Another *GAD1*-module DEG, *TRBC2*, was significantly downregulated in females with BD specifically in L2/3_CT_ and L2_SP_ (**Figure 5F**). Although *TRBC2* encodes a core component of T-cell receptor beta and has established roles in T-cell cancers (Ferrari et al. 2024; Ge et al. 2025), our SRT data showed spot-level co-expression of *TRBC2* and *GAD1*, and human PFC snRNA-seq data (Ruzicka et al. 2024) indicated that *TRBC2* expression was greatest in inhibitory neurons, specifically SST+ neurons (**SF19C ii-iii**). To confirm that this sex- and diagnosis-specific DEG was expressed in inhibitory neurons, we used smFISH to examine co-expression of *TRBC2* with *GAD1* and the immune cell marker *AIF1* (encoding IBA1) in four NTC donors (N=2M/2F, **Methods 4.8**). Although both *AIF1* and *TRBC2* were sparsely expressed, the *TRBC2* signal colocalized with *GAD1,* but not *AIF1* (**Figure 5G, SF19D**). These results support an unexpected association of *TRBC2* with inhibitory neurons in the dlPFC, and a female-specific downregulation in BD.

In summary, our results support existing evidence of the involvement of stress-sensitive inhibitory neuron populations in mood disorders. Inhibitory neuron-associated transcriptional changes were more extensive in BD, with reduced expression of inhibitory neuron markers in both male and female donors. However, reductions in the stress-sensitive neuropeptides *SST*, *CORT*, and *CRH* were shared by females with MDD and donors with BD of both sexes. Our spatially-resolved data further localized reduction in the PV+ neuron markers *PVALB* and *TAC1,* as well as the inhibitory neuron-associated transcript *TRBC2* to supragranular layers of the dlPFC, which are important for intracortical connectivity.

### 2.6 Genetic risk and DE converge on intracellular signaling cascades in excitatory neurons

Biological pathways that have consistently emerged at the nexus of stress, inflammation, and mood disorders include glucocorticoid signaling, innate immune activation, and MAPK signaling (I.-B. Kim et al. 2022; Pinzi et al. 2025). However, it remains unclear whether disorder-associated transcriptional changes in these pathways reflect downstream consequences of living with the disorder or are influenced by genetic variation that contributes to disease risk. To begin to distinguish these possibilities, we integrated paired genetic and expression data from our donors to identify relationships between genetic variation and diagnosis- and sex-associated transcriptional changes.

We performed *cis* expression quantitative trait loci (eQTL) analysis to identify genomic variants independently associated with gene expression in our dlPFC SRT data (**Figure 6A**, **Methods 4.9.1**) (Taylor-Weiner et al. 2019). We identified thousands of SNP-gene associations, occurring predominantly across neuron-rich domains_CT_ (**Figure 6B**, **ST6**). To identify genes with both diagnosis-by-sex DE and genetic associations, we examined genes with at least one significant eQTL (eGenes) that were also significant in our diagnosis-by-sex DE models. We identified 50 unique eGene-DEGs across all domain_CT_ contexts (**Figure 6C**), including several that were predominantly expressed in cell types comprising the NVU and associated with glial responses (e.g. *ANP32B*, *NEAT1*, *IFITM3*). Expression of *IFITM2*, a component of the *IFITM3*-module and the FOS regulon (**Figure 4G**), and *IFITM3* were each increased in females with MDD and associated with independent variants specifically in the Ast_CT_ domain (rs10751647 *q* = 0.0072 and rs61876236 *q* = 0.034, respectively) (**Figure 6D-E**). These eQTL associations indicate that genetic variation is statistically linked with the expression of genes involved in the astrocytic transcriptional responses observed in females with MDD (**Figure 4**), linking genotype-dependent expression to disorder-associated glial responses.

**Figure 6.**
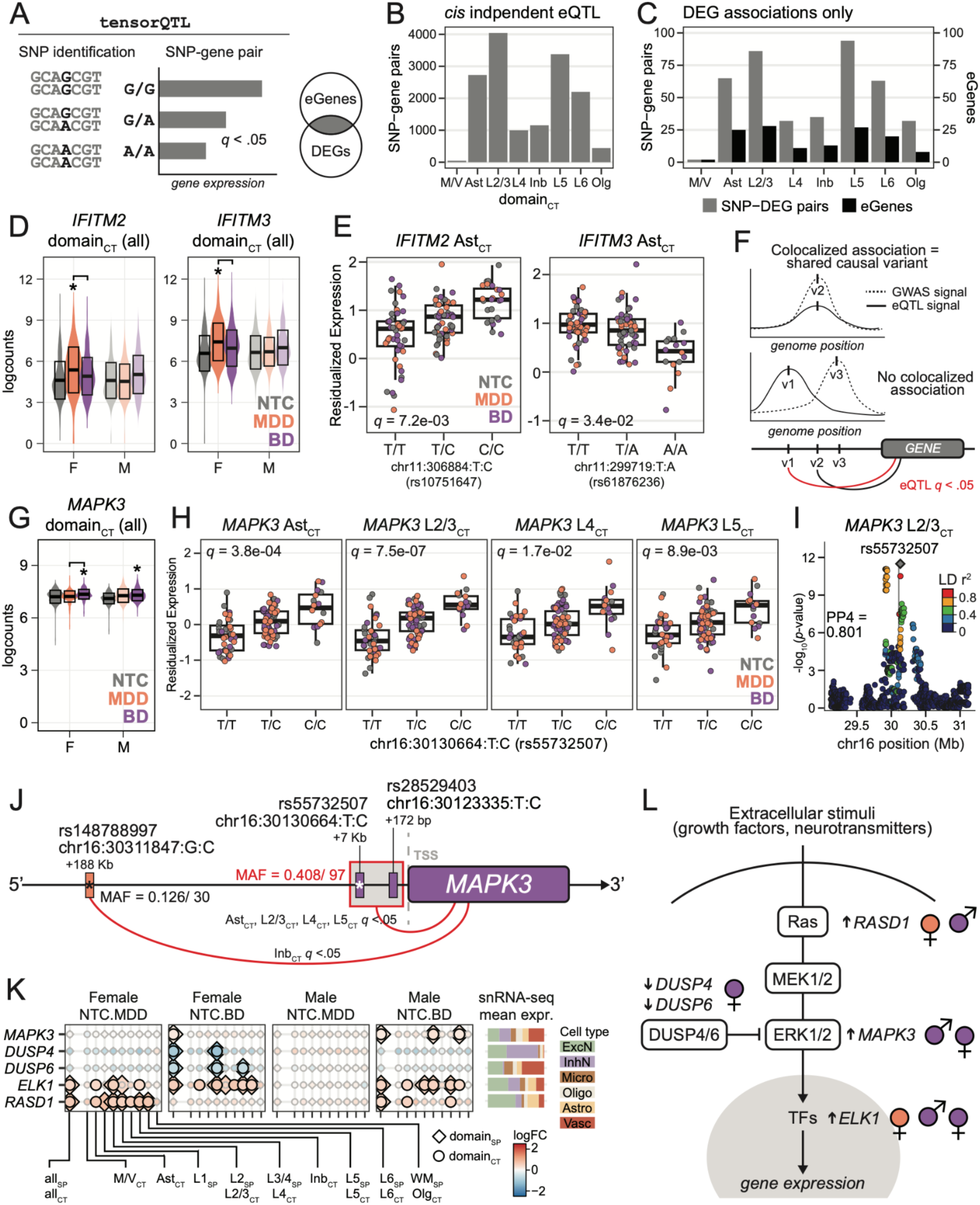
Genetic risk and differential gene expression converge on intracellular signaling cascades in excitatory neurons. (**A**) Schematic representation of `tensorQTL` workflow for expression quantitative trait loci (eQTL) analysis. Single nucleotide polymorphism (SNP) genetic variants were evaluated within the genomic region of genes for a statistical association with the expression of that gene. A gene with one or more SNP-gene pairs is called an eGene. We identified overlaps between eGenes and DEGs to highlight strong gene-by-environment influences within disease-associated DEGs. (**B**) Bar graph of number of SNP-gene pairs (*y*-axis) identified for each domain_CT_ context (*x*-axis). (**C**) Bar graph of the number of SNP-gene pairs matched to DEGs (grey, left *y*-axis) and the number of eGenes that are also DEGs (black, right y-axis) per domain_CT_ context (*x*-axis). (**D**) Violin and boxplot of *IFITM2* (left) and *IFITM3* (right) for whole-tissue model domain_CT_. Statistical significance (*t*-test (adj. *p*<.05) versus sex-matched control group indicated by asterisk. Bar symbols indicating statistical significance (adj. *p*<.05) between MDD and BD. (**E**) Box plots show genotype-associated Ast_CT_ expression patterns for *IFITM2* and rs10751647 (left) and *IFITM3* and rs61876236 (right). (**F**) Schematic of colocalization analysis to identify shared causal variants for eQTL and GWAS signals. For the three toy variants (v1, v2, v3), eQTL and GWAS signals are only colocalized for v2. Additionally, eQTL *q*-values are indicated (red, *q* < .05) to reflect that colocalization can produce significant association for non-significant eQTL signals. (**G**) Violin and boxplot of *MAPK3* for whole-tissue model domain_CT_. Statistical significance (*t*-test (adj. *p*<.05) versus sex-matched control group indicated by asterisk. Bar symbols indicating statistical significance (adj. *p*<.05) between MDD and BD. (**H**) Box plots show significant genotype-associated expression patterns for *MAPK3* (*y*-axis, residualized expression) and variant rs55732507 (*x*-axis) for Ast_CT_, L2/3_CT_, L4_CT_, L5_CT_. Given the perfectly correlated LD of the variants rs55732507 and rs28529403 in our cohort, the model results (including statistical tests and residualized expression) are therefore the box plot is identical for rs28529403. (**I**) Colocalization plot of *MAPK3* variant rs55732507 (diamond) in L2/3_CT_. Each point indicates the position (*x*-axis, megabases (Mb)) of a single SNP on chromosome 16 within the *cis* window of *MAPK3*, with vertical grey line indicating the position of the *MAPK3* gene (chr16:30,114,105-30,123,506, negative strand). *Y*-axis presents the nominal, non-independent eQTL *p*-value (-log10 transformed). Point color represents the binned linkage disequilibrium (LD) r^2^ value between each variant and rs55732507. PP4 is the locus-level posterior probability that GWAS_BD_ and *MAPK3* eQTL signals share a causal variant (strong evidence at > 0.8). (**J**) Summary schematic of significant results from genetic and transcriptomic integration of *MAPK3*, with variants represented by small vertical rectangles along the genomic axis and the genomic region of MAPK3 represented by a large rounded rectangle. Distance along the genomic axis is not to scale. Expression of *MAPK3*, a BD DEG (indicated by purple), was significantly associated with two promoter-proximal variants by eQTL analysis (red arc, contexts labeled) that exhibited identical minor allele frequency (MAF) in our donors and therefore had perfectly correlated LD (indicated by light grey box with red border). Both of these variants reached the exploratory threshold for GWAS_BD_ support (indicated by purple color), and one variant exhibited significant colocalization of eQTL and GWAS_BD_ signals (white asterisk). A third, distal variant was significantly associated with *MAPK3* expression in Inb_CT_ domain (red arc), reached exploratory threshold for GWAS_MDD_ support (indicated by orange color), and exhibited significant colocalization of eQTL and GWAS_MDD_ signals (black asterisk). (**K**) Dotplot of Ras/MEK/ERK neuronal DEGs across domain annotations (x-axis, ‘all’ represents whole-tissue model results) where large size indicates DEG status, shape indicates domain annotation strategy, and color reflects log_2_ fold change (logFC). Fold change is calculated for each diagnosis-by-sex group, with comparisons to NTC shown in facets. (**L**) Schematic of Ras/MEK/ERK neuronal DEGs signaling pathway relationships.

To connect eQTLs with genetic risk for mood disorders, we obtained GWAS summary statistics for MDD (GWAS_MDD_), BD (GWAS_BD_), and schizophrenia (GWAS_SCZ_) to identify variants associated with genetic risk for these conditions (Adams et al. 2025; O’Connell et al. 2025; Trubetskoy et al. 2022) (**SF20**). We included GWAS_SCZ_ because of its greater statistical power relative to GWAS_MDD_ and GWAS_BD_ and the substantial shared heritability among these disorders (Schulze et al. 2014; Ruderfer et al. 2018; Bigdeli et al. 2021; Grotzinger et al. 2026). To link genetic risk for mood disorders with gene expression, we used colocalization analysis, which evaluates whether eQTL and GWAS association signals at the same locus are consistent with a shared causal variant (Giambartolomei et al. 2014) (**Figure 6F**, **Methods 4.9.2**). Across the three GWAS datasets, colocalization analysis identified 66 variants at 54 genes with evidence of a shared genetic signal in specific spatial contexts (PP4 > 0.8) (**ST7**). Because colocalization analysis alone does not establish that a shared variant is causally linked to diagnosis, we annotated whether each strong colocalized variant met the strict (*p* < 5e-08; GWAS_MDD_, GWAS_BD_, GWAS_SCZ_) or exploratory (*p* < 1e-05; GWAS_MDD_, GWAS_BD_) GWAS significance threshold (**Methods 4.9.3**). Of these colocalized loci, 44 variants met at least one threshold, with 19 variants exhibiting GWAS_MDD_ support and 14 variants exhibiting GWAS_BD_ support across all spatial contexts (**SF20D**, **ST7**). Of these potential causal variants, four were linked to mood disorder DEGs (*MAPK3* (two variants), *FBLN7*, and *PTP4A3*, **SF20E**).

Integrated genetic and transcriptomic analyses highlighted *MAPK3* as a point of convergence between BD-associated DE and BD genetic risk (**Figure 6G-J**). *MAPK3* was significantly increased in the tissue-wide model in both male and female donors with BD (**Figure 6G**). Two variants (rs55732507 and rs28529403) within 7kb of the *MAPK3* transcription start site exhibited exploratory associations with BD risk (both GWAS_BD_ *p* < 7e-08) and significant associations with SCZ risk (both GWAS_SCZ_ *p* < 7e-10). These variants exhibited perfectly correlated allelic frequencies in our cohort and were significantly associated with increased *MAPK3* expression in several spatial contexts (Ast_CT_, L2/3_CT_, L4_CT_, L5_CT_; minor allele = C, eQTL max *q* = 0.0173) (**Figure 6H**). One variant (rs55732507) exhibited colocalization of GWAS_BD_ and *MAPK3* eQTL signals in Ast_CT_, L2/3_CT_, Inb_CT_, and L5_CT_ (PP4 = 0.80-0.92) (**Figure 6I**), supporting a shared genetic signal underlying BD risk and regulation of *MAPK3* expression. A third variant (rs148788997) at the *MAPK3* locus was associated with *MAPK3* expression in Inb_CT_ (eQTL *q* = 0.00115), exhibited an exploratory association with MDD risk (risk allele = C, GWAS_MDD_ = 5.91e-07), and showed colocalization of the GWAS_MDD_ and eQTL signals (PP4 = 0.94) (**Figure 6J**).

The convergence of genetic and transcriptional associations at *MAPK3* was accompanied by broader disorder-associated alterations in the MAPK signaling cascade. *MAPK3* encodes ERK1, a central kinase in Ras/MEK/ERK signaling that has been genetically and transcriptionally linked to mood disorders (Yuan et al. 2010; Malki et al. 2015; Shao et al. 2024; Chauhan and Jain 2025). Additional DEGs implicated multiple components of this pathway, including the ERK1 inhibitors *DUSP4* and *DUSP6*, the ERK1-regulated transcription factor *ELK1*, and the glucocorticoid-sensitive Ras family GTPase *RASD1* (**Figure 6K**). The direction of these transcriptional changes is consistent with a potential increase in Ras/MEK/ERK signaling (**Figure 6L**). Together, these findings identify Ras/MEK/ERK signaling as a point of convergence between disorder-associated transcriptional changes and genetic risk, with particularly strong genetic support in BD.

## 3 Discussion

Here, we used spatial transcriptomics to characterize the molecular architecture of the postmortem human dlPFC across 119 brain donors with MDD or BD and neurotypical controls, with approximately equal representation of males and females in each diagnostic group. By preserving cortical architecture and integrating complementary spatial domain annotations, we identified shared and distinct transcriptional changes across white matter, the neurovascular unit (NVU), and neuronal compartments. Our findings revealed prominent sex-dependent differences in MDD, including inflammatory responses within the NVU and altered oligodendrocyte-associated transcriptional programs in females. Across mood disorders, we identified shared alterations in inhibitory neuropeptides and transcriptional evidence of neurovascular dysfunction, while broader inhibitory neuron changes were apparent in BD. Integration with donor genotype data further implicated *MAPK3* as a point of convergence between genetic risk for BD and diagnosis-associated transcriptional changes, further implicating dysregulation of Ras/MEK/ERK signaling in BD. Together, these findings demonstrate that the molecular alterations associated with mood disorders are strongly dependent on biological sex and spatial context, providing insight into both shared and disorder-specific mechanisms in the human dlPFC.

Although sex-specific transcriptional changes were evident in certain cellular contexts in BD, we observed a much more pronounced difference between males and females with MDD. Specifically, females with MDD exhibited extensive changes in gene expression, whereas relatively few significant DEGs were identified in males with MDD (**SF7**, **SF8**). Previous studies in dlPFC have reported sex-specific transcriptional changes in MDD, including altered expression in males (Labonté et al. 2017; Mansouri et al. 2023; Nagy et al. 2020; Maitra et al. 2023). However, subsequent re-analysis of previously published bulk RNA-seq data (Labonté et al. 2017) found that male-associated transcriptional changes did not retain statistical significance following correction for multiple comparisons (de Souza et al. 2024). Previous bulk RNA-seq analyses also demonstrated substantial regional heterogeneity in MDD-associated transcriptional changes, with considerably more differential expression in the dorsal anterior cingulate and medial amygdala compared to the dlPFC (Jaffe et al. 2022). Together these findings raise the possibility that the molecular manifestations of MDD differ across both biological sex and brain region, and that the limited transcriptional changes observed in males with MDD in the dlPFC may not reflect the extent of molecular pathology elsewhere. Given known sex differences in the clinical presentation of MDD, including higher suicidality, impulsivity, and comorbid substance use in males (Oliffe et al. 2019; Cavanagh et al. 2017), and previous studies identifying MDD-associated transcriptional alterations in other cortical and subcortical regions involved in emotional regulation and behavioral control, including the orbitofrontal cortex, anterior cingulate cortex, and amygdala, (Jaffe et al. 2022; Daskalakis et al. 2024; Kamimura et al. 2022; Fröhlich et al. 2024; Zandi et al. 2022; Goes et al. 2025), our findings raise the possibility that molecular alterations in males with MDD may be more prominent in brain regions outside the dlPFC.

In females with MDD, we identified a coordinated inflammatory transcriptional response within the NVU, characterized by astrocyte reactivity, inflammatory signaling, and immune activation (**Figure 4**, **SF18**). Inflammation has been implicated in the pathophysiology of depression (I.-B. Kim et al. 2022; Pinzi et al. 2025; Beurel et al. 2020), and our findings extend this literature by identifying a prominent female-specific inflammatory phenotype within the dlPFC NVU that was not observed in BD. Consistent with this sex-specific association, a recent meta-analysis found that elevations in peripheral inflammatory immune markers in depression were frequently restricted to female participants (Jarkas et al. 2024). Our analyses further identified *CEBPD* and *IFITM3* as potential central components of a coordinated inflammatory program. Enriched CEBPD motifs in the regulatory regions of *IFITM3* and other inflammatory response genes suggest that CEBPD may contribute to the transcriptional regulation of NVU inflammation in females with MDD (**Figure 4**). These associations suggest a potential regulatory relationship between CEBPD and *IFITM3,* consistent with the established role of CEBPD in promoting reactive glial inflammation (S.-M. Wang et al. 2018; Ko et al. 2015), and implicate this pathway in MDD-associated NVU inflammation.

Integrating genetic and transcriptomic data identified *cis*-eQTLs associated with *IFITM2* and *IFITM3* expression in astrocyte-enriched regions (**Figure 6**). Although these variants (rs10751647, rs61876236) have not been associated with MDD risk, they suggest that genetic variation may influence components of the inflammatory transcriptional response observed in females with MDD. *IFITM3* is induced by inflammatory cytokines and contributes to antiviral defense through phosphatidyl inositol signaling and lipid raft organization (J. Lee et al. 2020; Klein et al. 2023). In Alzheimer’s disease and cerebral ischemia-reperfusion injury, increased *IFITM3* expression in vascular cells and astrocytes has been linked to neuroinflammation through modulation of γ-secretase activity (Feng et al. 2025; Ni et al. 2025; Hur et al. 2020). Together, these findings suggest that genotype-dependent regulation of *IFITM3* expression may contribute to inflammatory responses within the NVU in females with MDD, highlighting a potential mechanism for future investigation.

Beyond the inflammatory response observed in females with NDD, our findings implicate broader NVU dysfunction across mood disorders. Previous studies have identified BBB and NVU alterations in MDD and BD, while preclinical models have demonstrated that stress-induced changes in neurovascular gene expression and BBB integrity can contribute to depression-related behaviors (Menard et al. 2017; Dion-Albert et al. 2022; Sun et al. 2024; Solich et al. 2025; Wu et al. 2022; Wakonigg Alonso et al. 2024). Consistent with these findings, we identified reduced expression of genes involved in BBB integrity and vascular function, including *A2M*, *TNFSF10*, and *ABCG2*, in females with MDD and donors with BD of both sexes (**Figure 4**). However, the accompanying glial transcriptional responses differed by diagnosis and sex. Females with MDD exhibited astrocytic inflammatory and innate immune responses, whereas males with BD showed astrocytic changes associated predominantly with neurovascular coupling and reduced expression of microglial immune markers. Previous studies have also implicated astrocytic and microglial alterations in BD, although biological sex has not consistently been examined (Wei et al. 2024; Giridharan et al. 2020; Chaves-Filho et al. 2024). Notably, our findings in males with BD are consistent with previous transcriptomic studies of the dlPFC and the subgenual anterior cingulate cortex using male-dominated cohorts (Xia et al. 2024; Zandi et al. 2022). Together, these findings suggest that shared alterations in neurovascular function may be accompanied by distinct glial responses depending on diagnosis and biological sex.

The effects of stress on *SST* expression in the PFC are well-documented (Lin and Sibille 2015; Banasr et al. 2017; Girgenti et al. 2019). Consistent with this literature, we identified reduced expression of *SST* and two other inhibitory neuron neuropeptides, *CORT* and *CRH*, in females with MDD and donors with BD of both sexes (**Figure 5**). Reductions in *SST* expression and the abundance of SST+ neurons have been reported across cortical and subcortical regions in neuropsychiatric and mood disorders (Sibille 2017; Liguz-Lecznar et al. 2022). However, findings in the dlPFC of donors with BD have been inconsistent; our study and Fung et al., identified decreased *SST* (Fung et al. 2014), whereas other studies reported no significant changes in BD, despite reductions in MDD or SCZ (Sibille et al. 2011; Okuda et al. 2024; Dienel et al. 2023). Adverse life events can alter neuropeptide expression in the PFC (W. Li et al. 2018), suggesting that neuropeptide biosynthesis may contribute to the vulnerability of stress-sensitive inhibitory signaling in neuropsychiatric disorders (Casello et al. 2022). Together with our NVU findings, these results implicate stress-associated transcriptional changes across both inhibitory neurons and neurovascular compartments as a shared feature of MDD and BD.

Stress-induced outcomes in the brain are mediated, in part, by glucocorticoid-stimulated gene expression (Joëls et al. 2012; Mifsud and Reul 2018), and we identified increased expression of the glucocorticoid-sensitive gene *RASD1* (Kemppainen and Behrend 1998) in males with BD and females with MDD (**Figure 6**). *RASD1* encodes a Ras family GTPase (DEXRAS1), which has been implicated in nitric oxide signaling and NMDA receptor-dependent iron homeostasis in cortical neurons (Fang et al. 2000; Cheah et al. 2006; Yong Chen et al. 2013; White et al. 2016). *RASD1* expression also influences Ras/MEK/ERK signaling and resulting transcription factor response, promoting ELK1 transcriptional activity (Timothy E. Graham et al. 2002; T. E. Graham et al. 2004; Greenwood et al. 2016). Glucocorticoid- and ERK-dependent ELK1 activity is associated with stress-induced and depressive behaviors (Gutièrrez-Mecinas et al. 2011; Apazoglou et al. 2018). Together with increased *ELK1* expression in females with MDD and donors with BD of both sexes, these findings implicate altered glucocorticoid-responsive Ras/MEK/ERK signaling as a potential link between stress and neuronal dysfunction across mood disorders.

In our study, donors with BD exhibited transcriptional changes consistent with inhibitory neuron dysfunction, including reduced expression of the PV+ inhibitory neuron markers *TAC1* and *PVALB* localized to the supragranular layers of the dlPFC (L2/L3/L4; **Figure 5**). These findings are consistent with previous reports of reduced *PVALB* expression in the dlPFC of donors with BD but not MDD (Sibille et al. 2011; Okuda et al. 2024). PV+ neurons provide perisomatic inhibition of excitatory neurons and are critical for regulating cortical network activity, functions that are implicated in neuropsychiatric disorders (Ferguson and Gao 2018; Selten et al. 2018). The reductions of *TAC1* and *PVALB* in the supragranular layers suggests that altered inhibitory regulation of cortical microcircuits may be particularly relevant to BD-associated dlPFC dysfunction (K.-M. Han et al. 2019; Y. Lee et al. 2018).

In addition to the inhibitory neuron and NVU-associated alterations, we identified prominent female-specific transcriptional changes in white matter-enriched spatial domains (**Figure 3**). Females with both MDD and BD exhibited increased expression of genes associated with cell survival and stress responses, but showed distinct oligodendrocyte-associated transcriptional profiles. In females with BD, reduced expression of myelin-associated genes suggested impaired myelination, whereas females with MDD exhibited increased expression of stress-responsive genes, including *SGK1* and *NEAT1*. Although mature oligodendrocytes are primarily responsible for myelination, oligodendrocyte precursor cells (OPCs) play a critical role in maintaining myelinating oligodendrocytes and OPC loss in the PFC has been linked to depressive-like symptoms (Birey et al. 2015). OPCs engage in a variety of functions and participates in inter-dependent networks with other white matter-resident glia (e.g. microglia, vascular cells) (Foerster et al. 2019; J. Li et al. 2025), including modulating immune response and can contribute to altered myelination in response to inflammation (Q. Wang et al. 2025; Kirby et al. 2019). Given these roles, OPCs may be involved in the shared stress-associated response programs and distinct myelination-related changes observed in females with MDD and BD, consistent with previous evidence implicating oligodendrocyte dysfunction in mood disorders (Favre et al. 2019; Rajkowska et al. 2015; Abraham et al. 2022; Ali et al. 2024; Zhou et al. 2024).

To investigate whether genetic variation contributes to the diagnosis-associated transcriptional changes we identified, we integrated donor genotype data with spatially resolved gene expression. Unlike bulk tissue eQTL analyses, our spatial approach enabled us to identify associations between genetic variants and gene expression in specific cellular contexts. Among these findings, *MAPK3* emerged as particularly interesting, with distinct variants associated with its expression and genetic risk for MDD and BD (**Figure 6**). While MDD and BD share some genetic risk (Keser et al. 2026; Blokland et al. 2022; Grotzinger et al. 2026), we identified different *MAPK3* variants associated with each disorder. Specifically, an exploratory GWAS_MDD_ variant (rs148788997) was associated with *MAPK3* expression in inhibitory neuron-enriched regions, whereas a separate exploratory GWAS_BD_ variant (rs55732507) was associated with expression across multiple neuronal and astrocyte-enriched regions. These findings illustrate how distinct genetic variants linked to risk of mood disorders may influence the expression of a common gene in different spatial and cellular contexts.

The genetic association between *MAPK3* and BD was notable given that *MAPK3* expression was increased in donors with BD of both sexes (**Figure 6**). Colocalization analysis evaluates whether genetic associations with disease risk and gene expression within the same locus are consistent with a shared causal variant, providing stronger evidence that genetic risk and transcriptional regulation may be linked. The BD-associated variant rs55732507 exhibited colocalization between GWAS_BD_ and *MAPK3* eQTL signals in neuron-enriched (L2/3_CT_, L5_CT_) and astrocyte-enriched (Ast_CT_) domains (**Figure 6**). This variant has also been associated with brain morphology, amygdala volume, and performance on a higher-level cognitive task, further supporting its potential relevance to neuronal function (van der Meer et al. 2020; Y. Ji et al. 2025; de la Fuente et al. 2021). *MAPK3* encodes ERK1, a critical component of the Ras/MEK/ERK pathway, and we identified additional transcriptional changes consistent with altered regulation of this pathway. In particular, expression of the ERK inhibitors *DUSP4* and *DUSP6* were decreased in females with BD, consistent with previous evidence indicating female-specific importance of DUSP6 in regulation of ERK signaling in BD (S. H. Kim et al. 2012; Labonté et al. 2017).

Previous studies have reported reduced ERK signaling in MDD and BD, and stimulation of ERK signaling is associated with improvements in depression-related behaviors and circadian rhythm regulation (Yuan et al. 2010; M. J. McCarthy et al. 2016; J. Q. Wang and Mao 2019). Circadian rhythm disturbances are common in mood disorders (Dollish et al. 2024; X.-L. Wang et al. 2019), and several Ras/MEK/ERK pathway genes that are altered in our data (*MAPK3, DUSP4, DUSP6, RASD1, ELK1*, **Figure 6**) have established roles in circadian rhythm regulation (Hamnett et al. 2019; Cheng and Obrietan 2006; M. J. McCarthy et al. 2016). While the dlPFC is not a primary circadian pacemaker region, it supports wakefulness, attention, and goal-directed behavior, functions that may be affected by circadian disruption (Scammell et al. 2017; Krause et al. 2017). The increased expression of *MAPK3*, *RASD1* and *ELK1*, together with decreased expression of the ERK inhibitors *DUSP4* and *DUSP6* is consistent with coordinated changes that would increase Ras/MEK/ERK signaling. In the context of previous reports of reduced ERK in mood disorders (Yuan et al. 2010; M. J. McCarthy et al. 2016; J. Q. Wang and Mao 2019), these transcriptional changes may reflect a compensatory response Together, our findings implicated Ras/MEK/ERK signaling as a potential molecular link between stress-responsive transcription and circadian dysregulation in MDD and BD.

Overall, our findings demonstrate that the ‘overlapping yet distinct’ clinical presentation of MDD and BD is reflected by shared and divergent molecular alterations in the human dlPFC. Across disorders, transcriptional changes converged on stress-associated signaling pathways, inhibitory neuropeptide expression, and neurovascular function, suggesting that these shared biological processes may contribute to common features of mood disorder pathology. However, the specific transcriptional changes differed substantially by diagnosis, sex and spatial context. Females with MDD exhibited prominent inflammatory responses associated with the NVU, females with mood disorders showed distinct oligodendrocyte-associated transcriptional alterations, and donors with BD exhibited broader inhibitory neuron changes. Integration of genetic information and transcriptomic data identified Ras/MEK/ERK signaling as a point of convergence between genetic risk and diagnosis-associated transcriptional changes, further implicating this critical pathway in mood disorder pathology. These findings highlight the importance of biological sex and spatial context in understanding the molecular pathology of mood disorders, demonstrating that shared biological processes may give rise to transcriptional consequences that are differently manifested in distinct cell types and spatial contexts.

### Study limitations

Clinical heterogeneity in MDD and BD is an important consideration when interpreting case-control transcriptomic studies. In MDD, differences in symptom presentation and diagnostic criteria may have contributed to the limited differential gene expression observed in male donors. In BD, diagnostic subtype (BD I vs BD II) and mood state may also influence transcriptional expression, but our study was not designed or powered to evaluate these differences. Future studies incorporating more detailed clinical phenotyping may help to determine how specific clinical characteristics relate to diagnosis- and sex-associated molecular alterations we observed here.

The spot-based Visium platform captures transcripts from multiple cells and cellular compartments, limiting the ability to attribute transcriptional changes to individual cell types. To partially address this limitation, we employed complementary spatial annotation strategies that enabled analyses across canonical cortical layers (domain_SP_) and spatial domains enriched for specific cell populations (domain_CT_). This parallel approach provided additional cellular context for interpreting diagnosis-associated transcriptional changes, including reductions in inhibitory neuron-associated genes across the tissue and spatially restricted changes in *PVALB* and *TAC1* expression in supragranular cortical layers. Importantly, Visium preserves the spatial organization of the tissue, captures extranuclear transcripts, and enables co-expression analysis of genes expressed in neighboring cells. We leveraged these features to identify spatially organized transcriptional programs, including those associated with multicellular compartments, such as the NVU. Future studies using higher resolution SRT approaches will help resolve the contribution of individual cell populations to these coordinated transcriptional changes.

Several donor-level factors may influence gene expression, but could not be fully evaluated in our analyses. Pharmacological treatments for MDD and BD, as well as substance use, may affect transcriptional profiles; however, uncertainty regarding medication adherence and exposure to other substances limited our ability to incorporate these variables into the differential expression models. We evaluated additional donor-level covariates and included age in our DE model given its known associations with inflammation, HPA axis responsiveness, and sex hormone levels. Information on menstrual cycle phase and menopausal status was unavailable, preventing us from evaluating the contribution of hormonal factors to the observed sex-specific transcriptional differences. Similarly, the absence of acute or longitudinal measures of HPA axis activity limited our ability to relate stress-sensitive transcriptional changes directly to glucocorticoid exposure. Despite these limitations, our findings identify coherent diagnosis- and sex-associated transcriptional programs across multiple spatial contexts, providing a foundation for future studies examining how hormonal, environmental, and genetic factors contribute to the molecular alterations identified here.

## 4 Materials and Methods

### 4.1 Tissue preparation and processing

Postmortem human brain tissue from the Lieber Institute for Brain Development (LIBD) Human Brain and Tissue Repository was collected through the following sites and protocols at the time of autopsy with informed consent from the legal next of kin: the Office of the Chief Medical Examiner of the State of Maryland, under the Maryland Department of Health IRB protocol #12–24; the County of Santa Clara Medical Examiner-Coroner Office in San Jose, CA, the Departments of Pathology at Western Michigan University Homer Stryker MD School of Medicine and at the University of North Dakota School of Medicine and Health Sciences, all under WCG IRB protocol #20111080. Additional samples were consented through the National Institute of Mental Health Intramural Research Program under NIH protocol #90-M-0142, and were acquired by LIBD via material transfer agreement. Using a standardized strategy, all donors were subjected to clinical characterization and diagnosis. Details of tissue acquisition, handling, processing, dissection, clinical characterization, diagnoses, neuropathological examinations, and quality control measures have been described previously (Lipska et al. 2006). Fresh brain tissue was cut into coronal slabs, flash-frozen on dry ice, and stored at −80°C until further dissections.

#### 4.1.1 Postmortem human tissue sample preparation

Demographics for the 119 donors are listed in **ST1** and aggregate data in **Table 1**. Briefly, we selected male and female donors of European Ancestry for each of three diagnostic groups: neurotypical control (NTC), major depressive disorder (MDD), and bipolar disorder (BD). In total, our dataset consists of data from 40 NTC donors (20 male), 39 MDD donors (19 male), and 40 BD donors (20 male). Fresh frozen coronal brain slabs at the level of the anterior striatum were selected for dlPFC sub-dissections. Approximately 8 × 8 × 5 mm cortical tissue blocks encompassed Brodmann Areas (BA) 46 were dissected perpendicular to the pial surface with a hand-held dental drill to ensure the inclusion of complete laminar structure, spanning cortical layers 1-6 and white matter. To preserve the integrity of all cortical lamina, dlPFC dissections were carried out across a sulcus, maximizing the probability of retaining the most superficial layer 1 (Kwon et al. 2026; Huuki-Myers et al. 2024). Following dissection, tissue blocks were flash frozen on dry ice and stored at −80°C in sealed cryogenic bags until cryosectioning and between tissue collection rounds. Data generators for tissue processing, anatomical validation (AV), and Visium data generation were blinded to donor demographic and diagnostic information.

#### 4.1.2 Tissue processing and anatomical validation

At the time of cryosectioning, tissue blocks were acclimated inside a cryostat (Leica CM3050s) at −14°C for 15 to 30 minutes. Each tissue block was mounted onto a chuck using Optical Temperature Compound (OCT, TissueTek Sakura, Cat #4583). Approximately 50 µm of tissue was trimmed from the block to achieve a flat surface and several 10 µm sections were collected on microscope slides (VWR SuperFrost Microscope Slides, Cat #48311703) and banked at −80°C for anatomical validation and quality control (QC) work. Prior to sample collection on Visium spatial gene expression slides, candidate tissue samples were examined to validate anatomical dissections. First, visual inspection to assess gross neuroanatomical structure was performed. Visual inspection of tissue blocks relied on the presence of clear white matter/ grey matter border and a sulcus to the adjacent gyrus. In some cases, when tissue blocks did not pass visual inspection, additional experiments were performed to assess tissue morphology and presence of all 6 cortical layers and white matter. H&E staining and RNAscope smFISH were performed according to manufacturer’s instructions as previously described, and images were acquired using an Leica Aperio CS2 slide scanner or Nikon AX-R. For RNAscope, probes for established marker genes (Advanced Cell Diagnostics Bio [ACD]) were used to identify laminae in the human cortex, including *AQP4* (ACD, Cat No. 482441) as a marker gene for layer 1*, HPCAL1* (ACD, Cat No. 846051) as a marker gene for layer 2*, TRABD2A* (ACD, Cat No. 532881) as a marker gene for layer 5, and *MBP* (ACD, Cat No. 411051) as a marker gene for white matter.

#### 4.1.3 Polygene risk score (PRS) calculation

Genotype samples used in this study were genotyped as part of larger multiple Illumina microarray batches. Before imputation, standard quality control (QC) was performed separately within each batch to remove variants with evidence of Hardy-Weinberg equilibrium (HWE) deviation (p < 1 × 10⁻5), low minor allele frequency (MAF < 0.05) or missingness rate higher than 5%. Imputation was then performed separately for each batch using the NHLBI TOPMed Imputation Server with the hg38 reference panel. After imputation, variants with imputation quality R² > 0.9 were retained from each batch and merged into a single study-level dataset. Final variant-level QC was then applied to retain variants with missingness <0.1, HWE p > 1 × 10⁻⁶, and MAF >0.01.

For the donors in this study, we computed polygene risk scores (PRS) using the standard linkage disequilibrium (LD)-pruning and *p*-value thresholding approach implemented in `PLINK2` (Chang et al. 2015). Single-nucleotide polymorphisms (SNPs) were pruned using *p*-value–informed clumping with an LD threshold of *r*^2^ = 0.1 within a 1,000-kb window. For each discovery GWAS, 4 sets of PRS were generated from the pruned SNP sets using the following association *p*-value thresholds: 1e-08, 1e-07, 1e-06, 1e-05. Specifically, we generated separate PRS using summary statistics from the most recent PGC bipolar disorder GWAS (GWAS_BD_) (O’Connell et al. 2025) and the PGC major depressive disorder GWAS excluding 23andMe (GWAS_MDD_) (Adams et al. 2025), both of which were based on European ancestry samples (**SF1A-B**). PRS weights were defined as the natural logarithm of the odds ratios from the corresponding GWAS. We assessed whether individual genome-aggregate risk for MDD or BD predicted diagnosis in our cohort with binary logistic regression models (`dx ∼ PRS`), including the top 5 genomic principal components, biological sex, and donor age as model covariates to adjust for (Choi et al. 2020). For each GWAS risk set (GWAS_MDD_ and GWAS_BD_) and each GWAS association threshold, we examined the effect of PRS on the predicted probability of MDD diagnosis (N=79, 40 NTC/ 39 MDD), of BD diagnosis (N=80, 40 NTC/ 40 BD), and of either mood disorder diagnosis (N=119, 40 NTC/ 79 mood disorder). We used the Benjamini & Hochberg false discovery rate (FDR) procedure (Benjamini and Hochberg 1995) to adjust for an inflated Type I error due to testing multiple GWAS association thresholds. PRS was considered to be significantly associated with diagnosis with an adjusted *p*-value < 0.05.

### 4.2 Spatially-resolved transcriptomics (SRT) data collection

#### 4.2.1 Visium H&E data generation

Following anatomical confirmation, tissue blocks were acclimated inside the cryostat (Leica CM3050s) at −14°C for 30 minutes. Each issue block was mounted onto a chuck using Optical Temperature Compound (TissueTek Sakura, Cat #4583). Approximately 30-50µm of tissue was trimmed to achieve a flat surface, and tissue blocks were scored with a razor blade (ULINE, Cat #H-595B) if the surface was larger than 6.5 × 6.5 mm in dimension to fit the most representative area within the sample on a 6.5 × 6.5 mm array. Given that each Visium slide contained four capture areas, to balance across the cohort, we included two diagnostic groups on each slide, one male and one female, such that each slide contained either MDD/BD, MDD/NTC, or NTC/BD.

Hematoxylin and Eosin (H&E) staining was performed according to manufacturer’s protocols using manufacturer-recommended third-party products (10x Genomics, CG000160 Rev C). High-resolution brightfield images were acquired using Leica Aperio CS2 slide scanner equipped with a 20x/0.75NA objective and a 2x doubler. cDNA synthesis and library generation were performed according to manufacturer’s protocols (10x Genomics, CG000239 Rev F) as previously described (Huuki-Myers et al. 2024; Kwon et al. 2026; Thompson et al. 2025). Samples were sequenced at a minimum depth of 60,000 reads per spots at either the Johns Hopkins Single Cell Transcriptomics Core (12 samples, S4 NovaSeq 6000 (Illumina)), the Johns Hopkins Sidney Kimmel Comprehensive Cancer Center (8 samples, S4 NovaSeq 6000), or at Psomagen (Rockville, MD) (100 samples, 25B Novaseq X).

#### 4.2.2 Image preprocessing

Whole slide images were saved as raw TIFF files. Slide images were split into individual per-sample images using `VistoSeg` (Tippani et al. 2023). Images were aligned with the Loupe Browser 8 tool from 10x Genomics. The resulting JSON output file from the `Loupe Browser`, along with the segmented slide images, and FASTQ files for associated samples were used as inputs for `SpaceRanger`.

#### 4.2.3 Visium raw data processing

FASTQ and image data were pre-processed with 10x Genomics `SpaceRanger` (v2.1.0). Reads were aligned to reference genome GRCh38 2020-A. Unless otherwise noted, R version 4.4.3 was used for data analysis (R Core Team 2025). Outputs from the `SpaceRanger` pipeline were loaded into R and stored in a `SpatialExperiment` (v1.20.0) (Righelli et al. 2022) object using the `spatialLIBD::read10xVisiumWrapper` function (v1.3.16) (Pardo et al. 2022).

### 4.3 SRT pre-processing

#### 4.3.1 Spatial transcriptomics quality control

Spots identified as off-tissue by `SpaceRanger` and spots with no UMI counts were removed from analysis, leaving 548,579 spots across 119 tissue sections. We identified areas of low quality spots using `scuttle::isOutlier` (v1.16.0) (D. J. McCarthy et al. 2017), applying a raster transformation to isolate spatially contiguous regions, and characterizing these regions based on absolute size (number of spots) and the proportion of highly compromised spots (library size <=100 UMI). This approach identified edge artifacts (**SF2A**) and other outlier regions with >50% highly compromised spots (**SF2B**) and preserved biologically meaningful regions with lower expression like uncompromised L1 and WM regions (**SF2C**).

After removing edge artifacts (n=2420), other large outlier regions with >50% highly compromised spots (n=7266), and the remaining highly compromised spots (n=2904), we used `SpotSweeper` (v1.3.3) (Totty et al. 2025) to identify local outlier spots (n=741). In total, we removed 13,331 spots, retaining 97.6% of in-tissue spots (n=535,248 spots) which were balanced across sex and diagnosis groups (**SF2D-E**). Genes with fewer than 5 UMI counts were removed, leaving 28,965 genes.

#### 4.3.2 Spatially variable gene selection

For feature selection, we used `nnSVG` (v1.10.3) (Weber et al. 2023) to detect spatially variable genes (SVGs) in each donor and then merged these results to create a unified set of SVGs that were representative of all donors. Prior to SVG detection, gene expression was normalized and then log_2_-transformed based on spot-level scale factors computed from library size using `scuttle`. We pre-filtered to 6111 genes with at least 3 counts in at least 0.1% spots across all donors. Following `nnSVG` guidelines, the dataset was then split by donor and, for each donor, genes were then filtered to those expressed in at least 100 spots (6059.92 ± 161.79 genes, mean ± SD across all donors). After `nnSVG` was run for each donor, 2 genes were not expressed in >10% spots in any sample and 3 genes were not significant (adj. *p*<.05) in any `nnSVG` model. From the remaining 6106 genes we filtered out those that produced a significant result in less than 10 models (567 genes) and then genes that produced a significant result in less than one-third of the models for which it was included (1273 genes). To isolate candidate SVGs representing the strongest spatial expression patterns, we then removed any gene that was not ranked in the top 500 genes in at least one model in which it was significant (2604 genes). These filters produced a final set of 1663 SVGs.

### 4.4 Spot-level annotation

We generated two types of domains, one based on spatial clustering reflecting traditional dlPFC cytoarchitectural layers (domain annotation spatial clustering, domain_SP_) and another driven by snRNA-seq cell type expression profiles (domain annotation cell type-driven, domain_CT_)

#### 4.4.1 Spatial clustering domain annotations

We used `PRECAST` (v1.6.5) (W. Liu et al. 2023) to perform spatially-aware unsupervised clustering as this method intrinsically performed batch correction and integration across the 119 donor samples. `PRECAST` required the variance stabilizing transformation via `Seurat::SCTransform` (v5.3.0) (Hafemeister and Satija 2019) to normalize the raw counts of our feature list (n=1663 SVGs). We chose *k*=9 clusters based on repeated success at this resolution in the dlPFC (**SF3A**).

`PRECAST` clusters were annotated based on marker genes from a previous SRT dlPFC study (spatialDLPFC) (Huuki-Myers et al. 2024), obtained from the `spatialLIBD`::`fetch_data` function for `“spatialDLPFC_Visium_modeling_results“` (Pardo et al. 2022). We extracted the union of the top 100 *t*-statistics from each of the 9 spatialDLPFC domains, encompassing each of the traditional L1-L6 dlPFC layers, one domain consistent with the meninges (Mng), and two WM domains. In addition, we guaranteed the presence of canonical markers of inhibitory neurons (*SLC6A1*, *SLC32A1, GAD1*, *GAD2*, *SST*, *CCK*, *CNR1*, *TAC1*, *PVALB*, *RELN*), resulting in a list of 757 markers genes (with 147 genes in the top 100 *t*-statistics for multiple spatialDLPFC domains). This gene set provided preliminary annotations for 8 of our 9 clusters as menginges-enriched (Mng), GABA-enriched, L1, L2, L3/4, L5, L6, white matter (WM) (**SF3B**). We performed spatial smoothing to prioritize spatially contiguous laminar annotations, during which 62% of Mng spots were reassigned (n=4487 spots remaining) and 95.1% of GABA spots were reassigned (n=341 spots remaining) (**SF3C**). The remaining GABA and Vasc spots were dropped from the final spatial clustering domain annotations (domain_SP_).

After smoothing, there was still one cluster that was not enriched for dlPFC marker genes and exhibited decreased library size (**SF3D**). Silhouette analysis revealed little homogeneity within the low UMI cluster when pseudobulked (**SF3E**). The low UMI cluster was dropped from the final spatial clustering domain annotations (domain_SP_) (**SF3F**). In addition to the 13,526 spots removed during QC, this cluster-level QC amounted to the exclusion of only 7.5% of all in-tissue spots. The 508,372 retained spots were distributed across the following domains_SP_ and equally distributed across diagnosis-by-sex groups: 12.7% L1, 19.7% L2, 28.3% L3/4, 14.3% L5, 17.3% L6, and 7.7% WM (**SF3G**).

Domains_SP_ were pseudobulked by aggregating raw gene expression counts across spots within each domain for each donor using the `scuttle::aggregateAcrossCells`. We calculated QC metrics for the pseudobulk samples and removed samples with fewer than 7,500 detected genes, improving homogeneity (**SF3H-I**). Our final domains_SP_ pseudobulk dataset consists of 690 samples, with the top two PCs well integrated for diagnosis and sex and clearly separating the annotated dlPFC domains (**Fig 1K**).

#### 4.4.2 Cell type-driven domain annotations

##### Preparation of PFC snRNAseq for cell-type label projection

PsychENCODE v5 snRNA-seq PFC data (SZBDMulti-seq, BA10) (Ruzicka et al. 2024) was obtained via Synapse data repository (syn22755055). The SZBDMulti-seq dataset was selected from several PFC and dlPFC datasets (Emani et al. 2024) due to the large number of individuals sampled, the high quality of the snRNA-seq data, and the presence of neurotypical control donors and BD donors for validation of label projection accuracy between diagnosis groups. `scanpy` (1.10.2) (Wolf et al. 2018) in a `Python` (3.10.14) environment was used for QC since the data were provided as `adata` files. `Seurat` (v5.3.0) (Hao et al. 2024) was used for preprocessing and label transfer.

We removed a few snRNA-seq samples with outlier QC metrics, retaining 169,536 control nuclei (from 10 female and 11 male donors) and 131,598 BD nuclei (from 11 female and 11 male donors). The control and BD nuclei were processed separately with the standard `Seurat` pipeline which utilizes `SCTransform` counts to generate PCs and nearest neighbors based on 2000 highly variable genes.

Although the SZBDMulti-seq dataset contained Azimuth annotations consistent across all PsychENCODE V5 datasets, we performed graph-based clustering on control nuclei to obtain lower resolution cell type labels for projection into our SRT data. The control nuclei were distributed across the following annotated clusters: 1.1% microglia and vascular (M/V), 3.9% astrocytes (Ast), 11.9% oligodendrocytes (Olg), 19.2% inhibitory neurons (Inb), and the following excitatory neuron types: 10.5% L2, 18.2% L3, 10.4% L4, 11.8% L5, 12.9% L6 (**Figure 1F** (top)).

##### Cell-type label projection with Seurat

We used `Seurat::FindTransferAnchors` and `Seurat::TransferData` to project cell type labels from SZBDMulti-seq control nuclei into a query dataset. For each query dataset (SZBDMulti-seq BD nuclei and our SRT spots) the raw counts matrix was normalized with variance-stabilizing transformation and PCA was performed based on 2000 highly variable genes (SZBDMulti-seq BD nuclei) or 1663 SVGs (our SRT spots). Briefly, pairs of reference-query observations (nuclei or spots) were identified as anchors based on *k* = 50 nearest neighbors calculated from 30 PCs. Anchor sets were then used to transfer cell type annotation labels from the reference to the query dataset. For the SZBDMulti-seq BD query data, 30 PCs were used for weight reduction during transfer, and for the MBv SRT query data we used the 15 `PRECAST` embeddings for weight reduction.

The SZBDMulti-seq BD query dataset provided a basis for evaluating the fidelity of our label projection approach given the presence of Azimuth nuclei annotation provided for both control and BD nuclei. Using these native Azimuth labels as a “ground truth”, we observed strong concordance between control-projected Azimuth labels and native BD Azimuth labels (Rand index= 0.8986) (**SF4A**). Projecting the lower resolution clusters from control nuclei onto BD nuclei increased prediction scores for the sparsely sampled vascular and immune cell types (**SF4B**).

Projecting lower resolution clusters from control nuclei onto our SRT data produced a very small number of L2 annotated spots so L2 and L3 annotations were merged (L2/3) (**SF4C**). This cell type-driven approach allowed for low UMI spots to be rescued; these spots were distributed across different cell type labels but were predominantly annotated as Olg and M/V which fit with both the anatomical regions where most low UMI cluster spots were found and with the reduced number of RNA transcripts routinely collected from these cell types (**SF4D**). The 535,248 spots were distributed across the following cell type-driven domain annotations (domain_CT_): 4.6% M/V, 21.5% Ast, 18.6% L2/3, 7.7% L4, 9.9% Inb, 15.0% L5, 15.1% L6, 7.7% Olg. As with the domain_SP_ annotations, we found domains_CT_ to be equally represented across diagnosis-by-sex groups (**SF4E**).

Domains_CT_ were pseudobulked and we removed samples with fewer than 8,000 detected genes as well as three outlier M/V samples with fewer than 10,000 detected genes (**SF4F**). The filtered pseudobulk samples exhibited adequate homogeneity within domains (**SF4G**). The domain_CT_ pseudobulk dataset consisted of 942 samples, with the top two domain_CT_ PCs highly explained by cell type biology and integrated across diagnosis and sex (**Fig 1L**).

#### 4.4.3 Validating SRT annotation with domain-enrichment modeling

We performed spatial registration of enriched marker genes to validate the domain_SP_ and domain_CT_ annotations. Enrichment was evaluated via one-vs-all differential expression testing with `limma` (Ritchie et al. 2015) to obtain statistically enriched genes for both domains_SP_ and domains_CT_. In this model, we utilized covariates of PC3 (batch effect), diagnosis, sex, and the number of spots per pseudobulk sample. We used the `limma::duplicateCorrelation` function to estimate within-donor patterns (Smyth et al. 2005). We used `limma::eBayes` to compute the moderated *t*-test for each domain_SP_ and each domain_CT_. For each comparison we selected the 100 genes with the highest *t*-statistics (n= 592 domain_SP_ marker genes, n= 765 domain_CT_ marker genes) to cross-reference with previous dlPFC studies.

We validated our marker genes against the SZBDMulti-seq control nuclei and the spatialDLPFC dataset. The enrichment results from the spatialDLPFC dataset were obtained as described in **Methods 4.4.1**. For the SZBDMulti-seq control nuclei, we computed enrichment model results similarly to our own SRT data. Following QC, snRNA-seq data were pseudobulked by donor and cluster. Pseudobulk samples with fewer than 5000 detected genes were removed, resulting in a final snRNA-seq pseudobulk dataset of 188 samples. The one-vs-all differential expression model between snRNA-seq clusters included covariates for sex and the number of nuclei per pseudobulk sample and we used the `limma::duplicateCorrelation` function to estimate within-donor patterns.

Spatial registration of marker genes was accomplished by calculating the Pearson correlation coefficient of the moderated *t*-statistics from the external datasets and our SRT annotations. Spatial registration was performed separately for domain_SP_ and domain_CT_ based on their respective marker gene sets (n= 592 domain_SP_ marker genes, n= 765 domain_CT_ marker genes) (**Figure 1E,G**).

### 4.5 Differential expression (DE) analysis

#### 4.5.1 Model overview

We performed pseudobulk differential expression (DE) analysis utilizing the `limma` RNA-seq framework (Ritchie et al. 2015). We used `edgeR::filterByExpr` function to remove genes with low expression across donors and dlPFC domains (Yunshun Chen et al. 2016). For our DE models we used `voom` weights to account for the mean-variance relationship in log-normalized count data (Law et al. 2014) and we used the `limma::duplicateCorrelation` function to estimate the correlation between the different pseudobulk observations for each domain within each donor (Smyth et al. 2005). DE models were fit with additional covariates using `lmFit` (see **Methods 4.5.4**).

#### 4.5.2 Model design

We defined our primary groups of interest based on the interaction between diagnosis and sex variables (diagnosis-by-sex, `dx-sex`) using the group-means parametrization and we formally evaluated the interaction using `limma::makeContrasts` function. We chose this interaction approach to test the influence of sex as a biological variable on gene expression in mood disorders, rather than adjust for differences in sex by including it as a covariate. Also, this approach allowed us to test sex-specific outcomes through contrast matrix design while still fitting the model with all 119 donors. Specifically, we were interested in 6 contrasts that tested the sex-specific impact of each diagnostic group: NTC females vs MDD females (F_NTC.MDD), NTC females vs BD females (F_NTC.BPD), MDD females vs BD females (F_MDD.BPD), NTC males vs MDD males (M_NTC.MDD), NTC males vs BD males (M_NTC.BPD), MDD males vs BD males (M_MDD.BPD).

We considered two different sets of DE models to test for different hypotheses. The first model adjusted for differences in gene expression due to domain annotation to examine the impact of diagnosis-by-sex groups across the whole dlPFC section (`∼ dx-sex + domain`, **Figure 2A**). The second model allowed for a domain-restricted approach by examining the interaction between diagnosis-by-sex groups and domain on gene expression (`∼ dx-sex-domain`, **Figure 2C**). The second model allowed for post-hoc comparisons of the diagnosis-by-sex groups to be restricted within each domain (e.g. L1 NTC female samples vs L1 MDD female samples tested with a L1_F_NTC.MDD contrast).

#### 4.5.3 Hypothesis testing

For each gene, we performed omnibus *F*-tests (to test for differences in means across all groups) and *t*-tests (to test for differences between two groups as a single contrast) to identify differentially expressed genes. Statistical inference was based on empirical Bayes moderated *F*-statistics and *t*-statistics using `limma::eBayes` (Smyth 2004), which borrows information across genes to stabilize residual variance estimates, improve power, and protect against hypo- and hyper-variable genes common in RNA-seq data (Phipson et al. 2016). We used the Benjamini & Hochberg false discovery rate (FDR) procedure (Benjamini and Hochberg 1995) to adjust for an inflated Type I error due to performing an *F*-test for each gene. We calculated the FDR adjusted *p*-value of the moderated *t*-test based on the total number of tests, reflecting the repeated testing of each gene for each contrast (adjustment size= # genes * # contrasts; varied between annotations and models). This strict adjustment strategy is referred to as the “global” adjustment method in the `limma` user guide and is recommended when a set of closely related contrasts are being tested and when a goal is to interpret the number of DE genes as indicative of the strength of the contrast.

In order to be considered a differentially expressed gene (DEG) for a given model, we required a significant diagnosis-by-sex group effect (*F*-test adj. p<.05) and a significant contrast (*t*-test adj. *p*<.05).

#### 4.5.4 Covariate selection

We considered common donor-level covariates available for our cohort including age (**Figure 1B**), body mass index (BMI) calculated at autopsy, and RNA integrity number (RIN) calculated at time of dissection from cortical tissue (**SF6A**), and found these continuous measurements to not be collinear (age-BMI rho= −0.0729, age-RIN rho= 0.0516, BMI-RIN rho= −0.1660). In our dataset, smoking status (as reported by next-of-kin) was confounded with diagnosis for both males and females (◻-squared = 29.096, df = 2, *p* = 4.807e-07) (**SF6A**). We found no direct effects of diagnosis-by-sex group on the continuous donor-level covariates of age (F_5,113_=0.1855, *p*=0.9676), BMI (F_5,113_=0.2004, *p*=0.9617), and RIN (F_5,113_=1.773, *p*=0.1241).

Pseudobulk sample-level contributors to the top PCs with `scater::varianceExplainedPCs` demonstrated that biological factors accounted for most of the sample-level variance in the first two PCs (**SF6B**). Predictably, the metrics of library size, number of detected genes, and number of spots were highly collinear (domain_SP_: min rho between any pair= 0.711; domain_CT_: min rho between any pair=0.726). Given that library size is accounted for during `limma-voom` DE modeling, we selected the number of spots per pseudobulk sample as a potential covariate to represent this group of variables.

We found that PC3 represented sample-level variation largely attributable to technical elements of experimental design (donor/capture area, Visium slide, and sequencing batch) (**SF6B**). Gene-level variance of potential covariates assessed by `scater::getVarianceExplained` also showed strong correlation between PC3 and these technical and experimental factors (**SF6C**). To include Visium slide as a covariate in the DE model would result in 30 individual binary coefficients being estimated (one per slide), therefore we opted to utilize PC3 as a continuous representation of technical batch effects in our DE models.

We examined the following covariates for potential inclusion in our final DE model: PC3, number of spots per pseudobulk sample (nspots), mitochondrial fraction (chrM ratio), age, BMI, RIN, and smoking status. Among the continuous covariates, the maximum absolute correlation found was between chrM ratio and RIN (domain_SP_ rho= −0.196, domain_CT_ rho= −0.236). We used `scater::getVarianceExplained` to estimate the amount of gene-level variance attributed to each of the covariates within each of the domains_SP_ and domains_CT_ (**SF6D**). We found that PC3 explained a large amount of variance for the greatest number of genes, and that the amount of variance in gene expression explained by PC3 and nspots varied between domains, especially WM_SP_/Olg_CT_.

Covariate sensitivity was examined with both the whole-tissue model (`∼ dx-sex + domain`) and the domain-restricted model (`∼ dx-sex-domain`). All continuous covariates were scaled and centered (*z*-score) prior to inclusion. We tested the DE models with no additional covariates (“none”, `basic model`), the DE models with PC3 as the lone additional covariate (“pc3 only”, `basic model + pc3`), and the DE models plus PC3 and each of the following covariates in a one-at-a-time fashion (basic model + pc3 + *c*). Sensitivity of `limma` DE results to the different covariate models was evaluated by examining the number of DEGs (*F*-test adj. *p*<.05 and *t*-test adj. *p*<.05). Including PC3 as a covariate doubled the number of DEGs compared to the `basic model` (**SF6E**). Largely, the number of DEGs and the identity of the DEGs were robust to different covariates, aside from an increase in the RIN covariate model (**SF6F-G**).

Based on these analyses, our final models for DE include PC3, age, and nspots as covariates. The final whole-tissue model was `∼ dx-sex + domain + PC3 + age + nspots`. The final domain-restricted model was `∼ dx-sex-domain + PC3 + age + nspots`.

### 4.6 Cell type characterization of DEGs

To identify the cell types in which our DEGs are typically expressed, we again referenced the SZBDMulti-seq control snRNA-seq data (**Methods 4.4.2**). We collapsed the Aziumuth annotations to very broad cell types of vascular (Vasc), microglia and immune cells (Micro), oligodendrocyte lineage (Oligo), astrocytes (Astro), excitatory neurons (ExcN), and inhibitory neurons (InhN). We processed the single-nuclei raw counts with `scater::logNormCounts`, and then calculated the mean expression (logcounts) and proportion of nuclei with detected expression for all genes expressed in >100 nuclei (**SF10A-C**).

We implemented the *MeanRatio* approach (Huuki-Myers et al. 2025) to characterize DEGs into discrete groups based on the dominant-expressing cell type (**SF10D**). *MeanRatio* identifies the dominant-expressing cell type based on the ratio of cell type-averaged expression for the highest and second highest expressing cell types. For our implementation, we required that the gene be expressed in at least 10% of nuclei within the highest expressing cell type prior to calculating the *MeanRatio*. Of the 503 total unique DEGs, 410 genes qualified for *MeanRatio* cell typing. Genes were considered to have a dominant-expressing cell type with a mean ratio > 1.5. We classified genes that did not reach a *MeanRatio* threshold of 1.5 as “multi” due to the relatively equal expression in at least two of the examined cell types (e.g. SLC6A1, see **SF10**). This classification approach identified the following number of DEGs to be predominantly expressed in these broad cell types: Vasc = 95, Micro = 47, Astro = 25, Oligo = 25, ExcN = 18, InhN = 20, multi = 180. This approach helped us to identify commonalities between DEGs that were not evident through other analysis, such as the presence of InhN-dominant DEGs in supragranular layers (**Figure 2E**, **SF10E**).

### 4.7 Spot-level gene expression analysis

#### 4.7.1 Gene expression modules with GRNBOOST2

We used `GRNBOOST2` to identify modules of co-expressed DEGs within the `arboreto` (v0.1.6) framework (Moerman et al. 2019). `GRNBOOST2` is a machine learning method (based on gradient-boosted decision trees to capture non-linear relationships) to find gene-gene (predictor-target) relationships. For each predictor-target pair, it produces an importance score. Here, we provided a custom set of 817 features (any gene with a significant omnibus *F*-test in any model and any annotation) as predictors. This analysis uses spot-level resolution expression input, so we utilized the domain_SP_ annotations to remove spots that were annotated to the low UMI spatial cluster (n=513,200 spots). We performed gene-level quality control by using `edgeR::filterByExpr` to filter to genes with minimum expression by donor and domain_CT_. The domain_CT_ annotations were used for gene-level quality control to make sure that genes robustly expressed in the more specialized populations like M/V_CT_ and Inb_CT_ were included in co-expression analysis. A total of 13,162 genes passed the minimum expression filter, including 500 of the original 503 DEGs.

`GRNBOOST2` importance scores were generated using the raw counts matrix, with gene-gene Pearson correlations subsequently used to identify the directionality of the predictor-target relationship. We utilized the normalized, log_2_-transformed gene expression counts (zeros were removed) to calculate Pearson rho and increased the threshold for positive relationship from the default rho > 0.03 to rho > 0.20 due to the enhanced correlation expected from spot-level data. Co-expression modules were obtained by retaining predictor-target pairs with a minimum importance score of 1 and enforcing a minimum module size of at least 20 genes (19 target genes plus predictor gene). Finally, we isolated the DEG components of co-expression modules by removing non-DEG targets and enforcing a minimum of 10 DEG targets present per predictor module. This resulted in a set of 104 initial DEG modules that contained 485 of the possible 500 DEGs, many of which exhibited strong correlations with one another (**SF11A**).

To extract a subset of minimally overlapping DEG modules representative of the greatest possible number of DEGs, we isolated the top predictor module for each DEG target and constructed a network of the top predictors and all 485 DEG targets (**SF11B**). Examining the network centrality for each subnetwork of top predictors with at least 6 elements, we selected 19 top predictors modules that were highly connected with target DEGs (462 DEG targets). We re-evaluated these 19 modules for correlation of target gene importance and eliminated two based on a high similarity (*EIF1* removed due to Pearson rho with *EEF1A1*= 0.49, *UQCRH* removed due to Pearson rho with *COX4I1*= 0.63), leaving 448 DEG targets (**SF11C**). Next, these 17 DEG modules were refined so that target genes with larger importance in one module than others had the less important relationships pruned (**SF11D**). After refining, two modules (*JUNB* and *ADAMTS1*) contained fewer than 10 DEGs, which was an initial exclusion criteria. Therefore, the refining step was re-run for 15 module DEGs (excluding *JUNB* and *ADAMTS1*) to incorporate *JUNB* and *ADATMS1* target DEGs into any other qualifying module. The final set of 15 DEG modules (*SNHG14*, *PRKAR1A*, *COX4I1*, *EEF1A1*, *PLP1, GFAP*, *GLUL*, *HSPA1A*, *MT1M*, *IFITM3*, *CD74*, *A2M*, *GRIN1*, *CAMK2N1*, *GAD1*) contained 446 DEGs (**Figure 2F, SF11E-F**), achieving our goal of producing minimally overlapping DEG modules that still retained 92.0% of the DEGs present in the initial 104 DEG modules.

The contribution of modules to individual spots was quantified using `AUCell` (Aibar et al. 2017) which calculates whether the module DEGs were enriched in each spot by evaluating the top 5% of genes (658 genes) expressed in each spot (**SF12**).

#### 4.7.2 SCENIC analysis

The `SCENIC` (v0.12.1+11.g06bafba) pipeline for regulon analysis starts by implementing `GRNBOOST2` to build a co-expression network of 1892 human transcription factors (Aibar et al. 2017). We used the default threshold of rho > .03 to filter to positive associations, and filtered to the top 20% of targets for each transcription factor before applying a blanket cutoff selecting for targets with importance >0.5. Using pre-computed hg38 transcription factor motif rankings based on promoter-proximal (500bp upstream and 100bp downstream, “promoter” annotation set) and promoter-distal (10kbp upstream and 10kbp downstream, “enhancer” annotation set) genomic regions of target genes, `cisTarget` (v0.2.0) then determines whether a motif feature was enriched for the set of transcription factor targets. Motifs were matched to the corresponding transcription factor source (motif similarity FDR= 1e-5 and orthologous identity threshold of 0.8). Final regulons include the merged enhancer-enriched and promoter-enriched sets for all directly annotated motifs. `AUCell` was used to calculate spot-level regulon enrichment.

Regulon analysis with 513,200 spots from 119 donors found three AP1 family regulons that exhibited unusually high AUCell values in the same 10 donors (**SF15A**). Looking at the targets with the highest importance weights revealed the transcription factor *NPAS4* as highly co-expressed in the FOSB- and JUNB-regulons. We confirmed that the 10 donors with the highest average of these 3 AP1-regulons had abnormally high *NPAS4* expression and were not concentrated in any diagnosis or sex group (**SF15B-C**). *NPAS4* expression is neuron-specific and is exclusively expressed in a short window (hours) following neuronal activity, implicating peri-mortem factors as a source for this expression pattern. Thus, we excluded these samples and performed regulon analysis with N=109 donors (**SF15D**, **SF16**).

### 4.8 smFISH validation

To validate cell-type identities and transcript expression patterns observed at spot-based resolution at a single-cell level, we ran smFISH using the RNAScope Multiplex Fluorescent Reagent Kit v2 (Advanced Cell Diagnostics [ACD], Cat No. 323100) and 4-Plex Ancillary Kit (ACD, Cat No. 323120), as previously described (Maynard et al. 2021), in N=4 (2M/2F) NTC donors with representative anatomy and tissue quality metrics (Br6432, Br5558, Br1113, Br8667). Several 10 μm dlPFC cryosections for each donor were collected on a Leica cryostat to utilize across two experimental panels. Panel one examined the distribution of TAC1+ neurons across the dlPFC layers, using *PCP4* as boundary of superficial vs. deep cortical layers. Probes targeted *SLC17A7* (assigned to Opal dye 520), *TAC1* (assigned to Opal dye 690), *GAD1* (assigned to Opal 570), and *PCP4* (assigned to Opal dye 620) with catalog numbers 415611, 310711, 404031, and 446111, respectively. Z-stacked images were acquired using identical acquisition settings across all donors, collecting tiles from L1-WM with a Nikon APO lambda D 20x / 0.80 NA objective. Within Nikon NIS-Elements software v6.20.02, tiled z-stack images were stitched and projected as maximum-intensity projection images. Panel two aimed to validate the identity of a *GAD1*/*TRBC2*-double positive cell-type rather than *TRBC2/AIF1* colocalization. Probes targeted *TRBC2* (assigned opal dye 690), *GAD1* (assigned Opal dye 520), and *AIF1* (assigned opal 570), using catalogue numbers 433451, 404031, and 433121, respectively. In this experiment, single inset images were acquired using identical settings across donors with a Nikon APO lambda D 40x / 0.9 NA objective. All images across both panels were spectrally unmixed within the Nikon NIS-Elements software using optimized linear unmixing to isolate and remove lipofuscin autofluorescence and to deconvolve overlapping signal from the shared 570/620 laser track.

The tiled, unmixed image files were analyzed using HALO software v2.4.4 (Indica Labs) as previously described (Shah et al. 2026; Mulvey et al. 2026). Cell phenotyping was restricted to detection of *PCP4* and *TAC1* transcript expression based on standardized HALO analysis settings available on GitHub (spatialDLPFC_mdd_bpd/processed-data/15_RNAscope_Validation). Image orientation was manually reviewed to ensure consistent anatomical directionality of L1–WM and alignment of L1 to a common reference axis. Within HALO, regions of interest (ROIs) from L1 to WM were defined for each section. To account for differences in tissue width, cortical depth was standardized across sections by computing a relative position metric (x_rel), where 0 and 1 correspond to the L1 and WM boundaries. Cell centroids, derived from X minimum and maximum boundaries of DAPI+ nuclei, were used to assign spatial coordinates along x_rel. For visualization of layer-specific transcript abundance, *PCP4+* and *TAC1+* single-positive populations were analyzed using Gaussian kernel density estimation (KDE) implemented in ggplot2::geom_density. Density was plotted along the normalized cortical axis.

### 4.9 Spatial eQTL analysis and colocalization analysis

#### 4.9.1 eQTL analysis in domain_CT_ annotations

To identify genetic variants associated with cell type-driven domain-specific gene expression, we performed *cis* expression quantitative trait loci (eQTL) mapping using donor-level pseudobulk expression profiles for each domain_CT_, after removing spots annotated to the low UMI spatial domain (n=513,200 spots). Genes on the Y chromosome and from the mitochondrial genome were excluded, and genes were retained for eQTL testing if they were expressed with a minimum threshold of counts per million (CPM) >= 0.10 and raw counts >= 6 in at least 20% of donors within a domain_CT_ group (on average 13,185 genes were retained, 8,377 to 15,386 range). Pseudobulk expression was represented as Trimmed Mean of M-values (TMM)-normalized log_2_CPM values with a prior count of 1 using `edgeR` (Robinson et al. 2010), followed by rank-based inverse normal transformed across donors following GTEx’s eQTL mapping practice (THE GTEX CONSORTIUM 2020).

We performed cis-eQTL mapping with `tensorQTL` (v1.0.10) (Taylor-Weiner et al. 2019), testing variants with minor allele frequency >= 0.05 within 1 Mb of each gene transcription start site. The all-donor linear model adjusted for diagnosis, sex, donor age, PC3, number of spots contributing to each pseudobulk, the first five genotype principal components (PCs), and domain_CT_-specific expression PCs. Expression PCs were calculated from TMM-normalized *z*-scored expression values using the top 5,000 highly variable genes, while the number of expression PCs was estimated separately for each domain_CT_ group using `sva::num.sv` (v3.58.0) (Leek et al. 2012), yielding 14-23 expression PCs across domain_CT_ groups. Gene-level cis-eQTL significance was estimated from permutation-based `map_cis` *q*-values at FDR < 0.05. Conditionally independent cis-eQTL signals were identified with `tensorQTL`’s `map_independent`, restricted to genes significant in `map_cis` (15,002 eQTL signals significant with both `map_cis` and `map_independent`). For downstream colocalization analysis, nominal variant-gene associations were generated with `map_nominal`. Final curated cis-independent, nominal (filtered to FDR<.05), summary, and gene-intersection outputs are provided in **ST6**.

#### 4.9.2 Colocalization with SCZ, MDD, and BD GWAS loci

To evaluate whether domain_CT_ cis-eQTL signals and loci associated with mood disorders in genome wide association studies (GWAS) were consistent with a shared causal variant, we performed Bayesian colocalization using `coloc.abf` from `coloc` (v5.2.3) (Giambartolomei et al. 2014). Colocalization was performed separately using the latest European-ancestry GWAS summary statistics from the Psychiatric Genomics Consortium: SCZ (Trubetskoy et al. 2022), MDD (Adams et al. 2025) and BD (O’Connell et al. 2025) (**SF20A**). For each domain_CT_ and gene locus, we assembled nominal eQTLs and GWAS summary statistics for variants in the cis-eQTL testing window and retained loci with at least 10 overlapping variants for colocalization. After sensitivity testing using `coloc::sensitivity`, we summarized posterior probabilities for the standard `coloc` hypotheses, focusing on strong support for a single shared causal variant between gene expression and GWAS association (`PP.H4` >= 0.8). `PP.H4` (PP4) is the posterior probability for `H4` in `coloc.abf`, the hypothesis of a single shared causal variant in that locus. Gene loci with strong colocalization (PP4 > 0.8) are presented in **ST7** and instances where the exact colocalization variant was also a cis-independent eQTL result (*q* < .05) or a nominal eQTL result (FDR < .05) are labeled.

#### 4.9.3 Overlap of eQTLs and colocalization results with DEG and GWAS data

Cis-independent eQTLs and colocalization results were examined for overlaps between the associated gene and the 503 DEGs identified in this study (**Figure 6C**, **ST6**, **ST7**). We summarized overlap between cis-independent eQTLs and colocalization results with psychiatric GWAS evidence using GWAS annotation thresholds. Variant-level GWAS support was defined by an exact match between an eQTL variant and a GWAS risk variant (*p* < 5e-08) in the corresponding disorder set (**SF20E**). Additionally, we implemented an exploratory threshold for mood disorder GWAS variants (*p* < 1e-05), which increased the overlaps between GWAS_MDD_-GWAS_BD_ and GWAS_BD_-GWAS_SCZ_ and revealed overlap with DEG-associated eQTL and colocalization results (**SF20F-G**). Variants identified by either cis-independent eQTL analysis or colocalization analysis that overlapped with both DEG support and GWAS variant-level support are presented in **SF20**. Gene-level GWAS support was defined by overlap between the eGene or colocalized gene and curated GWAS risk-gene lists. Although gene-level GWAS support identified no additional DEG-associated cis-independent eQTL or colocalization results, instances of gene-level GWAS support are labeled in **ST6** and **ST7**, along with columns indicating exact-variant GWAS support.

### 4.10 Data and Code Availability and Visualization

Project data is available via the National Institute of Mental Health Data Archive (NDA) and can be found with study ID 3235. Image and SpaceRanger output data are available through Gene Expression Omnibus (GEO) under accession GSE333974. The code for this project is publicly available through GitHub at https://github.com/LieberInstitute/spatialDLPFC_mdd_bpd, which is archived on Zenodo (jac-thom et al. 2026). Data objects used for analysis are available on Globus (`jhpce#spatialDLPFC_mdd_bpd` at https://research.libd.org/globus/) along with a `README.txt` file with a list of files and descriptions. Two code-free companion visualization tools are available at https://research.libd.org/spatialDLPFC_mdd_bpd/ and can be accessed through the project GitHub. These tools include visualization of pseudobulk data gene expression and DE results for all donors, with a Samui Browser-powered SRT visualization for 24 donors (split evenly between diagnosis and sex) (Sriworarat et al. 2023).

## 4.11 Acknowledgements, Funding, Authorship Contributions

We gratefully thank the families who donated tissue to make this research possible. We thank the Essel Foundation and the families of Connie and Stephen Lieber and Milton and Tamar Maltz for their generous support of this work. We also would like to acknowledge Dr. Fernando Goes, the late Dr. Llewellyn B. Bigelow, Anna Brandtjen, and James Tooke for their contributions to this work in sample curation and clinical characterization. We thank the Office of the Chief Medical Examiner of the State of Maryland, the Department of Pathology at Western Michigan University Homer Stryker MD School of Medicine, the University of North Dakota School of Medicine and Health Sciences, Department of Pathology, the County of Santa Clara Medical Examiner-Coroner Office, Gift of Life Michigan, and the NIMH Intramural Research Program for their collaboration on tissue collection. We gratefully acknowledge use of the facilities at the Joint High Performance Computing Exchange (JHPCE) in the Department of Biostatistics, Johns Hopkins Bloomberg School of Public Health that have contributed to the research results reported within this paper. We thank members of the Maynard, Martinowich, Hicks, and Page labs for their feedback on the manuscript in preparation.

## Author Contributions

Conceived and designed the study: KM, SCH, SCP

Performed experiments and collected data: SO, SEM, SVB, BJM, MRA, SHK, MRV, YD, ABN, SCP

Software: RAM, MT

Formal analysis: JRT, GP, MT, SH

Data curation: JRT, GP, RAM, MT, AD-S, LCT

Tissue Resources: AD-S, JEK, TMH

Writing-Original Draft: JRT, KM, SCH, SCP

Writing-Review and editing: JRT, SEM, SVB, BJM, GP, SH, LCT, KM, SCH, SCP

Visualization: JRT, GP, MRA, SHK

Supervision: TMH, KRM, LCT, KM, SCH, SCP

Funding Acquisition: KM

Project Administration: KM, SCH, SCP

