## Supplemental Figures & Legends, Supplemental Table Legends for "Overlapping yet distinct spatiomolecular profiles across mood disorders and biological sex in the human prefrontal cortex"

### Supplementary Figures

**Supplemental Figure 1. Polygenic risk for mood disorders is not strongly predictive of diagnosis in our cohort of 119 donors.**

(**A**) Multiple polygenic risk scores for MDD ($PRS_{MDD}$, *y*-axis, centered and scaled) were calculated from GWAS of MDD ($GWAS_{MDD}$) using different association *p*-value thresholds (facets). Cohort donors are delineated by diagnosis (*x*-axis, color) and biological sex (point shape).

(**B**) Multiple polygenic risk scores for BD ($PRS_{BD}$, *y*-axis, centered and scaled) were calculated from GWAS of BD ($GWAS_{BD}$) using different association *p*-value thresholds (facets). Cohort donors are delineated by diagnosis (*x*-axis, color) and biological sex (point shape).

(**C**) Logistic regression models tested whether $PRS_{MDD}$ (left) or $PRS_{BD}$ (right) predicted diagnosis (DX, *y*-axis) in our cohort. Results are present for models where DX = MDD (79 total donors, NTC = 40 donors), DX = BD (80 total donors, NTC = 40 donors), and DX = MDD or BD (119 total donors, NTC = 40 donors). Model results are presented as odds ratio (OR, *x*-axis) with range indicating 95% confidence interval (CI) and a dashed vertical line at OR = 1 (equal odds). Point color indicates the GWAS association threshold used to calculate PRS in the tested model. Point outline and CI line color indicates whether the result was significant at $p < .05$ after adjusting for multiple threshold tests.

(**D**) Results of models predicting diagnosis from $PRS_{BD}$ (calculated with $GWAS_{BD}$ association $p <$ 1e-07) are present for each mood disorder diagnosis tested (DX, facets). The adjusted *p*-value for predicted association is labeled for each model. The *x*-axis reflects centered and scaled $PRS_{BD}$. The fitted values of the model for each donor (point) are plotted on the *y*-axis, representing probability of predicted diagnosis (NTC=0, DX=1) with values of 0.5 indicating equal predicted likelihood for either diagnosis. Point color indicates the true diagnosis of donors in our cohort.

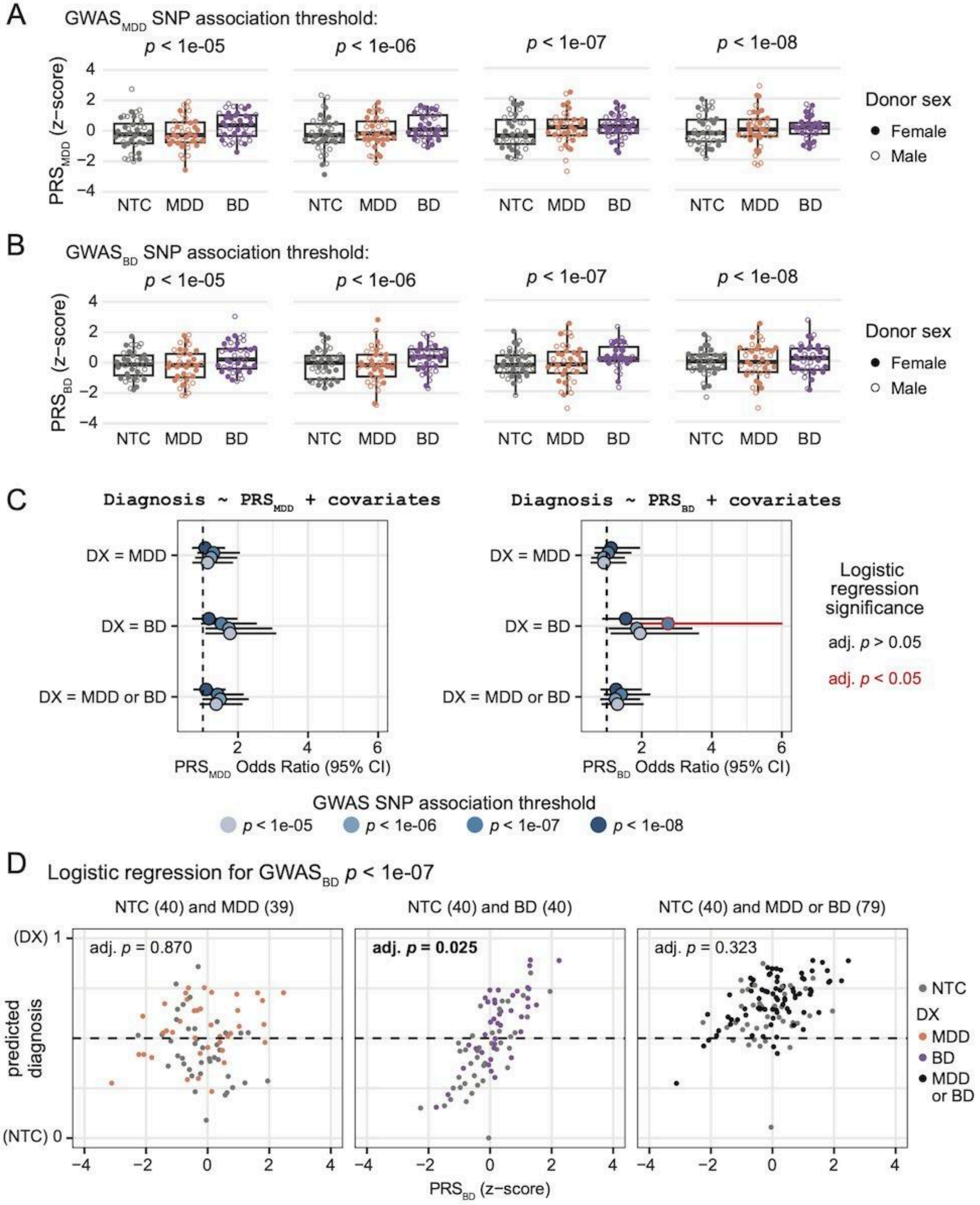
A
GWAS_MDD SNP association threshold:
p < 1e-05
p < 1e-06
p < 1e-07
p < 1e-08
PRS_MDD (z-score)
NTC MDD BD
Donor sex
Female
Male
B
GWAS_BD SNP association threshold:
PRS_BD (z-score)
C
Diagnosis ~ PRS_MDD + covariates
Diagnosis ~ PRS_BD + covariates
DX = MDD
DX = BD
DX = MDD or BD
PRS_MDD Odds Ratio (95% CI)
PRS_BD Odds Ratio (95% CI)
Logistic regression significance
adj. p > 0.05
adj. p < 0.05
GWAS SNP association threshold
p < 1e-05
p < 1e-06
p < 1e-07
p < 1e-08
D
Logistic regression for GWAS_BD p < 1e-07
NTC (40) and MDD (39)
NTC (40) and BD (40)
NTC (40) and MDD or BD (79)
adj. p = 0.870
adj. p = 0.025
adj. p = 0.323
predicted diagnosis
(DX) 1
(NTC) 0
PRS_BD (z-score)
NTC
DX
MDD
BD
MDD or BD

**Supplemental Figure 2. Overview of spot-level quality control.**

(**A**) Top: Example tissue section (Br5599) before and after quality control (QC) removing an edge artifact. Colors represent the (i) total library size ($log_{10}$ scale). Bottom: spots with less than 100 UMIs (red), spots identified as local outliers as defined by `Spotsweeper` (black), and spots with contiguous regions of outliers (pink).

(**B**) Example tissue section (Br5617) before and after QC removing a problem area corresponding to low-read white matter. Colors are the same as in (**A**).

(**C**) Example tissue section (Br5137) before and after QC for a typical tissue without edge artifacts or problem area. Colors are the same as in (**A**).

(**D**) Distributions of QC metrics summarized per donor based on whether the spots were kept (grey) or discarded (red). Plots are stratified by diagnosis status (x-axis) and sex (facet column). QC metrics (y-axis) from top to bottom are library size ($log_{10}$-transformed total number of UMIs), detected genes ($log_{10}$-transformed total number of detected genes), mitochondrial fraction (proportion of reads mapping to the mitochondrial genome), and the number of discarded spots as proportion of in-tissue spots.

(**E**) Distribution number of detected genes (y-axis, $log_{10}$-transformed) across spots for each sample (x-axis) after QC filtering faceted by diagnosis status (rows) and sex (columns).

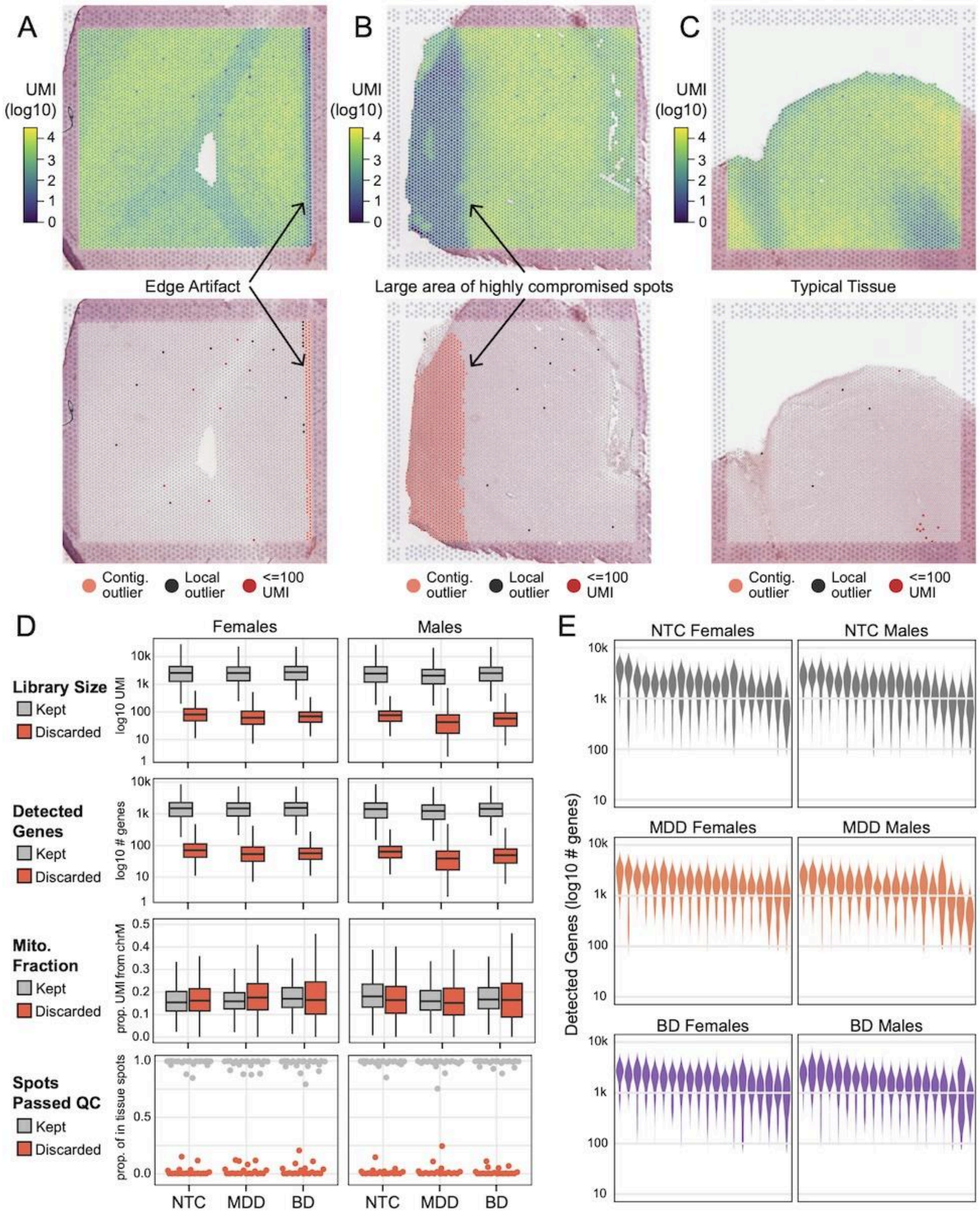

A
B
C
UMI (log10)
4
3
2
1
0
Edge Artifact
Large area of highly compromised spots
Typical Tissue
Contig. outlier
Local outlier
<=100 UMI
D
Females
Males
Library Size
Kept
Discarded
log10 UMI
10k
1k
100
10
1
Detected Genes
log10 # genes
Mito. Fraction
prop. UMI from chrM
0.5
0.4
0.3
0.2
0.1
0.0
Spots Passed QC
prop. of in tissue spots
1.0
0.5
0.0
NTC
MDD
BD
E
NTC Females
NTC Males
MDD Females
MDD Males
BD Females
BD Males
Detected Genes (log10 # genes)

**Supplemental Figure 3. Annotating SRT spots as spatial clustering domains ($\text{domain}_{SP}$) reflecting traditional dlPFC cytoarchitectural layers.**

(**A**) Spot plot of initial predicted spatial clusters (*k* = 9) in Br6432 predicted by `PRECAST`. Annotations included menginges-enriched (Mng), GABA-enriched, layers (L) L1, L2, L3/4, L5, L6, white matter (WM) domains, and a low UMI cluster.

(**B**) Heatmaps of the marker genes (*x*-axis, columns; position identical for top and bottom) used to annotate `PRECAST` *k*=9 spatial clusters. Top: Marker genes are labeled based on the spatialDLPFC (Huuki-Myers et al. 2024) domain (*y*-axis, rows) for which it was present in the top 100 most enriched genes (filter based on *t*-statistics). A small number of inhibitory neuron markers were added and these are labeled as GABAergic. Bottom: For each `PRECAST` k=9 spatial cluster (y-axis, rows), marker gene expression (logcounts) was averaged (fill color, centered and scaled), allowing for annotation labels to be derived from corresponding spatialDLPFC domains.

(**C**) Spot plot of smoothed `PRECAST` spatial domains (*k* = 7) in Br6432. After smoothing, remaining Mng and GABA spots were removed due to sparsity. See (**A**) for color legend.

(**D**) Violin plot showing the distribution of total UMI counts (*y*-axis) from pseudobulk samples across smoothed `PRECAST` L1, L2, L3/4, L5, L6, WM, and low UMI domains (*x*-axis).

(**E**) Violin plot showing the silhouette widths for pseudobulk samples across smoothed `PRECAST` L1, L2, L3/4, L5, L6, WM, and low UMI domains. Pseudobulk samples are colored by the smoothed domain with the greatest similarity (see (**A**) for color legend), with silhouette widths < 0 (horizontal black line) indicating greater similarity to a non-annotated domain.

(**F**) Spot plot of final spatial domains (*k* = 6, $\text{domains}_{SP}$) in Br6432 after spatial smoothing and removal of remaining Mng, GABA, and low UMI spots. See (**A**) for color legend.

(**G**) Bar plots of the proportion of spots (*y*-axis) assigned to each $\text{domain}_{SP}$ (fill color; L1, L2, L3/4, L5, L6, WM), faceted by diagnosis and sex groups. Each donor is represented by a single bar on the *x*-axis.

(**H**) Violin plot of distribution of number of detected genes (*y*-axis) for each pseudobulk sample stratified by $\text{domains}_{SP}$ (*x*-axis). The dotted line at 7,500 detected genes indicates the pseudobulk QC threshold for this dataset.

(**I**) Distribution of silhouette widths (*y*-axis) for $\text{domain}_{SP}$ (*x*-axis) pseudobulk samples before (left) and after (right) dropping pseudobulk samples that were below the threshold of 7,500 detected genes. These plots demonstrate how removing low UMI cluster spots improved clustering coherency (compared to (E)) and how the pseudobulk QC filter improved coherency of the remaining 690 samples.

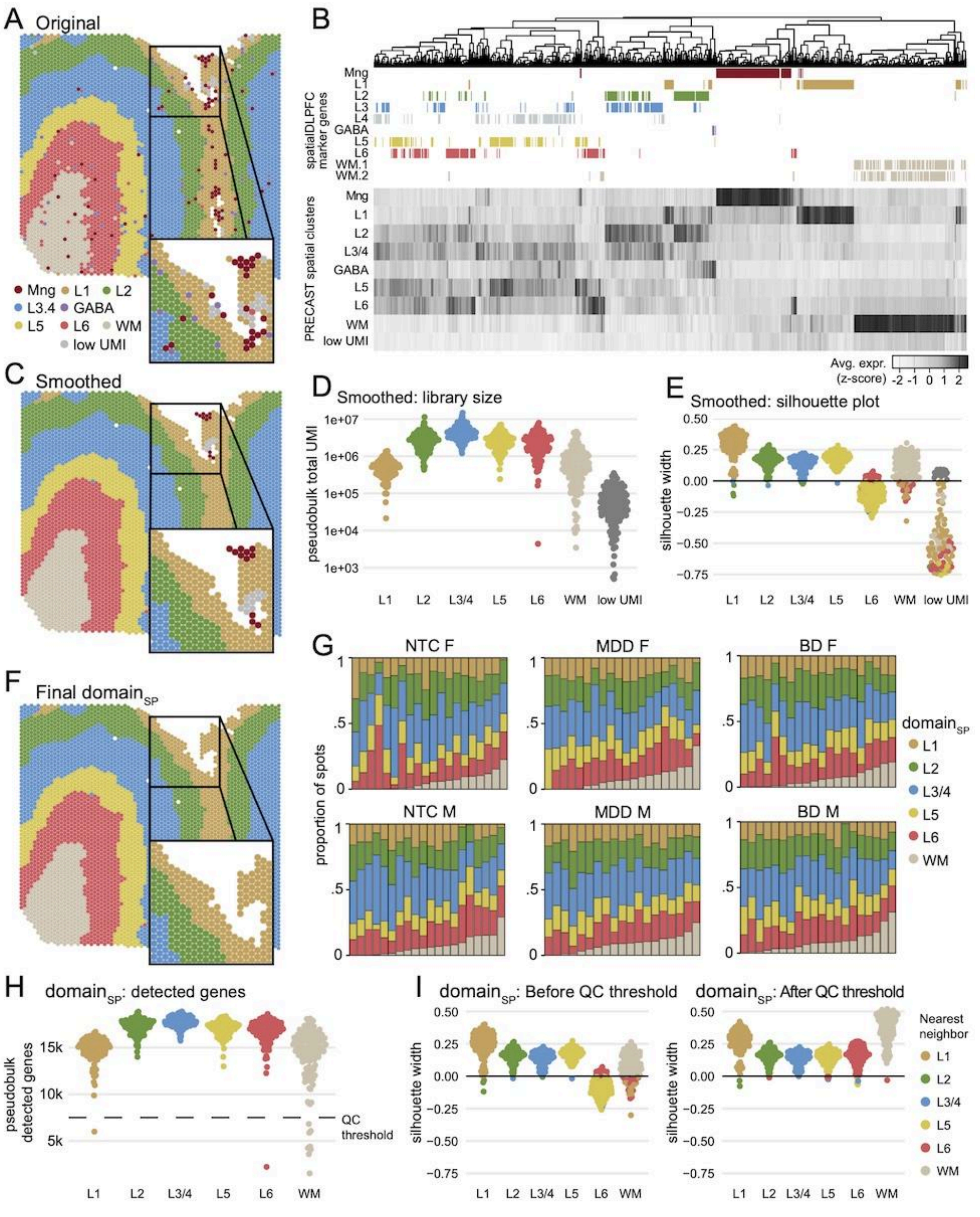

A Original
Mng L1 L2
L3.4 GABA
L5 L6 WM
low UMI
B
spatialDLPFC marker genes
Mng
L1
L2
L3
L4
GABA
L5
L6
WM.1
WM.2
PRECAST spatial clusters
Mng
L1
L2
L3/4
GABA
L5
L6
WM
low UMI
Avg. expr. (z-score) -2 -1 0 1 2
C Smoothed
D Smoothed: library size
pseudobulk total UMI
1e+07
1e+06
1e+05
1e+04
1e+03
L1 L2 L3/4 L5 L6 WM low UMI
E Smoothed: silhouette plot
silhouette width
0.50
0.25
0.00
−0.25
−0.50
−0.75
L1 L2 L3/4 L5 L6 WM low UMI
F Final domain_SP
G
NTC F
MDD F
BD F
NTC M
MDD M
BD M
proportion of spots
1
.5
0
domain_SP
L1
L2
L3/4
L5
L6
WM
H domain_SP: detected genes
pseudobulk detected genes
15k
10k
5k
QC threshold
L1 L2 L3/4 L5 L6 WM
I domain_SP: Before QC threshold
domain_SP: After QC threshold
silhouette width
0.50
0.25
0.00
−0.25
−0.50
−0.75
L1 L2 L3/4 L5 L6 WM
Nearest neighbor
L1
L2
L3/4
L5
L6
WM

**Supplemental Figure 4. Annotating SRT spots as cell type-driven domains ($domain_{CT}$) with label projection from snRNA-seq cell type expression profiles.**

(**A**) External snRNA-seq data from the PFC (Ruzicka et al. 2024) was used to test `Seurat`-based annotation projection by using the control donors as the reference and the BD donors as the query. Heatmap of congruence of the *SZBDMulti-seq* BD nuclei native Azimuth labels (*y*-axis) with the projected Azimuth cell type labels (*x*-axis) based on the Seurat prediction score (fill). High overall Rand index (0.8986) and strong preservation of annotations indicates the fidelity of our label transfer approach.

(**B**) Heatmap of congruence within *SZBDMulti-seq* BD nuclei (query dataset) testing the projection of lower resolution clusters (*x*-axis) from the *SZBDMulti-seq* control nuclei (reference dataset), in the context of the native *SZBDMulti-seq* BD Azimuth labels (*y*-axis). Lower resolution clusters are oligodendrocytes (Olg), astrocytes (Ast), microglia or immune and vascular (M/V), excitatory neurons from layers (L) 2, L3, L4, L5, L6, and inhibitory neurons (Inb). The projection of lower resolution clusters improved Seurat prediction score (fill) for sparse cell types relative to (**A**) and maintained high fidelity for more abundant cell types.

(**C**) Spot plots of example sample Br6432 after cell type projection from *SZBDMulti-seq* control nuclei into SRT data. Although reference cell types included separate labels for L2 and L3, the projected annotations were merged (L2/3) for the final $k$ = 8 final cell type annotation domains ($domain_{CT}$).

(**D**) Bar plots of the number of low UMI cluster spots (described in **SF3**) across the different $domains_{CT}$ (fill color). Each donor is represented by a single bar on the *x*-axis and plots are faceted by diagnosis and sex groups.

(**E**) Bar plots of the proportion of spots (*y*-axis) assigned to each $domain_{CT}$ (fill color; M/V, Ast, L2/3, L4, Inb, L5, L6, Olg), faceted by diagnosis and sex groups. Each donor is represented by a single bar on the *x*-axis.

(**F**) Violin plot of distribution of number of detected genes (*y*-axis) for each pseudobulk sample stratified by $domains_{CT}$ (*x*-axis). The dotted lines at 8,000 detected genes and 10,000 detected genes ($M/V_{CT}$ only) indicate the pseudobulk QC thresholds for this dataset.

(**G**) Distribution of silhouette widths (*y*-axis) for $domain_{CT}$ (*x*-axis) pseudobulk samples after dropping pseudobulk samples that were below the QC thresholds.

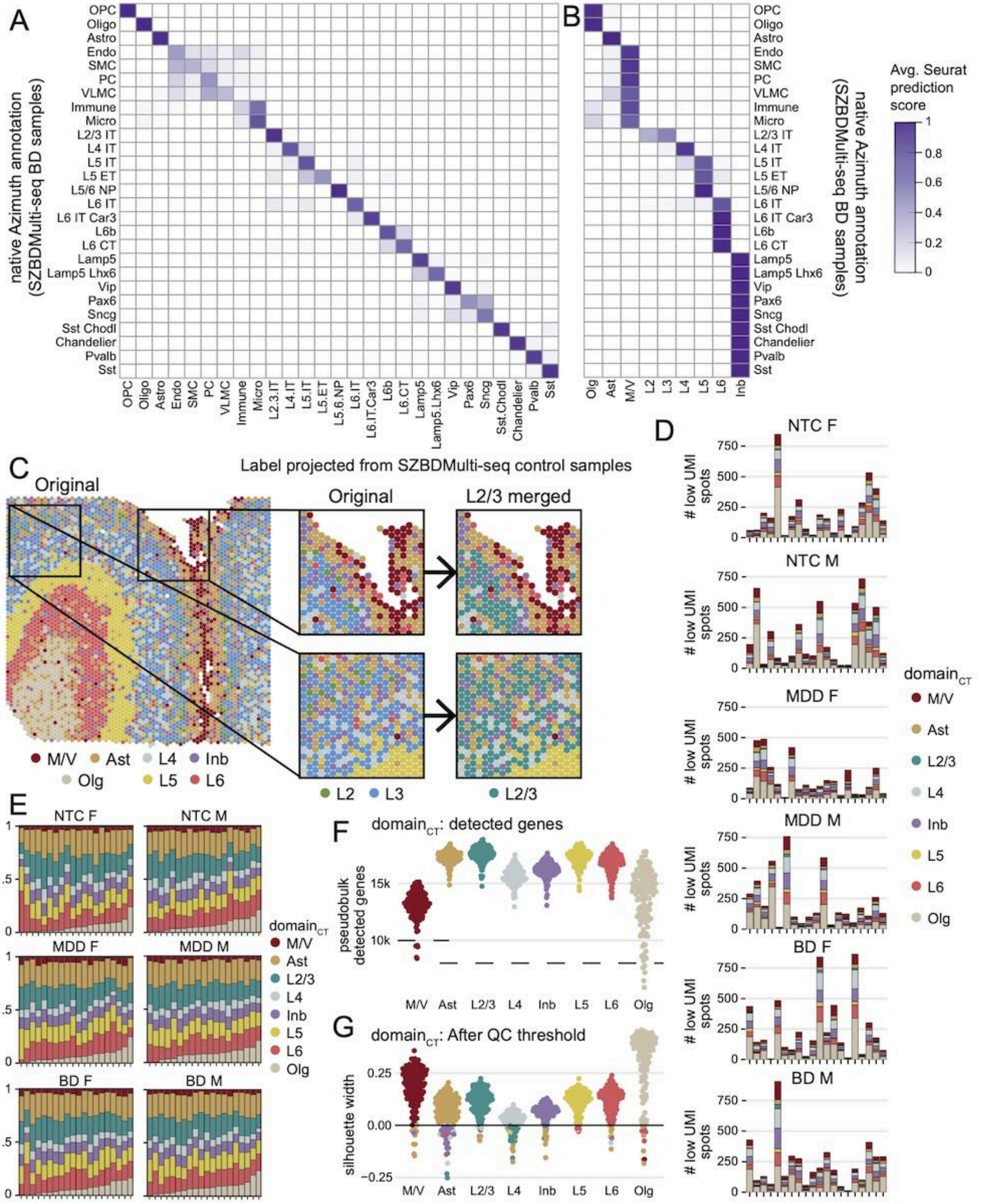

A
B
native Azimuth annotation (SZBDMulti-seq BD samples)
OPC
Oligo
Astro
Endo
SMC
PC
VLMC
Immune
Micro
L2/3 IT
L4 IT
L5 IT
L5 ET
L5/6 NP
L6 IT
L6 IT Car3
L6b
L6 CT
Lamp5
Lamp5 Lhx6
Vip
Pax6
Sncg
Sst Chodl
Chandelier
Pvalb
Sst
OPC Oligo Astro Endo SMC PC VLMC Immune Micro L2.3.IT L4.IT L5.IT L5.ET L5.6.NP L6.IT L6.IT.Car3 L6b L6.CT Lamp5 Lamp5.Lhx6 Vip Pax6 Sncg Sst.Chodl Chandelier Pvalb Sst
Olg Ast M/V L2 L3 L4 L5 L6 Inb
Avg. Seurat prediction score
1 0.8 0.6 0.4 0.2 0
C
Label projected from SZBDMulti-seq control samples
Original
Original
L2/3 merged
M/V Ast L4 Inb
Olg L5 L6
L2 L3
L2/3
D
NTC F
NTC M
MDD F
MDD M
BD F
BD M
### low UMI spots
750 500 250 0
domain_CT
M/V
Ast
L2/3
L4
Inb
L5
L6
Olg
E
NTC F
NTC M
MDD F
MDD M
BD F
BD M
1 .5 0
F
domain_CT: detected genes
pseudobulk detected genes
15k 10k
M/V Ast L2/3 L4 Inb L5 L6 Olg
G
domain_CT: After QC threshold
silhouette width
0.25 0.00 −0.25
M/V Ast L2/3 L4 Inb L5 L6 Olg

**Supplemental Figure 5. Comparison of domain$_{SP}$ and domain$_{CT}$ annotations at the spot- and gene-level.**

(**A**) Sankey plot comparing the spot-level assignment from (left to right) the (i) domain$_{SP}$ original labels, (ii) domain$_{SP}$ smoothed labels, (iii) domain$_{CT}$ merged labels, and (iv) domain$_{CT}$ original labels. The final groupings used for the domain$_{SP}$ and domain$_{CT}$ are shown in bold.

(**B**) Dot plot of select marker genes (*y*-axis) for the domain$_{SP}$ and domain$_{CT}$ annotations. Two marker genes are present for each of the following dlPFC domains or cell types (background color, top to bottom): vascular cells, microglia, astrocytes, L2, L3, L4, inhibitory neurons, L5, L6, oligodendrocytes. Spot-level logcount expression was aggregated to obtain donor- and annotation-level average expression and proportion of spots detected. Across diagnosis-by-sex groups (*x*-axis, shape), the average marker gene expression (centered and scaled) is denoted by point color and the average proportion of spots with detected marker gene expression is denoted by point size.

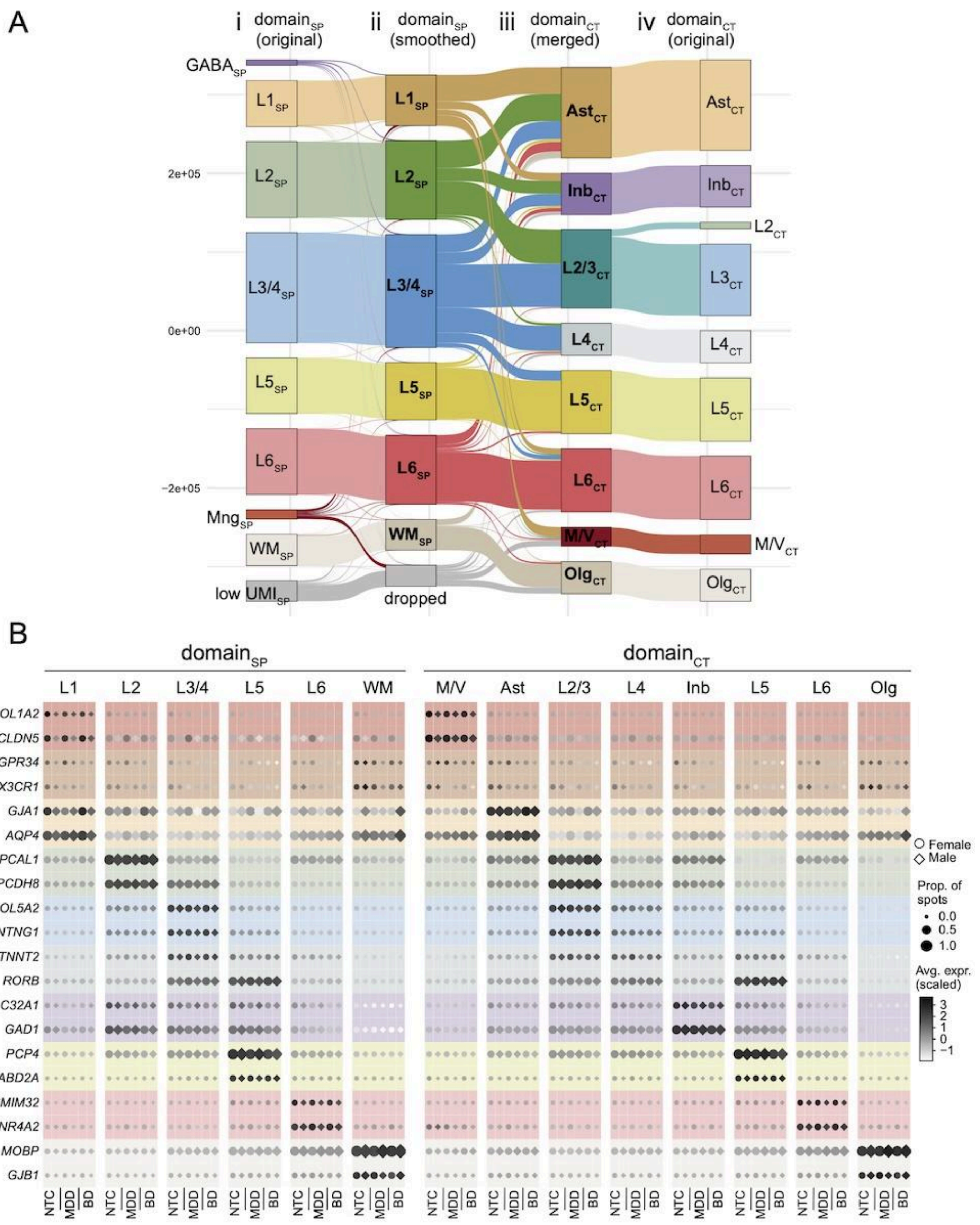
A
i domainSP (original)
ii domainSP (smoothed)
iii domainCT (merged)
iv domainCT (original)
GABASP
L1SP
L2SP
L3/4SP
L5SP
L6SP
MngSP
WMSP
low UMISP
dropped
AstCT
InbCT
L2/3CT
L4CT
L5CT
L6CT
M/VCT
OlgCT
L2CT
L3CT
2e+05
0e+00
−2e+05
B
domainSP
domainCT
L1
L2
L3/4
L5
L6
WM
M/V
Ast
L2/3
L4
Inb
Olg
COL1A2
CLDN5
GPR34
CX3CR1
GJA1
AQP4
HPCAL1
PCDH8
COL5A2
NTNG1
TNNT2
RORB
SLC32A1
GAD1
PCP4
TRABD2A
SMIM32
NR4A2
MOBP
GJB1
NTC
MDD
BD
Female
Male
Prop. of spots
0.0
0.5
1.0
Avg. expr. (scaled)
3
2
1
0
−1

**Supplemental Figure 6. Covariate selection and sensitivity analysis for differential expression modeling.**

(**A**) Distributions of donor-level variables including RNA integrity number (RIN), BMI, and smoking status. Donor-level variables are colored by diagnosis group and stratified by diagnosis and sex groups.

(**B**) Bar plots for the top 3 principal components (PCs; bar color) in the $domain_{SP}$ and $domain_{CT}$ pseudobulk datasets (facets), displaying the percent of variance for each principal component (PC) explained by spatial-biological variables (domain, number of detected genes, number of spots, sum of UMIs, chrM ratio), technical-experimental variables (donor, slide, sequencing batch), and donor-biological variables (age, smoking status, RIN, diagnosis, sex, BMI).

(**C**) For all genes in pseudobulk datasets, the proportion of gene-level variance explained by metadata variables was calculated and then correlated between variables to demonstrate the utility of PC3 as a continuous covariate for technical-experimental confounders. Color of heatmap indicates Pearson correlation coefficient (rho).

(**D**) Bar plots of the percent of gene-level variance explained by potential covariates (facet rows) was also calculated within each domain (*x*-axis, facet columns). The percent of variance explained was binned (fill color) and the number of genes in each bin was determined for each covariate and domain (*y*-axis).

(**E**) Bar plots of selected covariates that were included in differential expression (DE) models (facets) and the number of DEGs was determined (*F*-test adj. $p<.05$ and *t*-test adj. $p<.05$). Bar color indicates $domain_{SP}$ from $domain_{CT}$ results.

(**F**) The similarity between covariate models was evaluated by overlapping results (*x*-axis, number of DEGs) between sets of covariate models (*y*-axis). All models: none, pc3 only, age, nspots, BMI, chrM ratio, smoking, and RIN covariates; PC3 models: pc3 only, age, nspots, BMI, chrM ratio, smoking, and RIN covariates. Covariate model results were evaluated for each pseudobulk model design (facets, whole-tissue model and domain-restricted model) and each annotation strategy (fill, $domain_{SP}$ and $domain_{CT}$).

(**G**) The dissimilarity between covariate models was evaluated by isolating results that were unique to each covariate model. Different covariate models are present on the *y*-axis. The *x*-axis displays the number of DEGs not found in any other covariate model. Covariate model results were evaluated for each pseudobulk model design (facets, whole-tissue model and domain-restricted model) and each annotation strategy (fill, $domain_{SP}$ and $domain_{CT}$).

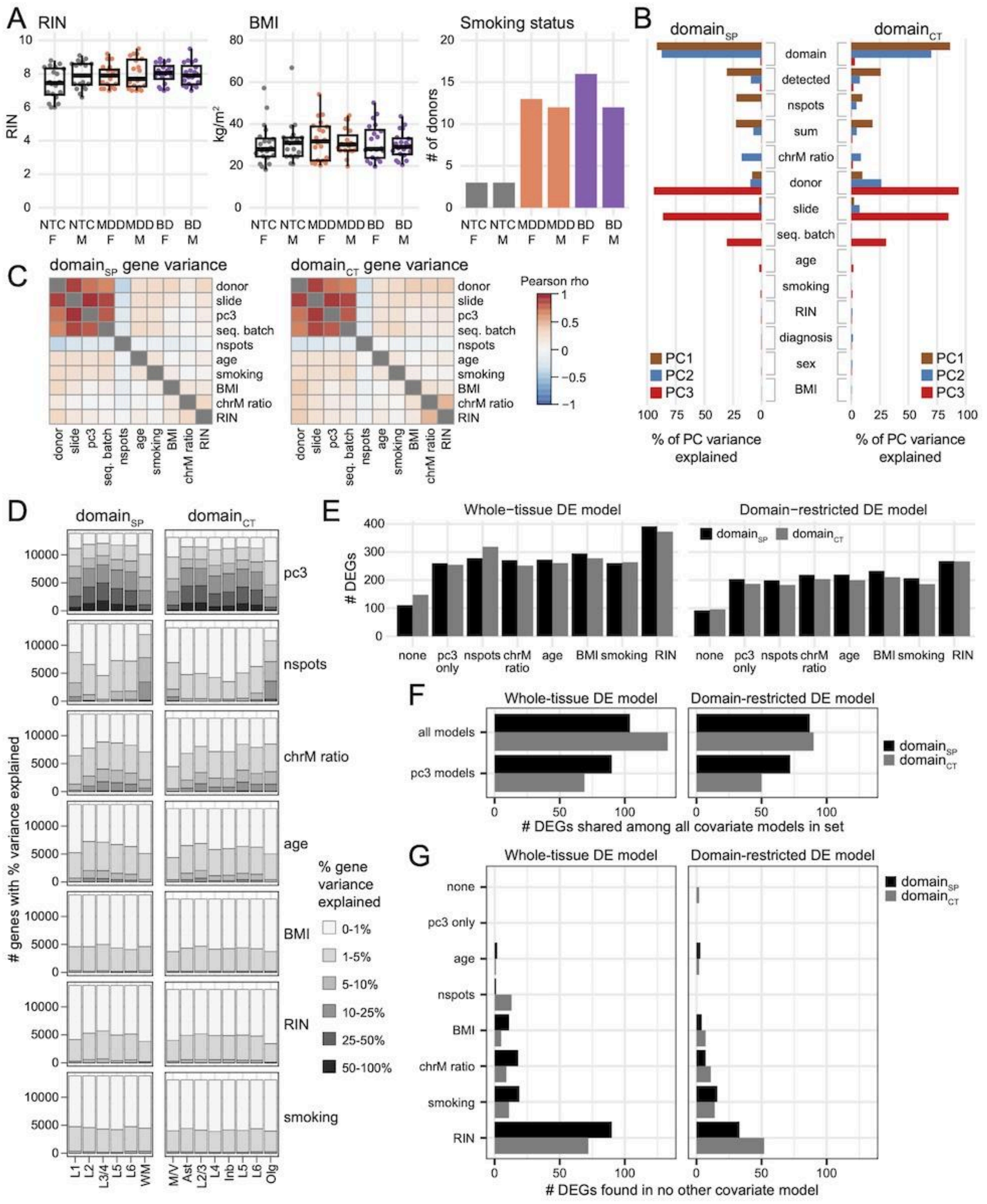

A
RIN
BMI
Smoking status
RIN
kg/m²
### of donors
NTC F
NTC M
MDD F
MDD M
BD F
BD M
B
domain_SP
domain_CT
domain
detected
nspots
sum
chrM ratio
donor
slide
seq. batch
age
smoking
RIN
diagnosis
sex
BMI
PC1
PC2
PC3
% of PC variance explained
C
domain_SP gene variance
domain_CT gene variance
donor
slide
pc3
seq. batch
nspots
age
smoking
BMI
chrM ratio
RIN
Pearson rho
D
domain_SP
domain_CT
### genes with % variance explained
pc3
nspots
chrM ratio
age
BMI
RIN
smoking
% gene variance explained
0-1%
1-5%
5-10%
10-25%
25-50%
50-100%
L1
L2
L3/4
L5
L6
WM
M/V
Ast
L2/3
L4
Inb
L5
L6
Olg
E
Whole-tissue DE model
Domain-restricted DE model
domain_SP
domain_CT
### DEGs
none
pc3 only
nspots
chrM ratio
age
BMI
smoking
RIN
F
Whole-tissue DE model
Domain-restricted DE model
all models
pc3 models
domain_SP
domain_CT
### DEGs shared among all covariate models in set
G
Whole-tissue DE model
Domain-restricted DE model
none
pc3 only
age
nspots
BMI
chrM ratio
smoking
RIN
domain_SP
domain_CT
### DEGs found in no other covariate model

**Supplemental Figure 7. Mood disorder differential expression results across and within domain$_{SP}$ annotations.**

(**A**) Volcano plots showing the domain$_{SP}$ whole-tissue model moderated *t*-test results where differentially expressed genes (DEGs) which are adjusted for domain composition. The *x*-axis is the $\log_2$ fold-change and the *y*-axis is the $-\log_{10}$-transformed FDR-adjusted *p*-value. DEGs (*F*-test adj. $p<.05$ and *t*-test adj. $p<.05$) are shown in black. Plots are faceted by each of the six diagnosis and biological sex contrasts (F_NTC.MDD, F_NTC.BD, F_MDD.BD, M_NTC.MDD, M_NTC.BD, M_MDD.BD).

(**B**) Similar to (**A**), but showing the domain-restricted model moderated *t*-test results using domain$_{SP}$ labels. Volcano plots are faceted by the six contrasts (columns) and the domain$_{SP}$ labels (rows).

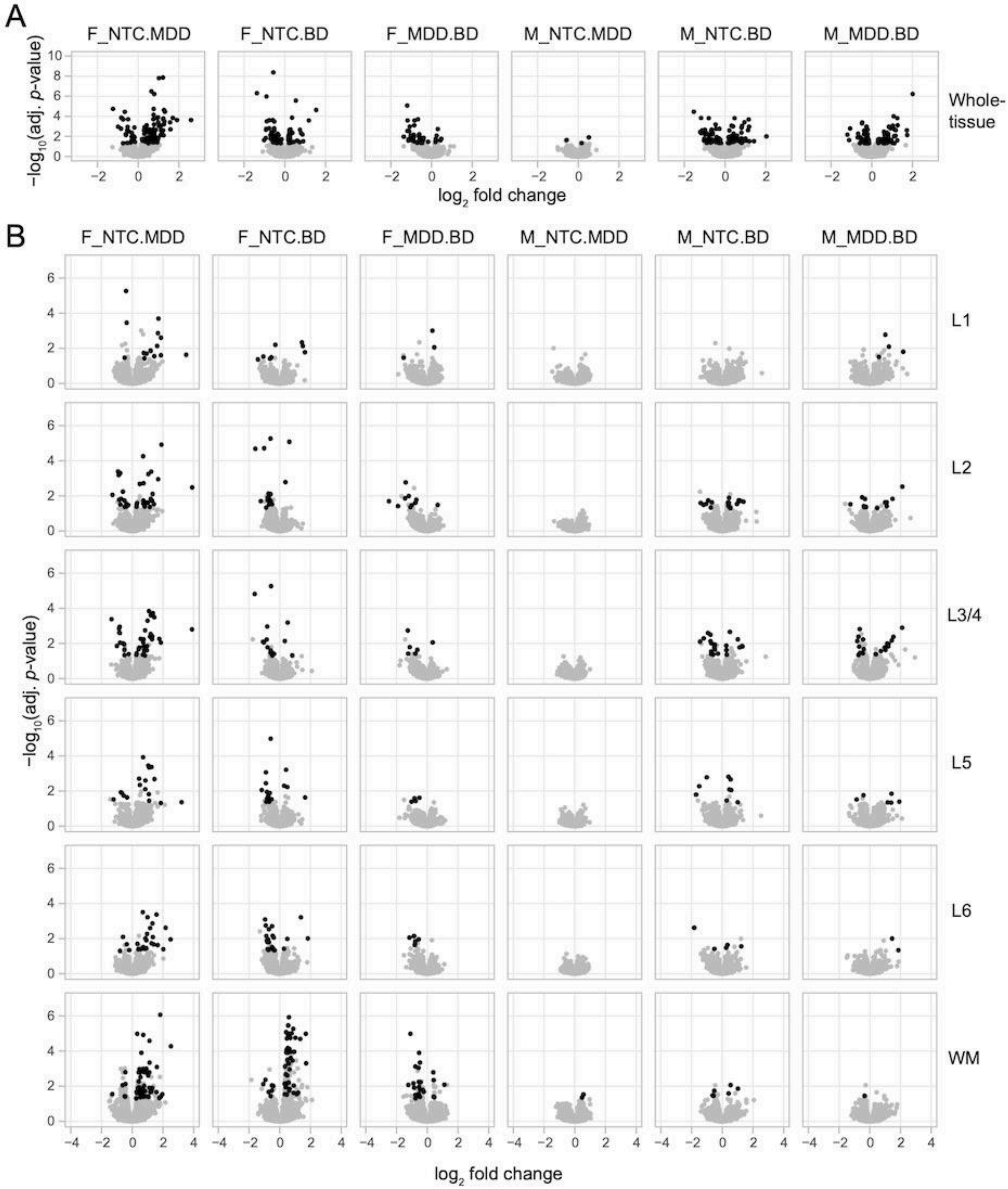
A
F_NTC.MDD
F_NTC.BD
F_MDD.BD
M_NTC.MDD
M_NTC.BD
M_MDD.BD
Whole-tissue
−log10(adj. p-value)
log2 fold change
B
F_NTC.MDD
F_NTC.BD
F_MDD.BD
M_NTC.MDD
M_NTC.BD
M_MDD.BD
L1
L2
L3/4
L5
L6
WM
−log10(adj. p-value)
log2 fold change

**Supplemental Figure 8. Mood disorder differential expression results across and within domain$_{CT}$ annotations.**

(**A**) Volcano plots showing the domain$_{CT}$ whole-tissue model moderated *t*-test results where differentially expressed genes (DEGs) which are adjusted for domain composition. The *x*-axis is the $\log_2$ fold-change and the *y*-axis is the $-\log_{10}$-transformed FDR-adjusted *p*-value. DEGs (*F*-test adj. $p<.05$ and *t*-test adj. $p<.05$) are shown in black. Plots are faceted by each of the six diagnosis and biological sex contrasts (F_NTC.MDD, F_NTC.BD, F_MDD.BD, M_NTC.MDD, M_NTC.BD, M_MDD.BD).

(**B**) Similar to (**A**), but showing the domain-restricted model moderated *t*-test results using domain$_{CT}$ labels. Volcano plots are faceted by the six contrasts (columns) and the domain$_{CT}$ labels (rows).

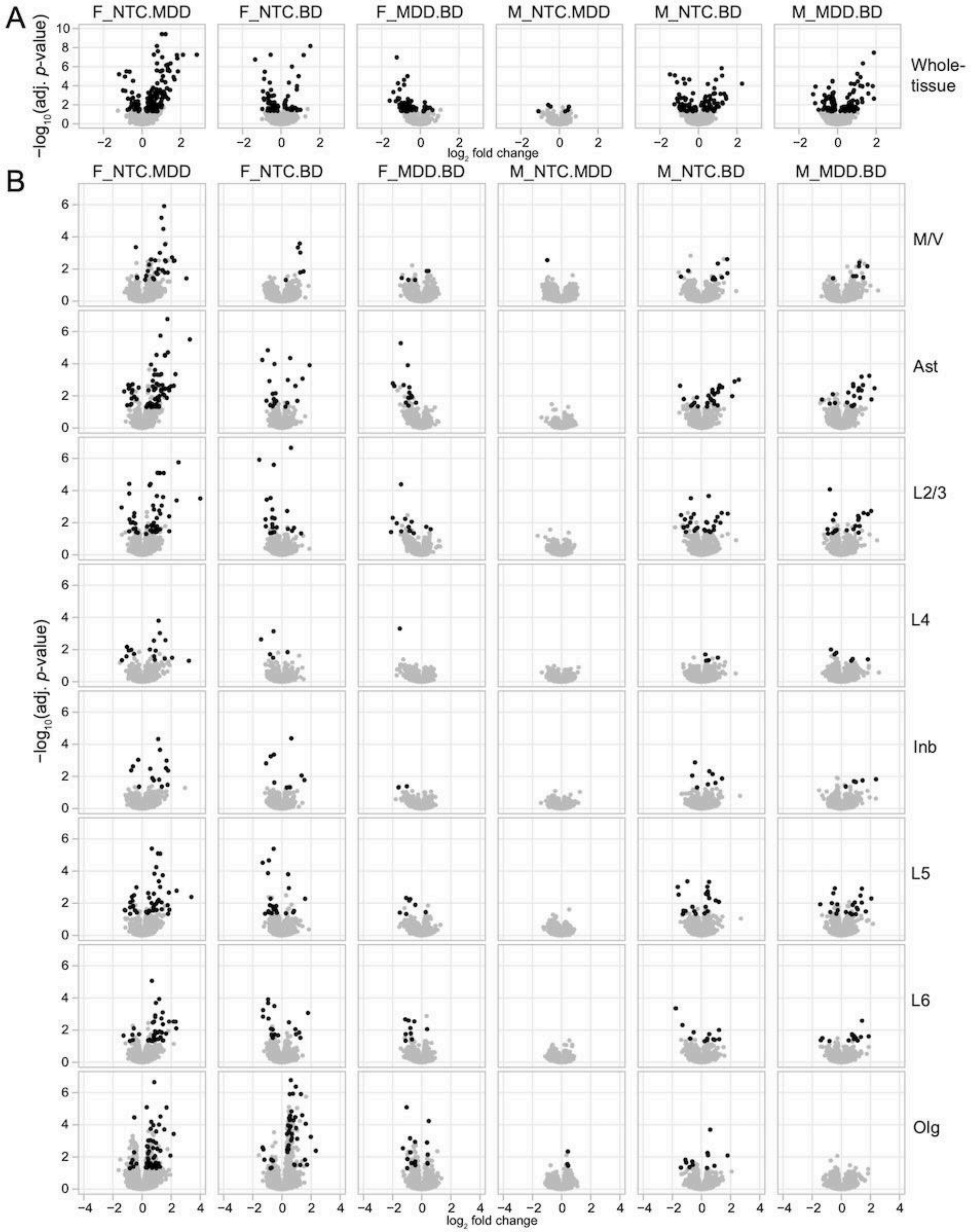
A
F_NTC.MDD
F_NTC.BD
F_MDD.BD
M_NTC.MDD
M_NTC.BD
M_MDD.BD
−log10(adj. p-value)
Whole-tissue
log2 fold change
B
M/V
Ast
L2/3
L4
Inb
L5
L6
Olg

**Supplemental Figure 9. Comparing the differential expression results between models (whole-tissue and domain-restricted) and spot annotations (domain$_{SP}$ and domain$_{CT}$).**

(**A**) Scatter plot of the whole-tissue model *F*-test results for both domain$_{SP}$ (*F*-statistic on *x*-axis) and domain$_{CT}$ (*F*-statistic on *y*-axis) results. Red lines correspond to statistical significance (*F*-test adj. $p<.05$). Pearson correlation coefficient (rho) measures agreement between *x*- and *y*-axis.

(**B**) Scatter plot of the whole-tissue model $\log_2$ fold-changes (logFC) estimated using the domain$_{SP}$ labels (*x*-axis) compared to domain$_{CT}$ (*y*-axis) labels. LogFC is calculated via contrast-specific *t*-tests; each facet corresponds to one of the six diagnosis-by-sex contrasts (F_NTC.MDD, F_NTC.BD, F_MDD.BD, M_NTC.MDD, M_NTC.BD, M_MDD.BD). Pearson correlation coefficient (rho) measures agreement between *x*- and *y*-axis.

(**C**) Similar to (**A**) comparing *F*-statistics from the omnibus *F*-tests (to test for differences in means across all groups), but now using the domain-restricted model results. Pearson correlation coefficient (rho) measures agreement between *x*- and *y*-axis.

(**D**) Similar to (**B**) comparing $\log_2$ fold-changes (logFC), but now using the domain-restricted model results. Due to the increased dimensionality of the domain-restricted test, each square in the heatmap represents a single Pearson correlation (rho, fill color) comparing the domain$_{SP}$ (*x*-axis, columns) domain-restricted logFC and the domain$_{CT}$ (*y*-axis, rows) domain-restricted logFC results for the given diagnosis-by-sex contrast (facets).

(**E**) Bar plots of the number of differentially expressed genes (DEGs) (*y*-axis, positive values indicated up-regulated DEGs, negative values indicate down-regulated DEGs) using the domain$_{SP}$ labels (*x*-axis, fill color) comparing the overlap of the whole-tissue model and domain-restricted model results (facet columns). Plots are stratified by each of the six diagnosis-by-sex contrasts (facet rows).

(**F**) Heatmap comparing the number of domain-restricted DEGs (fill color) between different domains$_{SP}$ for each diagnosis-by-sex contrast (facet row). Values on the diagonal represent DEGs that were exclusively significant within the given domain$_{SP}$ (overlapped with no other domain$_{SP}$).

(**G**) Bar plots of the number of differentially expressed genes (DEGs) (*y*-axis, positive values indicated up-regulated DEGs, negative values indicate down-regulated DEGs) using the domain$_{CT}$ labels (*x*-axis, fill color) comparing the overlap of the whole-tissue model and domain-restricted model results (facet columns). Plots are stratified by each of the six diagnosis-by-sex contrasts (facet rows). Bar color and *x*-axis position both encode domain$_{CT}$ labels, with “all” referring to whole-tissue results.

(**H**) Heatmap comparing the number of domain-restricted DEGs (fill color) between different domains$_{CT}$ for each diagnosis-by-sex contrast (facet row). Values on the diagonal represent DEGs that were exclusively significant within the given domain$_{CT}$ (overlapped with no other domain$_{CT}$).

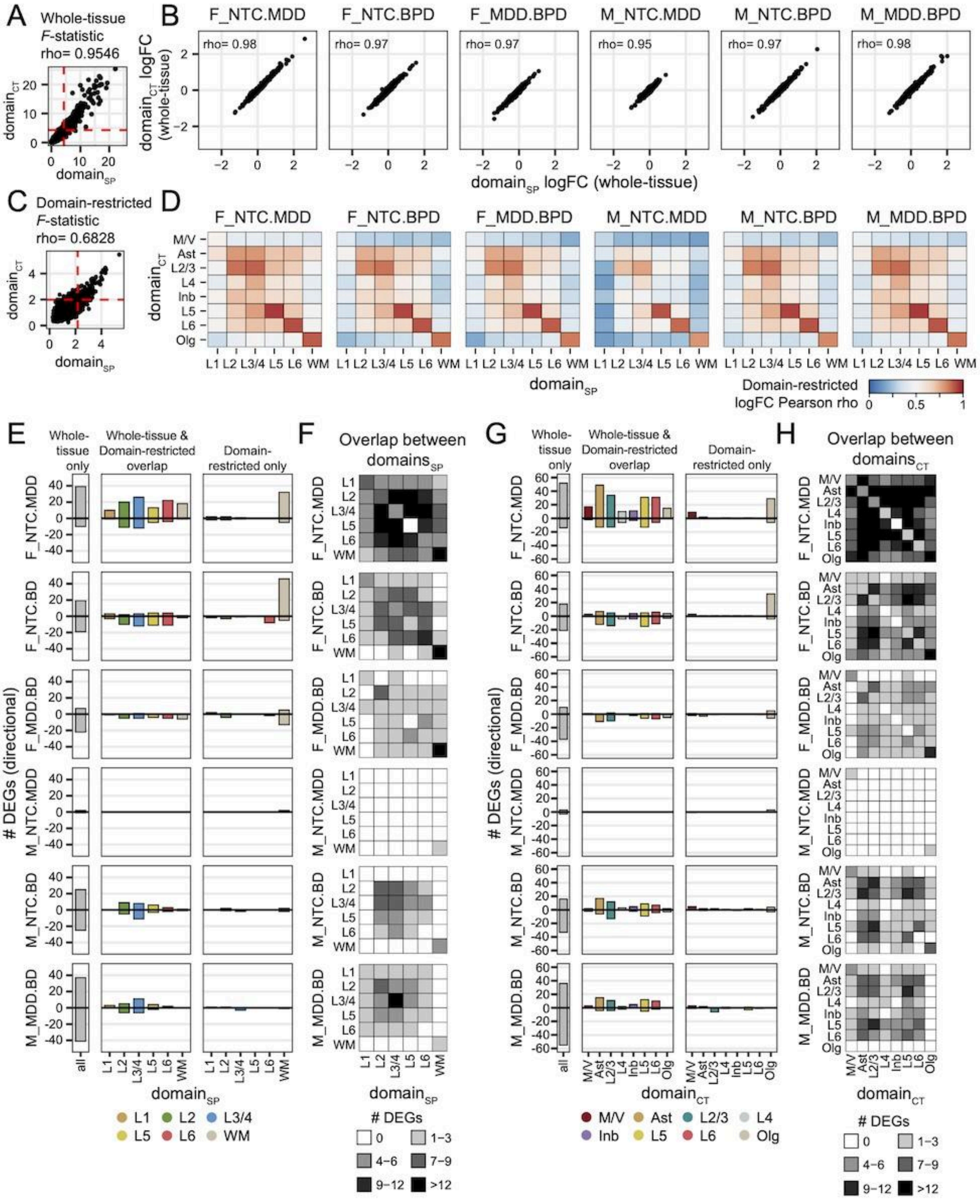

A
Whole-tissue
F-statistic
rho= 0.9546
domain_CT
domain_SP
B
F_NTC.MDD
F_NTC.BPD
F_MDD.BPD
M_NTC.MDD
M_NTC.BPD
M_MDD.BPD
rho= 0.98
rho= 0.97
rho= 0.97
rho= 0.95
rho= 0.97
rho= 0.98
domain_CT logFC (whole-tissue)
domain_SP logFC (whole-tissue)
C
Domain-restricted
F-statistic
rho= 0.6828
D
Domain-restricted logFC Pearson rho
E
Whole-tissue only
Whole-tissue & Domain-restricted overlap
Domain-restricted only
F_NTC.MDD
F_NTC.BD
F_MDD.BD
M_NTC.MDD
M_NTC.BD
M_MDD.BD
### DEGs (directional)
F
Overlap between domains_SP
G
H
Overlap between domains_CT
### DEGs
0
1-3
4-6
7-9
9-12
>12
L1 L2 L3/4 L5 L6 WM
M/V Ast L2/3 L4 Inb L5 L6 Olg

**Supplemental Figure 10. Utilizing snRNA-seq cell type expression averages to characterizing DEGs.**

(**A**) UMAP expression of SZBDMulti-seq snRNA-seq data for selected DEGs. Text indicates the proportion of nuclei expressing the DEG within the highest expressing cell type. Points (nuclei) are colored by `SCTransform` (SCT) expression values.

(**B**) Bar plots of average logcounts expression (*y*-axis) from pseudobulk of SZBDMulti-seq snRNA-seq cell types for (*x*-axis, color) of selected DEGs.

(**C**) Stacked bar plots of the average logcounts expression (*x*-axis) from pseudobulk of SZBDMulti-seq snRNA-seq cell types (fill color) selected DEGs (y-axis). Same data as (B) consolidated to single axis to reflect use throughout the manuscript.

(**D**) Schematic of *MeanRatio* approach to classify DEGs into discrete groups based on dominant cell type. Following this approach, the chosen example DEGs were classified as dominant in each of the following cell types: *ITIH5* = Vasc, *COL5A3* = Astro, *PLP1* = Oligo, *C3* = Micro, *GAD1* = InhN, *RALYL* = ExcN, *SLC6A1* = multi (not assigned to a dominant cell type).

(**E**) *MeanRatio* classification of $\text{domain}_{SP}$ DEGs for the whole-tissue model (“all”) and domain-restricted model (*x*-axis). Fill color indicates dominant cell type with *y*-axis reflecting the proportion of DEGs classified for each cell type.

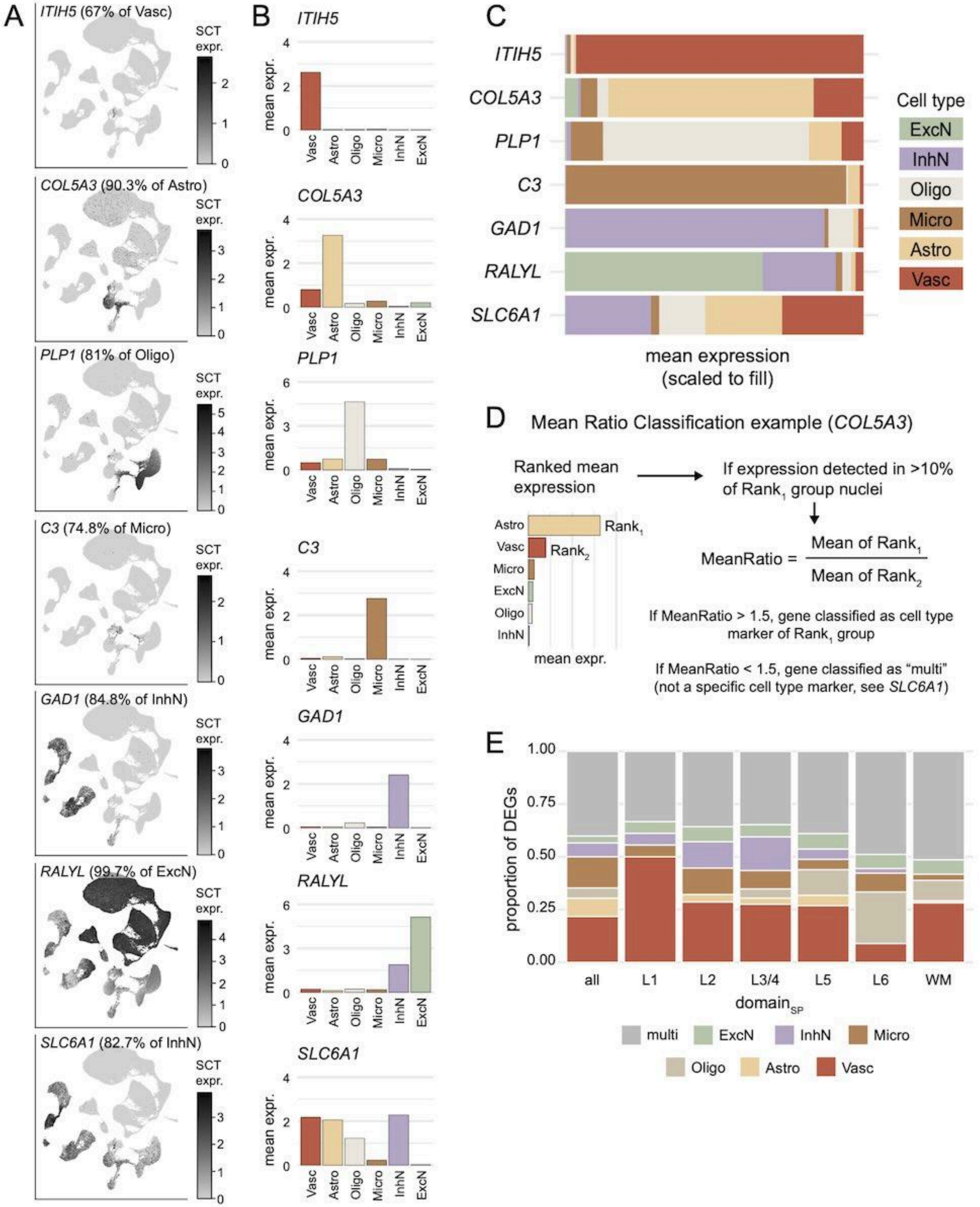

A
ITIH5 (67% of Vasc)
COL5A3 (90.3% of Astro)
PLP1 (81% of Oligo)
C3 (74.8% of Micro)
GAD1 (84.8% of InhN)
RALYL (99.7% of ExcN)
SLC6A1 (82.7% of InhN)
SCT expr.
B
mean expr.
Vasc
Astro
Oligo
Micro
InhN
ExcN
C
ITIH5
COL5A3
PLP1
C3
GAD1
RALYL
SLC6A1
mean expression (scaled to fill)
Cell type
ExcN
InhN
Oligo
Micro
Astro
Vasc
D
Mean Ratio Classification example (COL5A3)
Ranked mean expression
If expression detected in >10% of Rank1 group nuclei
Rank1
Rank2
mean expr.
MeanRatio = Mean of Rank1 / Mean of Rank2
If MeanRatio > 1.5, gene classified as cell type marker of Rank1 group
If MeanRatio < 1.5, gene classified as "multi" (not a specific cell type marker, see SLC6A1)
E
proportion of DEGs
1.00
0.75
0.50
0.25
0.00
all
L1
L2
L3/4
L5
L6
WM
domainSP
multi
ExcN
InhN
Micro
Oligo
Astro
Vasc

**Supplemental Figure 11. Spot-level co-expression analysis to identify coordinated transcriptional modules of DEGs.**

(**A**) Pearson correlation of target gene importance weights (rho, fill) for each pair of initial DEG modules (104 modules, rows and columns) identified with `GRNBOOST2`. Heatmap shows blocks of highly correlated modules, indicating similar groups of co-expressed genes are captured by multiple modules.

(**B**) Top predictor module network constructed from all DEG targets (tiny grey nodes) of the 104 modules and the single predictor module (medium grey nodes) with the greatest importance for each DEG target. Subnetworks with less than 6 connected components have been removed from this network image. From each subnetwork, we selected 1-2 top predictor modules (medium black nodes, labeled text).

(**C**) The correlation of target gene importance (rho, fill) for the 19 top predictor modules (rows and columns), revealing high correlation between the importance weights of *COX4I1*- and *UQCRH*-modules and between *EEF1A1*- and *EIF1*-modules.

(**D**) Overlaps between DEG module target genes were refined by pruning lowly-weighted target genes. For example, prior to refining (top) the *PLP1*-module contained some target genes that were much more highly weighted in other modules (e.g. *RPS12*, red = importance weight in another module, black = importance weight in *PLP1*-module). Conversely, some genes were present in other modules despite being much more highly weighted in the *PLP1*-module (e.g. *MAG*). By pruning DEG targets using an importance ratio >1.5 (removing a target gene from a query module if the target gene was present in a different module with an importance >1.5x the query), we reduced the number of overlapping DEGs between modules (bottom). In this example, *MAG* was pruned from the *GFAP*-module and *RPS12* was pruned from the *PLP1*-module, while *DYNC1I2* was kept in both the *PLP1*-module and the *EEF1A1*-module due to the importance ratio <1.5.

(**E**) Spot-level AUCell values averaged (fill, centered and scaled) for each of the final 15 DEG modules (columns) and each $\text{domain}_{SP}$ (rows).

(**F**) Spot-level AUCell values averaged (fill, centered and scaled) for each of the final 15 DEG modules (columns) and each $\text{domain}_{CT}$ (rows).

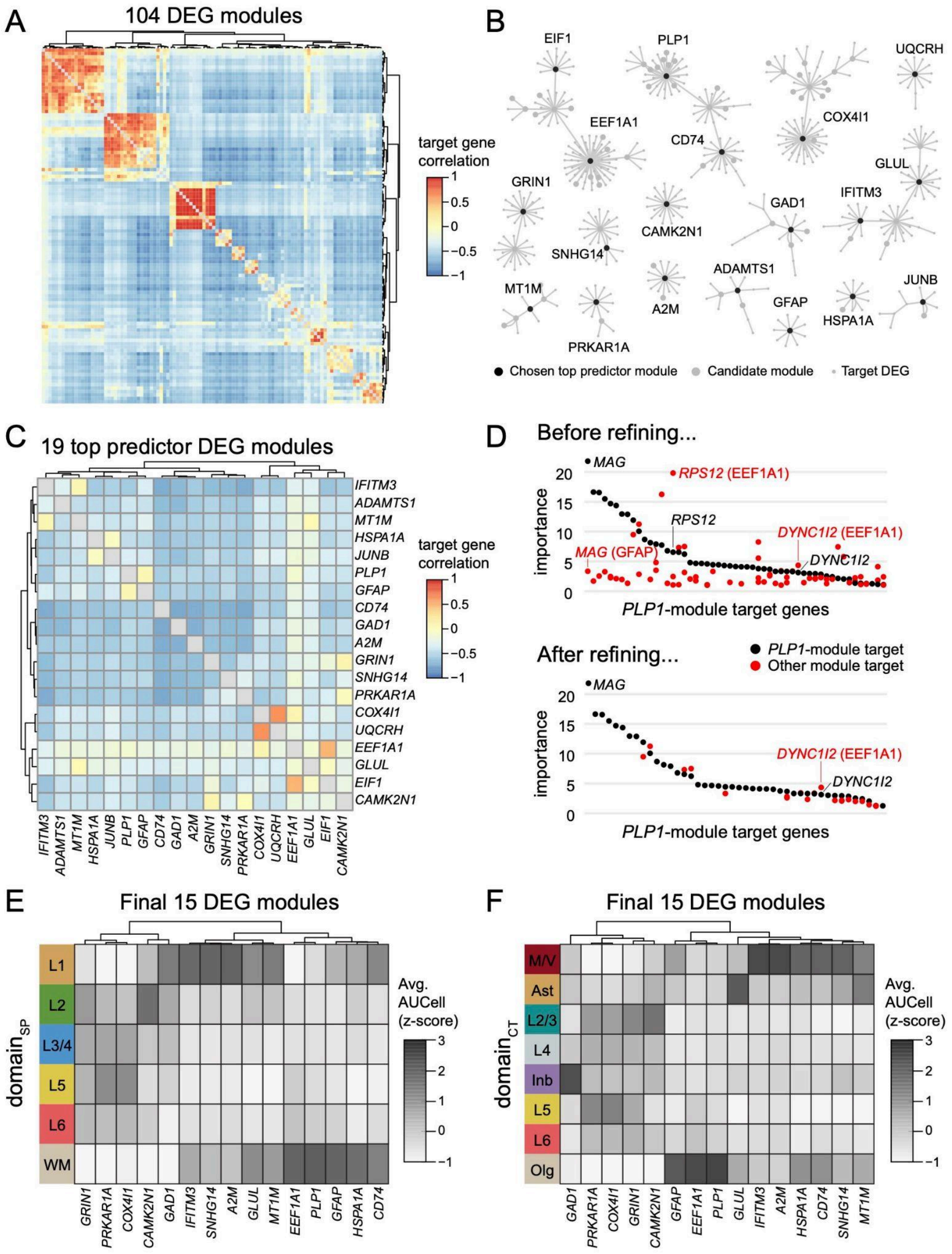

A
104 DEG modules
target gene correlation
1
0.5
0
−0.5
−1
B
EIF1
PLP1
UQCRH
EEF1A1
COX4I1
CD74
GLUL
GRIN1
IFITM3
GAD1
CAMK2N1
SNHG14
ADAMTS1
JUNB
MT1M
A2M
GFAP
HSPA1A
PRKAR1A
Chosen top predictor module
Candidate module
Target DEG
C
19 top predictor DEG modules
IFITM3
ADAMTS1
MT1M
HSPA1A
JUNB
PLP1
GFAP
CD74
GAD1
A2M
GRIN1
SNHG14
PRKAR1A
COX4I1
UQCRH
EEF1A1
GLUL
EIF1
CAMK2N1
target gene correlation
D
Before refining...
MAG
RPS12 (EEF1A1)
RPS12
DYNC1I2 (EEF1A1)
MAG (GFAP)
DYNC1I2
importance
PLP1-module target genes
After refining...
PLP1-module target
Other module target
MAG
DYNC1I2 (EEF1A1)
DYNC1I2
importance
PLP1-module target genes
E
Final 15 DEG modules
domain$_{SP}$
L1
L2
L3/4
L5
L6
WM
Avg. AUCell (z-score)
GRIN1
PRKAR1A
COX4I1
CAMK2N1
GAD1
IFITM3
SNHG14
A2M
GLUL
MT1M
EEF1A1
PLP1
GFAP
HSPA1A
CD74
F
Final 15 DEG modules
domain$_{CT}$
M/V
Ast
L2/3
L4
Inb
L5
L6
Olg
Avg. AUCell (z-score)
GAD1
PRKAR1A
COX4I1
GRIN1
CAMK2N1
GFAP
EEF1A1
PLP1
GLUL
IFITM3
A2M
HSPA1A
CD74
SNHG14
MT1M

**Supplemental Figure 12. Example spot plots of DEG modules corresponding to traditional dlPFC cytoarchitectural layers.** Normalized AUCell (spot fill) spot plots for the 15 DEG modules in example sample Br6432. Spot border color indicates $domain_{SP}$.

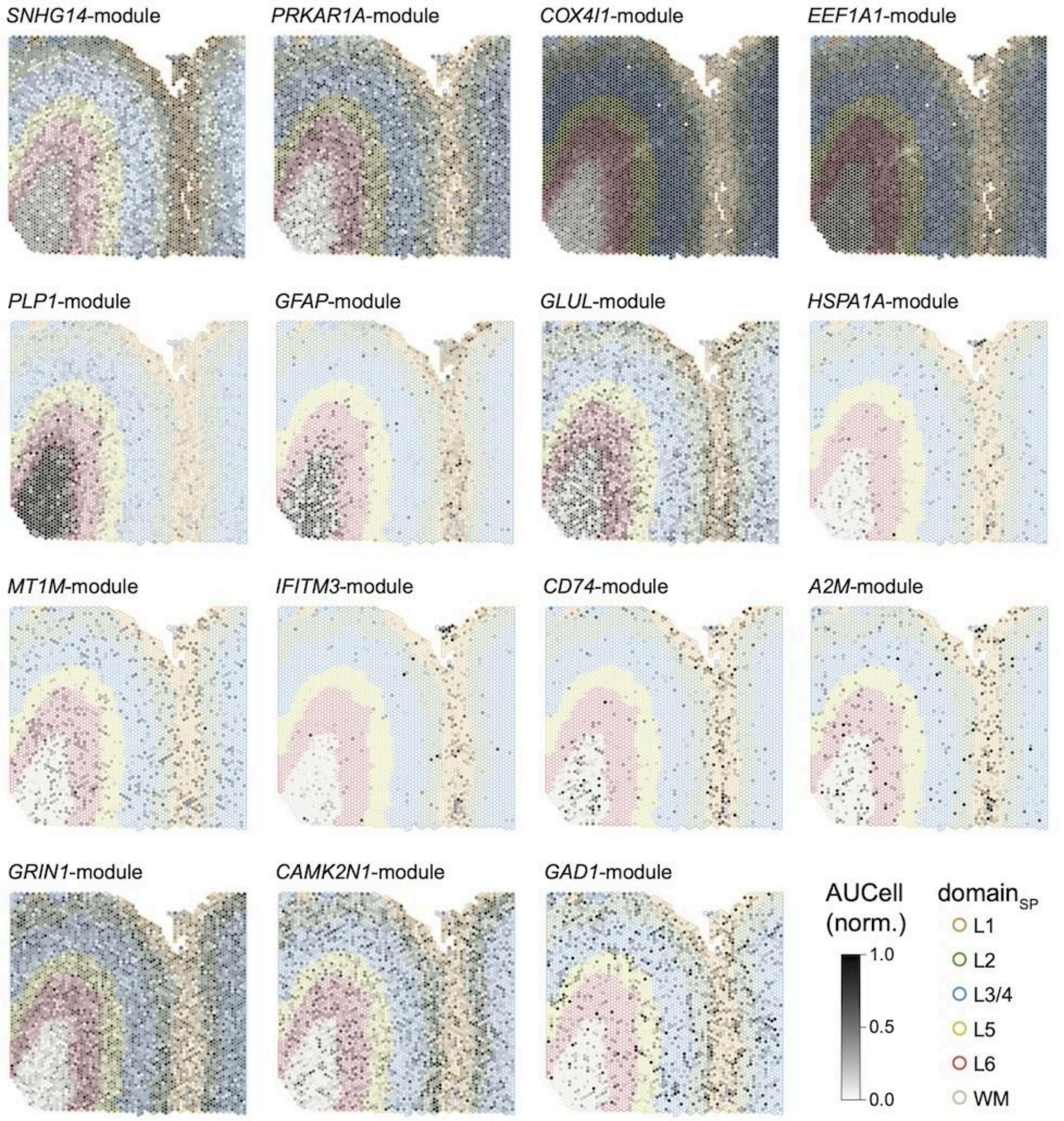

**Supplemental Figure 13. DEGs dropped from modules after applying filters.**

(**A**) (**i**) Dotplot of DEGs that were not associated with a co-expression module after filtering by importance weight, minimum module size, and minimum number of DEGs. Differential expression results are presented across domain annotations (x-axis, ‘all’ represents whole-tissue model results) where large size indicates DEG status, shape indicates domain annotation strategy, and color reflects $\log_2$ fold change (logFC). Fold change is calculated for each diagnosis-by-sex group (facet) with the reference group listed first and the comparison group listed second (i.e. NTC.MDD with logFC < 0 reflects expression where MDD is less than NTC). Some genes (e.g. *LINC00943*) were cut from DE modeling during the `edgeR` minimum expression pre-filtering and therefore only have DE results present for one domain annotation. (**ii**) Mean expression of *y*-axis DEGs for each broad cell type from snRNA-seq, scaled to fill *x*-axis. *SLC25A37* does not have corresponding mean expression data because this gene was not present in the snRNA-seq dataset.

(**B**) Overrepresentation analysis of GO-BP terms showed DEGs missing from modules are enriched for genes related to biological processes represented in some DEG modules.

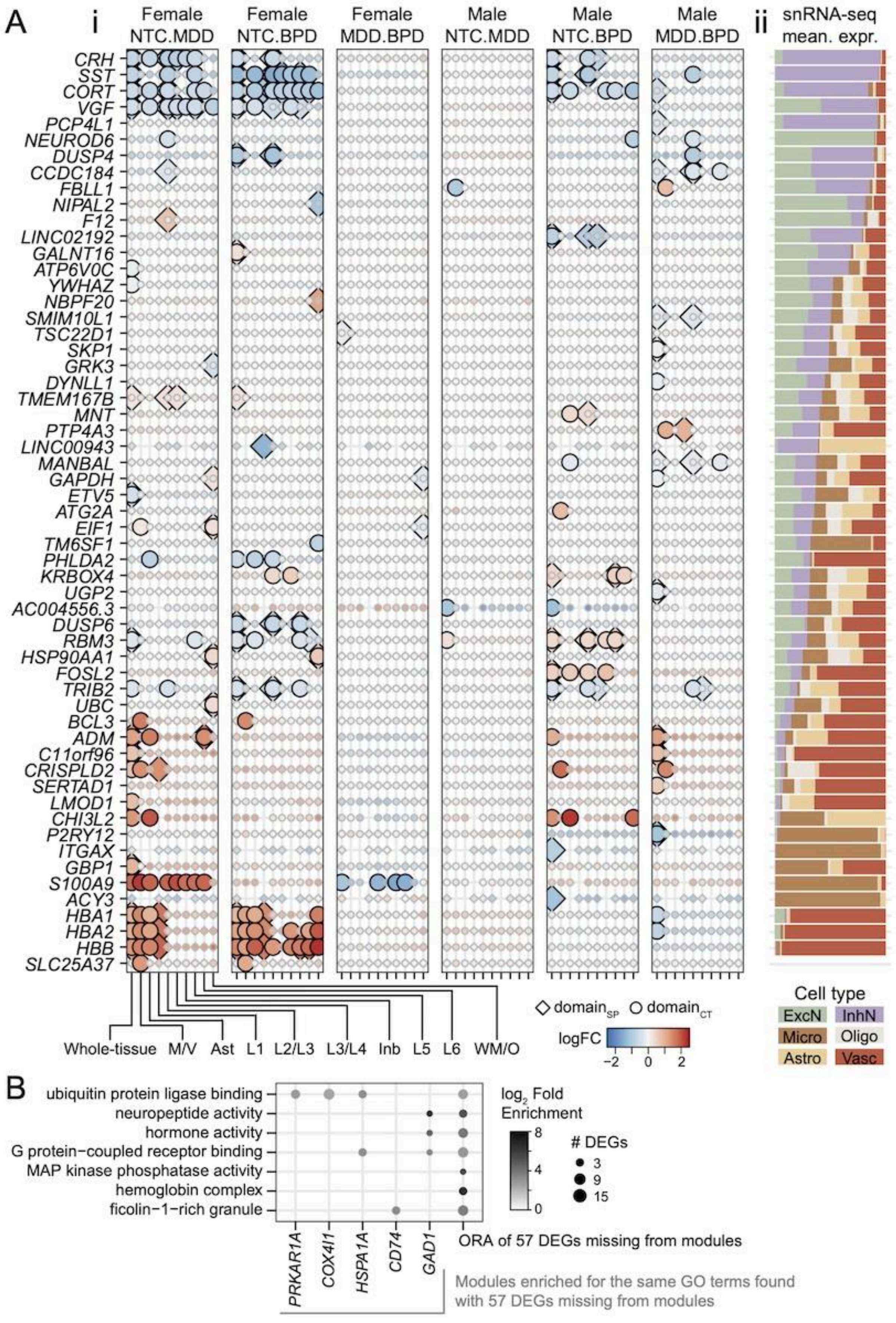

A
i
Female NTC.MDD
Female NTC.BPD
Female MDD.BPD
Male NTC.MDD
Male NTC.BPD
Male MDD.BPD
ii
snRNA-seq mean. expr.
CRH
SST
CORT
VGF
PCP4L1
NEUROD6
DUSP4
CCDC184
FBLL1
NIPAL2
F12
LINC02192
GALNT16
ATP6V0C
YWHAZ
NBPF20
SMIM10L1
TSC22D1
SKP1
GRK3
DYNLL1
TMEM167B
MNT
PTP4A3
LINC00943
MANBAL
GAPDH
ETV5
ATG2A
EIF1
TM6SF1
PHLDA2
KRBOX4
UGP2
AC004556.3
DUSP6
RBM3
HSP90AA1
FOSL2
TRIB2
UBC
BCL3
ADM
C11orf96
CRISPLD2
SERTAD1
LMOD1
CHI3L2
P2RY12
ITGAX
GBP1
S100A9
ACY3
HBA1
HBA2
HBB
SLC25A37
Whole-tissue
M/V
Ast
L1
L2/L3
L3/L4
Inb
L5
L6
WM/O
domain_SP
domain_CT
logFC
−2
0
2
Cell type
ExcN
InhN
Micro
Oligo
Astro
Vasc
B
ubiquitin protein ligase binding
neuropeptide activity
hormone activity
G protein−coupled receptor binding
MAP kinase phosphatase activity
hemoglobin complex
ficolin−1−rich granule
log2 Fold Enrichment
8
4
0
### DEGs
3
9
15
PRKAR1A
COX4I1
HSPA1A
CD74
GAD1
ORA of 57 DEGs missing from modules
Modules enriched for the same GO terms found with 57 DEGs missing from modules

**Supplemental Figure 14. Module DEG distribution across domains and diagnosis-by-sex groups.**

(**A**) Heatmap of DE results split by domain annotation strategies (facet columns), indicating the proportion of component DEGs from each DEG co-expression module (facet row, labeled with total number of DEGs in module), and *y*-axis rows representing the direction of change (increased if logFC>0 or decreased if logFC<0). *X*-axis columns distinguish the whole-tissue model ("all") from the different domain-restricted model results.

(**B**) Heatmap of DE results split by diagnosis-by-sex group (facet columns), indicating the proportion of component DEGs from each DEG co-expression module (facet row, labeled with total number of DEGs in module), and *y*-axis rows representing the direction of change (increased if logFC>0 or decreased if logFC<0). *X*-axis columns indicate aggregated results for the whole-tissue model and the domain-restricted model for each of the two domain annotation strategies.

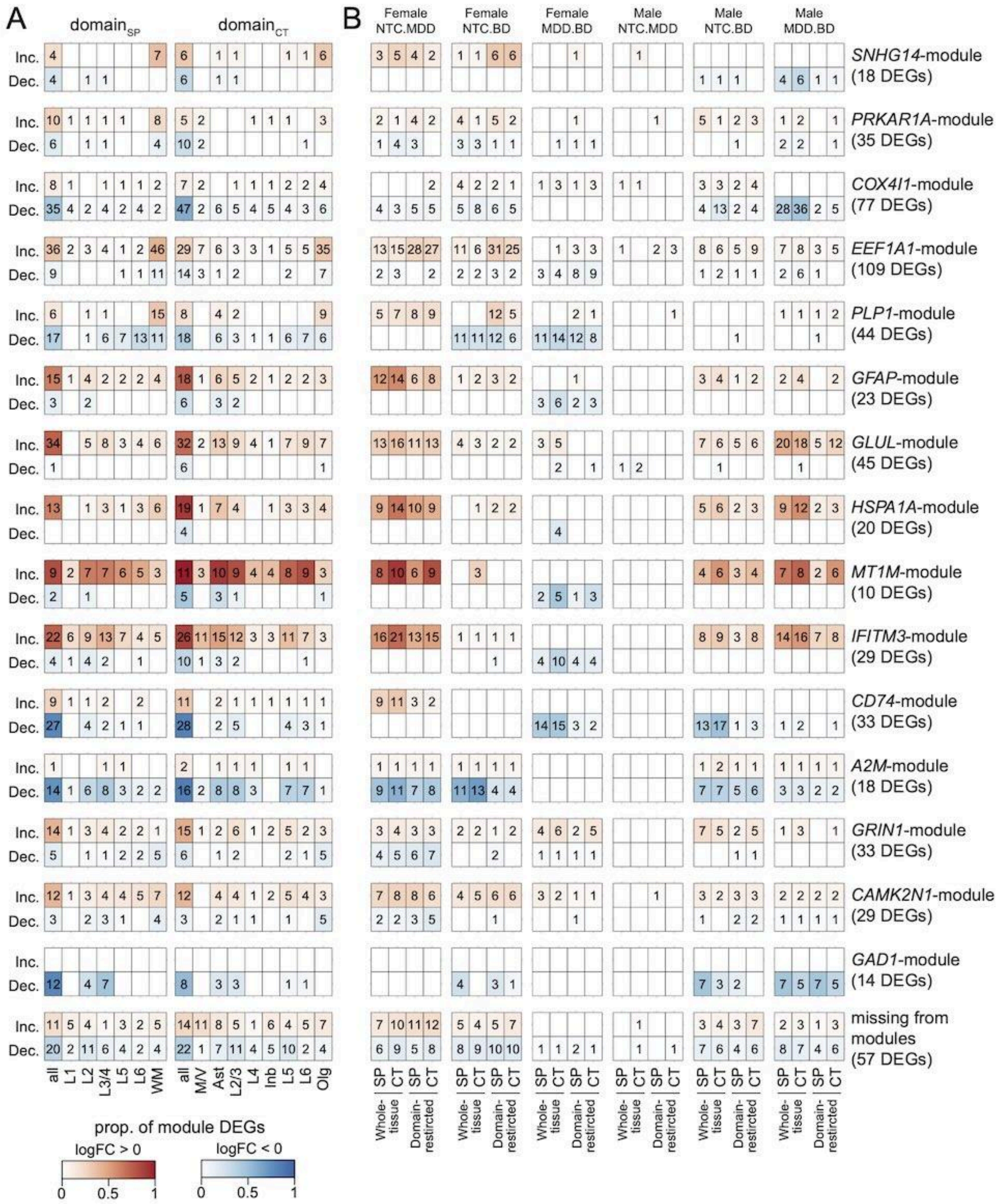
A
domain$_{SP}$
domain$_{CT}$
B
Female NTC.MDD
Female NTC.BD
Female MDD.BD
Male NTC.MDD
Male NTC.BD
Male MDD.BD
Inc.
Dec.
SNHG14-module (18 DEGs)
PRKAR1A-module (35 DEGs)
COX4I1-module (77 DEGs)
EEF1A1-module (109 DEGs)
PLP1-module (44 DEGs)
GFAP-module (23 DEGs)
GLUL-module (45 DEGs)
HSPA1A-module (20 DEGs)
MT1M-module (10 DEGs)
IFITM3-module (29 DEGs)
CD74-module (33 DEGs)
A2M-module (18 DEGs)
GRIN1-module (33 DEGs)
CAMK2N1-module (29 DEGs)
GAD1-module (14 DEGs)
missing from modules (57 DEGs)
all L1 L2 L3/4 L5 L6 WM
all M/V Ast L2/3 L4 Inb L5 L6 Olg
SP CT SP CT
Whole-tissue
Domain-restricted
prop. of module DEGs
logFC > 0
logFC < 0
0 0.5 1
0 0.5 1

**Supplemental Figure 15. Regulon analysis reveals strong activity-dependent transcription factor response in a small subset of donors.**

(**A**) Regulon analysis performed on all 119 donors revealed 3 AP1-family regulons (red dashed box) with a small number of donors exhibiting much higher average AUCell values (*y*-axis). Each donor is represented by a single black line, and the average AUCell across all donors for each transcription factor regulon (*x*-axis) is represented by a grey dot.

(**B**) All 119 donors were ranked by the average AUCell of the 3 AP1-family regulons in question (*x*-axis), with the vertical dashed line separating the top 10 donors. The top plot shows the average AUCell (*y*-axis) for each of the three outlier regulons (color, FOS, FOSB, and JUNB). The bottom plot shows the proportion of spots expressing *NPAS4* (*y*-axis), with donors labeled by sex (shape) and diagnosis group (color).

(**C**) Spot plots of *NPAS4* expression (spot fill, logcounts) for the top 12 donors (ranked by average AUCell of 3 AP1-family regulons in question) demonstrate the clear outlier nature of the top 10 donors (dashed outline). Spot border indicates $\text{domain}_{\text{SP}}$.

(**D**) Regulon analysis performed on 109 donors (removing 10 *NPAS4* outlier donors) no longer shows AP1-family outliers.

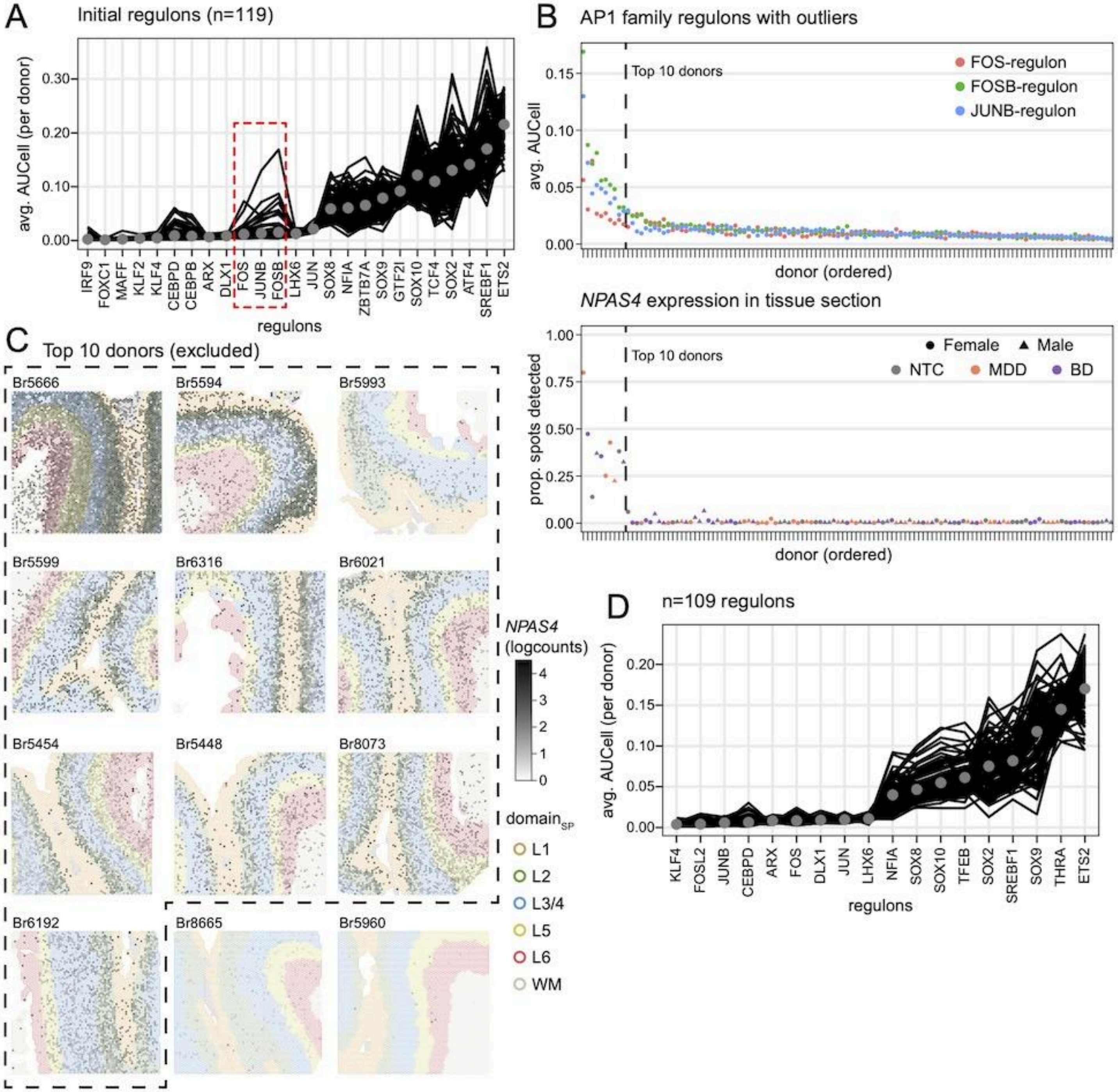

A
Initial regulons (n=119)
avg. AUCell (per donor)
regulons
IRF9
FOXC1
MAFF
KLF2
KLF4
CEBPD
CEBPB
ARX
DLX1
FOS
JUNB
FOSB
LHX6
JUN
SOX8
NFIA
ZBTB7A
SOX9
GTF2I
SOX10
TCF4
SOX2
ATF4
SREBF1
ETS2
B
AP1 family regulons with outliers
Top 10 donors
FOS-regulon
FOSB-regulon
JUNB-regulon
avg. AUCell
donor (ordered)
NPAS4 expression in tissue section
Female
Male
NTC
MDD
BD
prop. spots detected
C
Top 10 donors (excluded)
Br5666
Br5594
Br5993
Br5599
Br6316
Br6021
Br5454
Br5448
Br8073
Br6192
Br8665
Br5960
NPAS4
(logcounts)
domain
L1
L2
L3/4
L5
L6
WM
D
n=109 regulons
KLF4
FOSL2
JUNB
CEBPD
ARX
FOS
DLX1
JUN
LHX6
NFIA
SOX8
SOX10
TFEB
SOX2
SREBF1
SOX9
THRA
ETS2

**Supplemental Figure 16. SCENIC regulon analysis (n=109 donors) to identify transcription factor regulons associated with spot-level expression patterns.**

(**A**) Heatmap of spot-level AUCell correlation (rho, fill color) between pairs of regulons (rows and columns) detected for n=109 donors shows groups of correlated regulons.

(**B**) Heatmap of spot-level AUCell correlation (rho, fill color) between regulons (rows) and DEG modules (columns). Text labels indicate the number of overlapping genes between regulons and modules.

(**C**) Heatmap of spot-level AUCell values averaged for each regulon (columns) and each $domain_{SP}$ (rows). Heatmap color indicates the average AUCell value, centered and scaled.

(**D**) Heatmap of spot-level AUCell values averaged for each regulon (columns) and each $domain_{CT}$ (rows). Heatmap color indicates the average AUCell value, centered and scaled.

(**E**) Normalized AUCell (spot fill) spot plots for regulons with example sample Br6432. Spot border color indicates $domain_{SP}$.

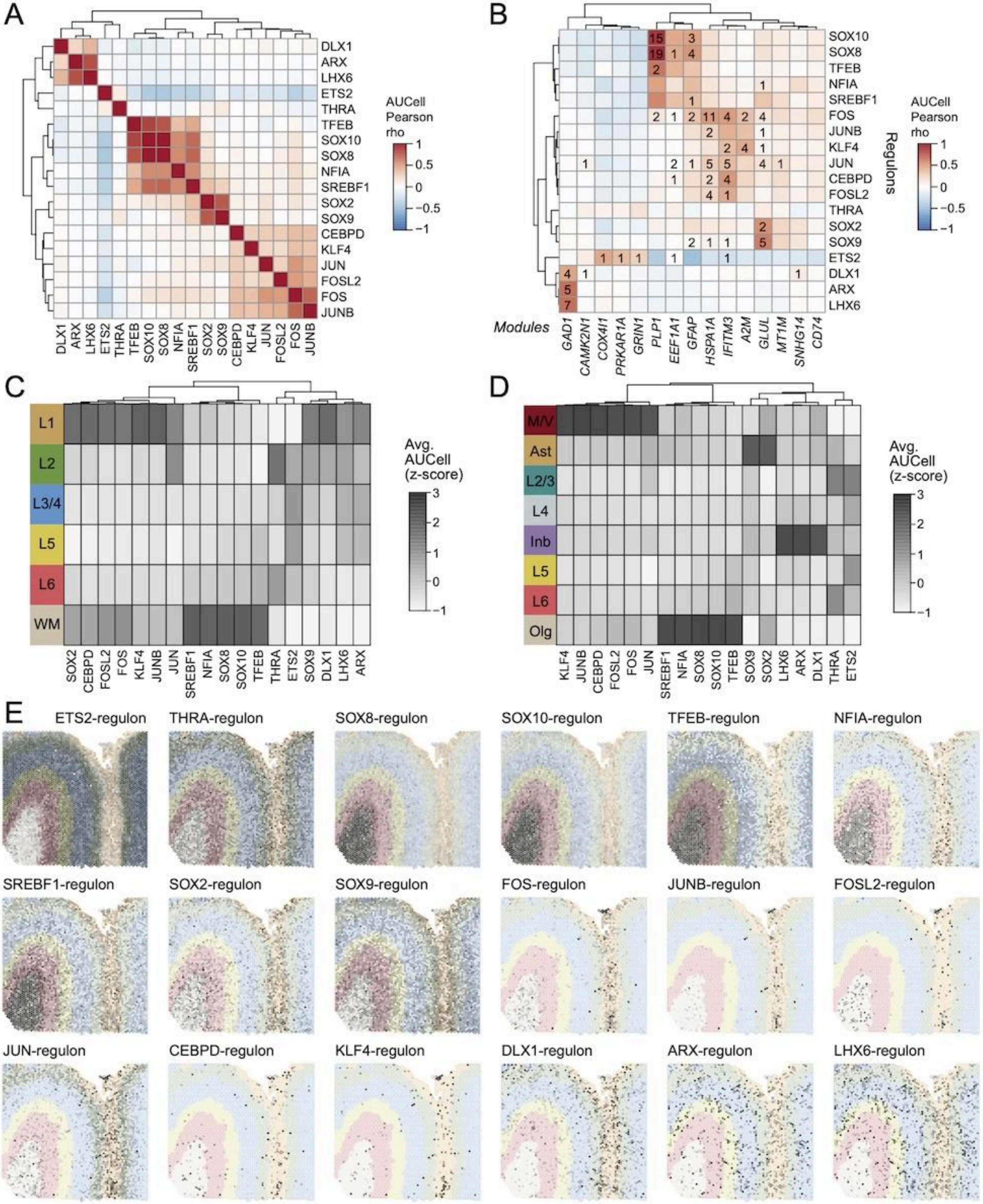
A
DLX1
ARX
LHX6
ETS2
THRA
TFEB
SOX10
SOX8
NFIA
SREBF1
SOX2
SOX9
CEBPD
KLF4
JUN
FOSL2
FOS
JUNB
AUCell Pearson rho
1
0.5
0
−0.5
−1
B
SOX10
SOX8
TFEB
NFIA
SREBF1
FOS
JUNB
KLF4
JUN
CEBPD
FOSL2
THRA
SOX2
SOX9
ETS2
DLX1
ARX
LHX6
Regulons
Modules
GAD1
CAMK2N1
COX4I1
PRKAR1A
GRIN1
PLP1
EEF1A1
GFAP
HSPA1A
IFITM3
A2M
GLUL
MT1M
SNHG14
CD74
AUCell Pearson rho
C
L1
L2
L3/4
L5
L6
WM
SOX2
CEBPD
FOSL2
FOS
KLF4
JUNB
JUN
SREBF1
NFIA
SOX8
SOX10
TFEB
THRA
ETS2
SOX9
DLX1
LHX6
ARX
Avg. AUCell (z-score)
3
2
1
0
−1
D
M/V
Ast
L2/3
L4
Inb
L5
L6
Olg
KLF4
JUNB
CEBPD
FOSL2
FOS
JUN
SREBF1
NFIA
SOX8
SOX10
TFEB
SOX9
SOX2
LHX6
ARX
DLX1
THRA
ETS2
Avg. AUCell (z-score)
E
ETS2-regulon
THRA-regulon
SOX8-regulon
SOX10-regulon
TFEB-regulon
NFIA-regulon
SREBF1-regulon
SOX2-regulon
SOX9-regulon
FOS-regulon
JUNB-regulon
FOSL2-regulon
JUN-regulon
CEBPD-regulon
KLF4-regulon
DLX1-regulon
ARX-regulon
LHX6-regulon

**Supplemental Figure 17**. **Female-specific mood disorder-associated changes in gene expression in white matter and oligodendrocyte-dominant spots.**

(**A**) $WM_{SP}$ domain volcano plots for F_NTC.MDD (left) and F_NTC.BD (right). Highlighted genes indicate DEGs identified with domain-restricted $Olg_{CT}$ and $WM_{SP}$ DE models that were not found in whole-tissue analysis.

(**B**) Violin and box plots of *ANP32B* expression for $domain_{SP}$ pseudobulk data with logcounts on *y*-axis and domains on *x*-axis ('all' on *x*-axis represents whole-tissue model results from all pseudobulk samples). Statistically significant *t*-test (adj. $p<.05$) versus sex-matched control group indicated by asterisk. Colors correspond to different groups and group comparisons that are statistically significant are in a dark color. Crossbar indicates mean ± standard deviation.

(**C**) (**i**) Dotplot of select *EEF1A1*-module genes across domain annotations (x-axis, 'all' represents whole-tissue model results) where large size indicates DEG status, shape indicates domain annotation strategy, and color reflects $log_2$ fold change (logFC). Fold change is calculated for each diagnosis-by-sex group (facet) with the reference group listed first and the comparison group listed second (i.e. NTC.MDD with logFC < 0 reflects expression where MDD is less than NTC). (**ii**) Mean expression of y-axis *EEF1A1*-module genes for each broad cell type from snRNA-seq, scaled to fill x-axis.

(**D-E**) Violin plot of *CLDN11* and *SGK1* expression, respectively) for $domains_{SP}$. Same as (**B**), with additional bar symbols indicating statistical significance (adj. $p<.05$) between MDD and BD.

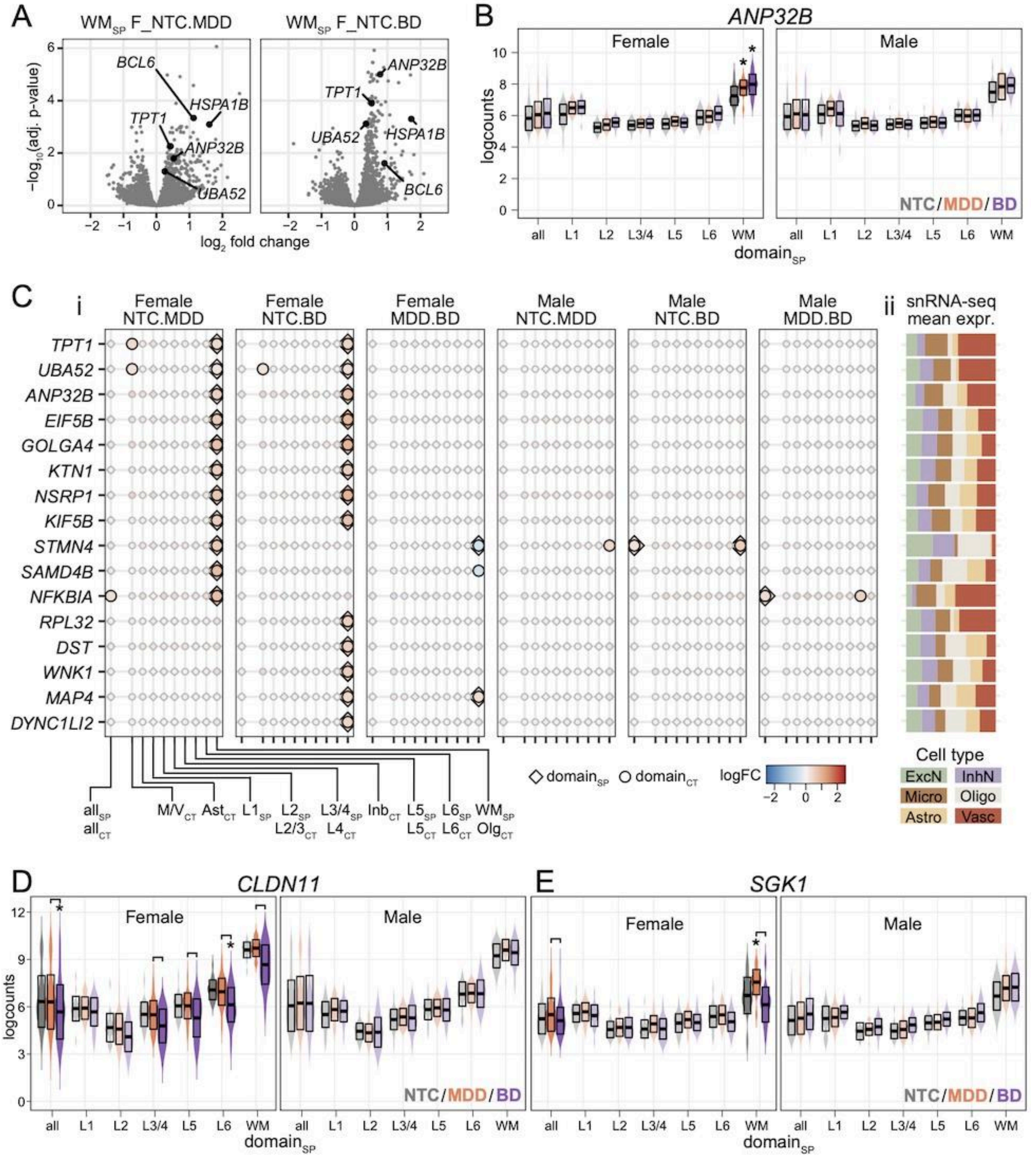

A
WM_SP F_NTC.MDD
WM_SP F_NTC.BD
BCL6
HSPA1B
TPT1
ANP32B
UBA52
−log10(adj. p-value)
log2 fold change
B
ANP32B
Female
Male
logcounts
NTC/MDD/BD
all L1 L2 L3/4 L5 L6 WM
domain_SP
C
i
Female NTC.MDD
Female NTC.BD
Female MDD.BD
Male NTC.MDD
Male NTC.BD
Male MDD.BD
ii
snRNA-seq mean expr.
TPT1
UBA52
ANP32B
EIF5B
GOLGA4
KTN1
NSRP1
KIF5B
STMN4
SAMD4B
NFKBIA
RPL32
DST
WNK1
MAP4
DYNC1LI2
all_SP all_CT
M/V_CT
Ast_CT
L1_SP
L2_SP L2/3_CT
L3/4_SP L4_CT
Inb_CT
L5_SP L5_CT
L6_SP L6_CT
WM_SP Olg_CT
domain_SP
domain_CT
logFC
Cell type
ExcN
InhN
Micro
Oligo
Astro
Vasc
D
CLDN11
Female
Male
logcounts
NTC/MDD/BD
domain_SP
E
SGK1
Female
Male
NTC/MDD/BD
domain_SP

**Supplemental Figure 18. Supporting evidence of astrocyte reactivity and neuroimmune response in mood disorders.**

(**A**) (**i**) Dotplot of *GFAP*-module genes that are overlapping with oligo phenotype (*PLP1*-module, SOX8-regulon, or SOX10-regulon) or astrocyte phenotype (*GLUL*-module, SOX2-regulon or SOX9-regulon). DE results are shown across domain annotations (x-axis, 'all' represents whole-tissue model results) where large size indicates DEG status, shape indicates domain annotation strategy, and color reflects $\log_2$ fold change (logFC). Fold change is calculated for each diagnosis-by-sex group, with comparisons to NTC shown in facets. (**ii**) Mean expression of corresponding genes in (i) for each broad cell type from snRNA-seq, scaled to fill x-axis.

(**B**)  Dotplot for all *CD74*-module DEGs, see legend of (**A**) for details.

(**C**) Proportion of spots with detected expression (median of all n=119 donors, *y*-axis) for all *CD74*-module DEGs. Red line corresponds to half of donors with detected expression in 1% of spots, highlighting the sparse nature of many *CD74*-module DEGs.

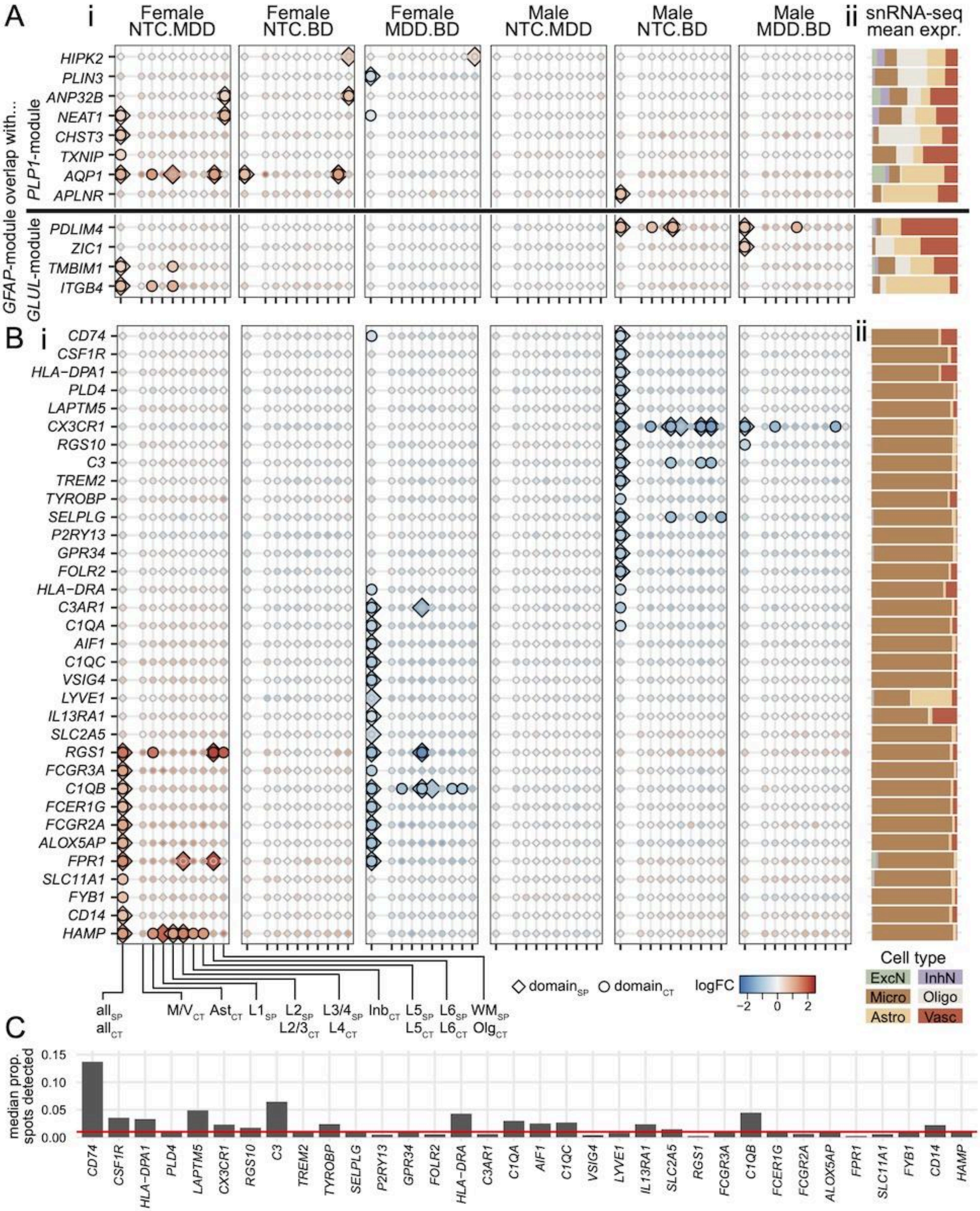
A
i
Female NTC.MDD
Female NTC.BD
Female MDD.BD
Male NTC.MDD
Male NTC.BD
Male MDD.BD
ii
snRNA-seq mean expr.
overlap with...
PLP1-module
GFAP-module
GLUL-module
HIPK2
PLIN3
ANP32B
NEAT1
CHST3
TXNIP
AQP1
APLNR
PDLIM4
ZIC1
TMBIM1
ITGB4
B
CD74
CSF1R
HLA-DPA1
PLD4
LAPTM5
CX3CR1
RGS10
C3
TREM2
TYROBP
SELPLG
P2RY13
GPR34
FOLR2
HLA-DRA
C3AR1
C1QA
AIF1
C1QC
VSIG4
LYVE1
IL13RA1
SLC2A5
RGS1
FCGR3A
C1QB
FCER1G
FCGR2A
ALOX5AP
FPR1
SLC11A1
FYB1
CD14
HAMP
all SP
all CT
M/V CT
Ast CT
L1 SP
L2 SP
L2/3 CT
L3/4 SP
L4 CT
Inb CT
L5 SP
L5 CT
L6 SP
L6 CT
WM SP
Olg CT
domain SP
domain CT
logFC
−2
0
2
Cell type
ExcN
InhN
Micro
Oligo
Astro
Vasc
C
median prop. spots detected
0.15
0.10
0.05
0.00

**Supplemental Figure 19**. **Inhibitory neuron involvement in BD-associated gene expression changes.**

(**A**) Bar plot showing the number of nuclei in native Azimuth labels for *SZBDMulti-seq* snRNA-seq dataset (Ruzicka et al. 2024), colored by collapsed inhibitory subtype and lineage groups.

(**B**) Violin plots of *SZBDMulti-seq* pseudobulk expression (y-axis, logcounts) for collapsed inhibitory subtype (x-axis). (Top) Genes that are shown support the CGE inhibitory subtypes: broad CGE marker *ADARB2* and CGE subtype markers *CNR1* and *LAMP5*. (Bottom) Genes that are shown support the MGE inhibitory subtypes: broad MGE marker *LHX6* and MGE subtype markers *SST* and *CORT*.

(**C**) GAD1-module dotplot (i) with mean expression of broad cell types (ii) and inhibitory subtype and lineage groups (iii). Lines separate MGE markers (top) from general inhibitory markers (middle) and CGE markers (bottom).

(**D**) Merged smFISH images (*GAD1* (green), *AIF1* (yellow), *TRBC2* (pink) for 3 NTC donors (Br5558-M, Br1113-F, Br6432-M) not presented in **Figure 5G** demonstrating *TRBC2* co-expression with *GAD1* (white arrow) and not with immune cell type marker *AIF1* (yellow arrow).

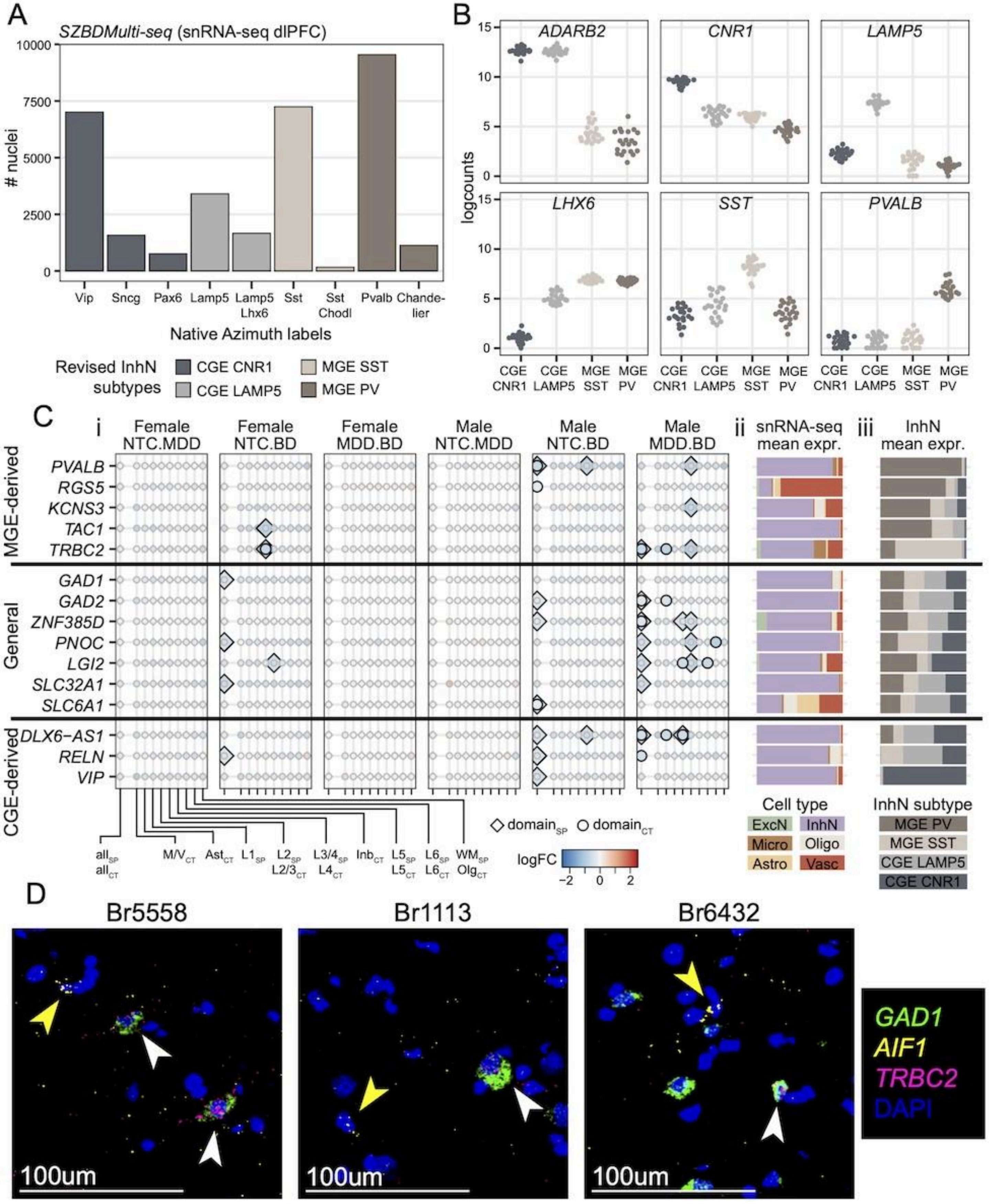

A
SZBDMulti-seq (snRNA-seq dlPFC)
### nuclei
Vip Sncg Pax6 Lamp5 Lamp5 Lhx6 Sst Sst Chodl Pvalb Chandelier
Native Azimuth labels
Revised InhN subtypes
CGE CNR1
CGE LAMP5
MGE SST
MGE PV
B
ADARB2
CNR1
LAMP5
LHX6
SST
PVALB
logcounts
C
i
Female NTC.MDD
Female NTC.BD
Female MDD.BD
Male NTC.MDD
Male NTC.BD
Male MDD.BD
ii
snRNA-seq mean expr.
iii
InhN mean expr.
MGE-derived
PVALB
RGS5
KCNS3
TAC1
TRBC2
General
GAD1
GAD2
ZNF385D
PNOC
LGI2
SLC32A1
SLC6A1
CGE-derived
DLX6-AS1
RELN
VIP
domain_SP
domain_CT
logFC
Cell type
ExcN
InhN
Micro
Oligo
Astro
Vasc
InhN subtype
MGE PV
MGE SST
CGE LAMP5
CGE CNR1
D
Br5558
Br1113
Br6432
100um
GAD1
AIF1
TRBC2
DAPI

**Supplemental Figure 20. Trait-associated genetic risk at sites of eQTL association in mood disorder cohort.**

(**A**) Bar plots of the number of participants in the latest European-ancestry GWAS studies from the Psychiatric Genomics Consortium including MDD (Adams et al. 2025), BD (O'Connell et al. 2025), and SCZ (Trubetskoy et al. 2022).

(**B**) Number of variants identified for each GWAS set using standard $p$<5e-08 threshold.

(**C**) Number of variants identified for each GWAS set when mood disorder sets ($GWAS_{MDD}$ and $GWAS_{BD}$) utilized exploratory threshold ($p$<1e-5).

(**D**) Table summary of overlaps between DEGs, GWAS exact variant support, and the results from eQTL analysis and colocalization analysis. Rows are split by whether the associated gene was identified as a DEG in this study. Numeric values indicate the number of unique variants with GWAS exact variant support at the indicated threshold; note that multiple variants can be within the same loci. For results with exact variant support that are associated with expression of a DEG, the associated gene name is listed in parentheses.

(**E**) A breakdown of the GWAS exact variant support for variants associated with DEGs with eQTL and/or colocalization analysis. The diagnosis-by-sex group contrasts for which the associated gene was identified as a significant DEG is given. eQTL and colocalization analyses columns list the different $domain_{CT}$ contexts for which the association was identified as significant.

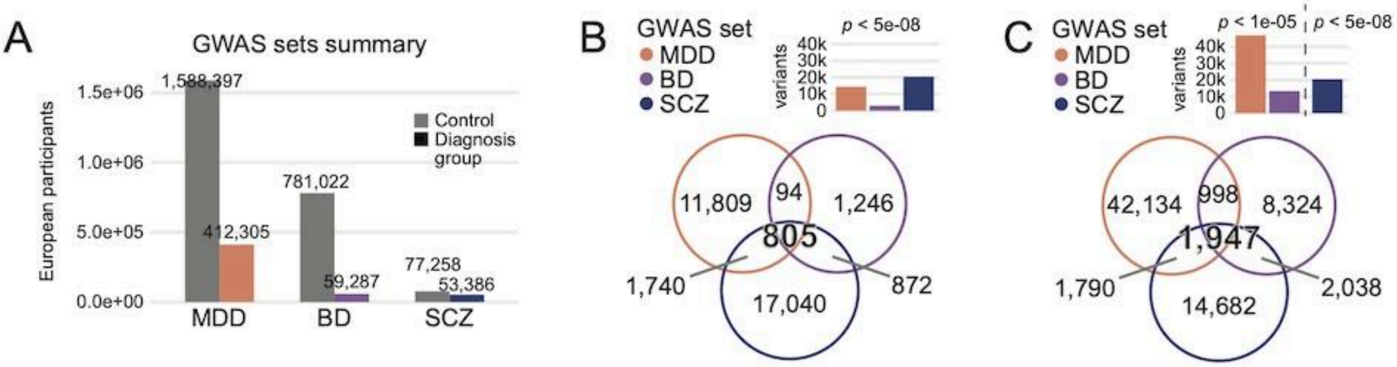


D

| | | Variant associated with genetic risk? (standard, $p < 5e-08$) | | | Variant associated with genetic risk? (exploratory, $p < 1e-05$) | |
|---|---|---|---|---|---|---|
| | | $GWAS_{MDD}$ | $GWAS_{BD}$ | $GWAS_{SCZ}$ | $GWAS_{MDD}$ | $GWAS_{BD}$ |
| Variant associated with DEG via significant **eQTL**? | non-DEG association | 72 | 68 | 136 | 120 | 75 |
| | DEG association | 0 | 0 | **2** (*MAPK3* x2) | **2** (*MAPK3*, *SPON2*) | **2** (*MAPK3* x2) |
| Variant associated with DEG via **colocalization**? | non-DEG association | 7 | 6 | 8 | 16 | 12 |
| | DEG association | 0 | 0 | **1** (*PTP5A3*) | **2** (*MAPK3*, *FBLN7*) | **1** (*MAPK3*) |

E

| Variant ID | RSID | Associated DEG | DE group | Variant associated with DEG via significant **eQTL**? | Variant associated with DEG via **colocalization**? | Variant associated with genetic risk? $GWAS_{MDD}$ | $GWAS_{BD}$ | $GWAS_{SCZ}$ |
|---|---|---|---|---|---|---|---|---|
| chr16:30130664:T:C | rs55732507 | *MAPK3* | F_NTC.BD, F_MDD.BD, M_NTC.BD | Ast, L2/3, L4, L5 | Ast, L2/3, Inb, L5 | | $p < 1e-05$ | $p < 5e-08$ |
| chr16:30123335:T:C | rs28529403 | | | Ast, L2/3, L4, L5 | | | $p < 1e-05$ | $p < 5e-08$ |
| chr16:30311847:G:C | rs148788997 | | | Inb | Inb | $p < 1e-05$ | | |
| chr4:1398767:G:T | rs13119951 | *SPON2* | F_NTC.MDD | L6 | | $p < 1e-05$ | | |
| chr2:112462329:C:T | rs13390931 | *FBLN7* | F_NTC.MDD | | Ast, L2/3 | $p < 1e-05$ | | |
| chr8:142231572:A:C | rs4129585 | *PTP4A3* | M_MDD.BD | | L4 | | | $p < 5e-08$ |

### Supplementary Table

**Supplementary Table 1. Demographic variables for study donors.** Demographic information for each donor included in the study, including sex, age, diagnosis, and RNA integrity number (RIN).

**Supplementary Table 2. Results from domain$_{SP}$ dx-sex differential expression (DE) models.** For each DE model (whole-tissue and domain-restricted) there are two sheets containing the results of the moderated omnibus *F*-test and the results of the moderated *t*-test. Moderated *F*-test results sheets are filtered to only genes with adj. $p<.05$ and include columns with the coefficients ($\log_2$ fold change estimate) for each of the contrasts tested in the model, columns for *F*-test statistics (average $\log_2$ expression across all pseudobulk samples, moderated *F*-statistic, raw *p*-value, adjusted *p*-value, the total degrees of freedom), columns for the gene details (ENSEMBL ID, gene name, *F*-test design, median proportion of spots with gene detected for each donor), and columns containing summaries of the *t*-test results (number of contrasts with significant *t*-tests, binned *t*-test significance across contrasts). Moderated *t*-test results sheets include the $\log_2$ fold change estimate, average $\log_2$ expression across all pseudobulk samples, moderated *t*-statistic, raw *p*-value, adjusted *p*-value, log-odds that the gene is differentially expressed (B), gene details, and design details (diagnosis-by-sex coefficient tested, domain tested, sex, diagnosis group comparison, linear model, and domain annotation).

**Supplementary Table 3. Results from domain$_{CT}$ dx-sex differential expression models.** For each DE model (whole-tissue and domain-restricted) there are two sheets containing the results of the moderated omnibus *F*-test and the results of the moderated *t*-test. Moderated *F*-test results sheets are filtered to only genes with adj. $p<.05$ and include columns with the coefficients ($\log_2$ fold change estimate) for each of the contrasts tested in the model, columns for *F*-test statistics (average $\log_2$ expression across all pseudobulk samples, moderated *F*-statistic, raw *p*-value, adjusted *p*-value, the total degrees of freedom), columns for the gene details (ENSEMBL ID, gene name, *F*-test design, median proportion of spots with gene detected for each donor), and columns containing summaries of the *t*-test results (number of contrasts with significant *t*-tests, binned *t*-test significance across contrasts). Moderated *t*-test results sheets include the $\log_2$ fold change estimate, average $\log_2$ expression across all pseudobulk samples, moderated *t*-statistic, raw *p*-value, adjusted *p*-value, log-odds that the gene is differentially expressed (B), gene details, and design details (diagnosis-by-sex coefficient tested, domain tested, sex, diagnosis group comparison, linear model, and domain annotation).

**Supplementary Table 4. Refined GRNBOOST2 output for 15 DEG modules.** Adjacencies for the 15 DEG modules (‘source’) and all component DEGs (‘target’) above the minimum importance filter (importance >1), listing the importance weight (‘importance’) output from GRNBOOST2 and the Pearson correlation rho (‘rho’). This is the final module table after all filters and refinement.

**Supplementary Table 5. SCENIC regulons.** Results of SCENIC analysis identifying regulon source transcription factor (‘TF’), the number of co-expressed genes with enrichment for TF motif around gene location (‘regulon_size’), and the identity of these predicted regulatory targets (‘regulon_targets’), with different genes delineated by forward slash. This is the final regulon table after all filters.

**Supplementary Table 6. Context-stratified domain$_{CT}$ cis-eQTL results**. This Excel workbook contains worksheets *cis_independent*, *nominal_BH05*, *eQTL_summary*, and *intersecting_genes* with all-donor domain$_{CT}$ eQTL results generated with tensorQTL and augmented with DEG and psychiatric GWAS annotations. Rows are pooled per domain$_{CT}$ contexts, reported in the *context* column.

1) *cis_independent* sheet

This sheet contains one row per unique significant gene-variant pair from the integrated lead cis and conditionally independent eQTL results computed with *tensorQTL*. Rows are unique by *context*, *gene_id*, and *variant_id*. Lead cis rows come from *tensorQTL* `map_cis` with `qval < 0.05`, independent rows have `pval_perm < 0.05`. The sheet reports context, gene, variant and signal-class identifiers, DEG and GWAS support flags (boolean variables indicating yes/no membership in respective sets), distance and allele-frequency fields, nominal association statistics, permutation q-value support, and independent signal rank.

2) *nominal_BH05* sheet

This sheet contains one row per nominal eQTL gene-variant pair retained at Benjamini-Hochberg FDR < 0.05 within context. This sheet reports context, gene and variant identifiers, DEG and strict GWAS support flags, distance and allele-frequency fields, nominal p-value, FDR, slope, and slope standard error.

3) *eQTL_summary* sheet

This sheet contains one row per eQTL result type and context, summarizing sample counts, genes tested, eGenes, DEGs, GWAS-supported eGenes, combined DEG+GWAS categories, significant pairs, independent signals, nominal pairs, and source-support counts.

4) *intersecting_genes* sheet

This sheet lists non-empty gene intersections by eQTL result type, context, disorder, and annotation set, including gene counts and gene ID/gene symbol lists.

DEG columns annotate overlap with the broad DEG union and disorder-specific DEG sets for MDD and BD (MDD_DEG, BD_DEG). Exact variant support indicates that the eQTL variant is in the corresponding GWAS variant set; gene-list support indicates overlap with curated GWAS risk-gene lists and does not imply variant-level significance. Combined strict support columns are strict ($p<5\times10^{-8}$) GWAS exact-variant support or curated gene-list support. Exploratory ($p<1\times10^{-5}$) exact-variant support is provided for MDD and BD where available.

**Supplementary Table 7**. **Colocalization between domain$_{CT}$ eQTLs and psychiatric GWAS data.** This Excel workbook contains worksheets *coloc_pass* and *coloc_summary* with final colocalization results integrating domain$_{CT}$ eQTL summary statistics with MDD, BD, and SCZD GWAS summary statistics.

1) *coloc_pass* sheet

This sheet contains one row per final disorder-gene-domain$_{CT}$ colocalization locus with colocalization statistics computed with *coloc*. For each signal, the sheet reports disorder, context, gene identifiers, lead SNP and rsID (variant ID), number of SNPs evaluated, ABF posterior probabilities (`PP0`-`PP4`), derived posterior summaries `PP34` and `PP4_over_PP34`, lead-SNP posterior support metrics, 95% credible-set size, DEG annotations, exact eQTL lead-variant support (as reported by *tensorQTL*), exact GWAS lead-variant support, and gene-level GWAS support.

Posterior columns follow `coloc.abf` definitions: `PP3` is the posterior probability that both traits have association signals in the locus but with distinct causal variants; `PP4` is the posterior probability of one shared causal variant; `PP34` is `PP3 + PP4`; and `PP4_over_PP34` is `PP4/(PP3 + PP4)`, the conditional probability of colocalization among loci with two-trait association evidence. Exact eQTL support indicates that the colocalized lead SNP is also present as the same gene-variant pair in the final *cis_independent* or *nominal_BH05* eQTL sheet. Exact GWAS support indicates that the lead SNP is in the row disorder's GWAS variant set at the documented threshold. Gene-level GWAS support indicates overlap with curated GWAS risk-gene lists, without requiring the lead SNP to be GWAS-significant.

2) *coloc_summary* sheet

This sheet contains one row per disorder and context, reporting the number of genes tested, final colocalization loci, unique final colocalization genes, and the corresponding gene list.
